# Re-Engineering P(V) Chemical Warfare: Harnessing Stereogenic Phosphorus-Azoles for Protein Ligand Discovery *In Vivo*

**DOI:** 10.64898/2026.01.27.702106

**Authors:** R. Justin Grams, Olivia Murtagh, Madeleine Ware, Serhii Vasylevskyi, Ku-Lung Hsu

## Abstract

P(V) electrophiles such as tabun, sarin, soman, and VX are notorious for their lethality and nefarious intent in chemical warfare. Consequently, these deadly agents have largely been abandoned except for fluorophosphonate tool compounds that were repurposed for activity-based protein profiling (ABPP). Stereogenic P(V) centers hold strong potential as enabling scaffolds for synthetic and medicinal chemistry due to their inherent chirality and favorable bioavailability but are limited principally by potent off-target toxicity. Herein, we developed phosphorus-azole exchange (PhAzE) chemistry for tuning reactivity of the stereogenic P(V) pharmacophore to increase selectivity and mitigate off-target activity in cells and animal models. We demonstrate ultrapotent (300 pM in cells, 1 mg kg^-1^ in mice), enantioselective, covalent inhibition of the serine hydrolases DPP8/9 with PhAzE ligand in cells and *in vivo*; no overt toxicity was detected in mice treated daily over the course of a week. These finding show the P(V) electrophile can potently and enantioselectively engage a target protein without a deadly outcome, charting a path towards broader adoption of these agents in laboratory and industry settings.

## INTRODUCTION

Among the chemical warfare agents employed throughout modern history, the organophosphorus compounds VX, tabun, sarin, and soman are notorious for their potent lethality. As P(V) electrophiles, these molecules can react covalently with biological targets, most notably enzymes involved in neurotransmission (Figure 1A).^1^ Sarin was stockpiled in large quantities in Germany during World War II, though it was never used on the European battlefield. Efforts to understand its deadly mechanism revealed that sarin, like tabun and soman, irreversibly inhibits acetylcholinesterase, the enzyme responsible for breaking down the neurotransmitter acetylcholine.^2-3^ Acetylcholinesterase inhibition leads to uncontrolled accumulation of acetylcholine at neuromuscular junctions and synapses, resulting in muscle paralysis, convulsions, respiratory failure, and ultimately death. The widespread condemnation of nerve agents culminated in global banning of the development, production, stockpiling, and use of chemical weapons.^4-5^ This tragic legacy has largely stigmatized this compound class, precluding its further exploration for drug development despite potent *in vivo* activity.

**Figure 1.**
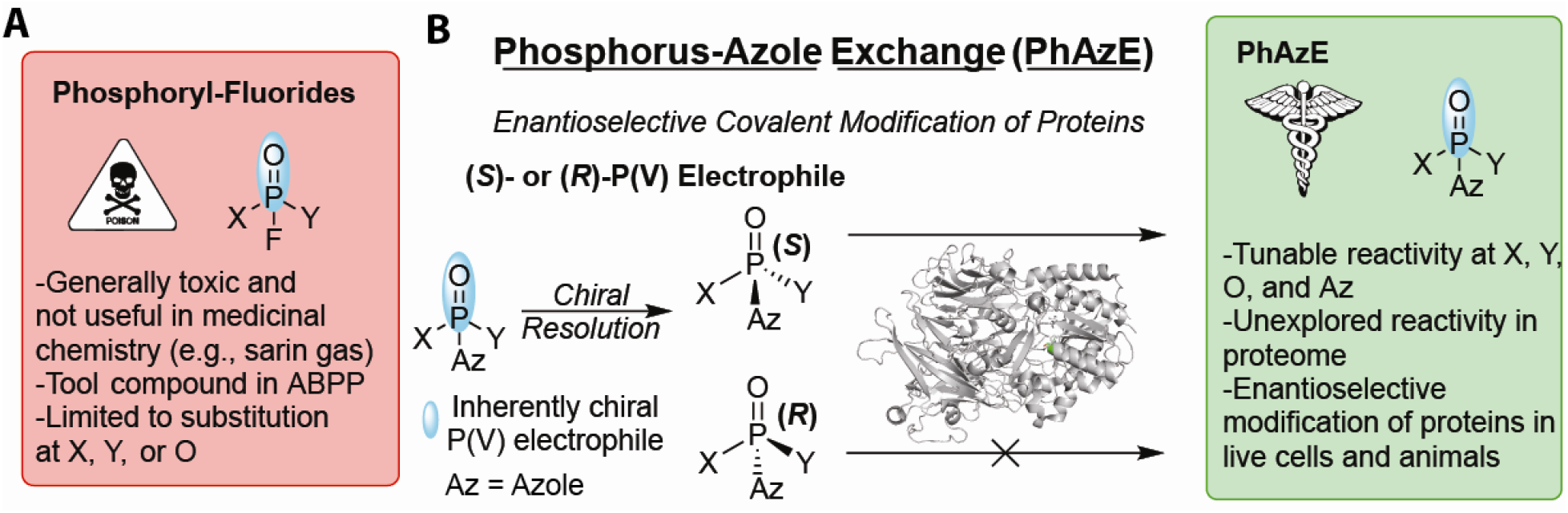
Development of PhAzE as a stereogenic P(V) electrophile for therapeutic investigation. Schematic of re-engineering phosphoryl-fluoride chemistry (**A**) to develop highly tunable enantioselective Phosphorus-Azole Exchange (PhAzE) chemistry (**B**).

While ill-suited for medicine because of toxicity, the P(V) electrophile has been successfully deployed for academic investigations. Fluorophosphonates covalently modify the catalytic serine in active sites of most, if not all, metabolic serine hydrolases that when coupled to proteomic readouts can facilitate activity-based protein profiling (ABPP, Figure S1A).^6-8^ This functional proteomic method has enabled basic discoveries in serine hydrolase biology, which led to inhibitor and drug development (Figure S1A).^9-19^ Moreover, the success of fluorophosphonates was foundational for exploring additional electrophiles for covalent targeting of reactive cysteines and less nucleophilic sites harboring, for example, a tyrosine, lysine or histidine (Figure S1A).^20-27^ The adoption of covalency into mainstream drug discovery campaigns had produced precision therapeutics such as ibrutinib, sotorasib, and others that exploit strategically located or mutated residues for selective and durable target protein engagement.^28 20, 29^ More recently, stereoselective labeling with electrophilic compounds offers a promising strategy to improve proteome-wide selectivity (Figure S1B).^24, 30-37^ However, because the electrophiles themselves are achiral, stereoselectivity relies on bulky, chiral substituents near the reactive site.^35-36, 38^ Rare examples like a chiral sulfonimidoyl fluoride demonstrate that chirality at the electrophilic center alone can drive enantioselective protein modification.^39,40^

P(V) electrophiles are uniquely positioned to enable enantioselective covalent modification of biomolecules, offering a direct strategy to interrogate and exploit chirality at the electrophilic center (Figure S1C).^41^ Only a few studies have explored stereoselective P(V)-based reactivity in complex proteomes. Notable examples include sulphostin, a chiral natural product that inhibits dipeptidyl peptidases 4, 8 and 9 (DPP4/8/9)^42-45^ and the antiviral pro-drug (*S*)-remdesivir, which is more potent than the (*R*)-stereoisomer, likely due to CES1-mediated stereoselective hydrolysis (Figure S1C).^46-48^ Yet, both molecules contain multiple stereocenters, limiting mechanistic understanding on whether the P(V) electrophile alone drives stereoselectivity. The use of P(V) chirality has been demonstrated in part through pioneering work in the development of chiral phosphorothioate “ψ-reagents” to synthesize antisense oligonucleotides (ASOs) with chiral backbones,^49-54^ nucleoside thioisosteres,^55^ and chemoselective phosphorylation reagents for alcohols^56^ and serine residues.^57^ These studies underscore the value of chiral P(V) centers in drug development. Phosphorothioate chemistry was recently extended to include tryptoline-based covalent ligands targeting various nucleophilic residues; however, these reagents were racemic at the P(V) center.^58^

Herein, we disclose the development of phosphorus-azole exchange (PhAzE) chemistry as a potent and safe stereogenic P(V) electrophile for enantioselective targeting of enzymes *in vivo*. The selection of an azole leaving group mitigated toxicity inherent to phosphoryl-fluorides while facilitating the design of picomolar inhibitors that enantioselectively engaged serine hydrolases DPP8/9 in the brain of treated mice. Importantly, daily dosing of the DPP8/9 PhAzE molecule over the course of a week did not produce overt toxicity or engagement with acetylcholinesterase as measured by quantitative chemoproteomics. PhAzE chemistry bridges the gap to therapeutics using stereogenic P(V) electrophiles and repositions this toxic pharmacophore for synthetic and therapeutic investigations.

## RESULTS

Initially, we tested whether enantiomers of fluorophosphonate (FP)-alkyne could be utilized for enantioselective ABPP. These studies would assess the extent to which a less sterically hindered stereogenic P(V) electrophile could facilitate enantioselective recognition of enzyme active sites. THP1 cells were treated with 20 µM (−)- or (+)-FP-alkyne followed by cell lysis, copper-catalyzed azide-alkyne cycloaddition (CuAAC)^59-63^ conjugation of biotin-azide, avidin enrichment, and tandem mass tag (TMT)-based quantitative ABPP (See Supporting Information).^64-65^ Despite detection of 40+ family members, only a single serine hydrolase displayed enantioselective binding as defined by a >50% difference in enrichment signals between (−) and (+) enantiomers ((+)/(−) ratio of >1 or <0.5; Figure 2B and S2A). Next, we assessed the effects of rigidifying the electrophile using a previously reported cyclic phosphoryl-fluoride strategy^66^ to incorporate recognition elements akin to ψ-reagents^49^. After purification of enantiomers, we treated THP1 cells with (−)- or (+)-RJG-2273 and observed substantially enhanced enantioselective ABPP labeling (9 or 4 serine hydrolases displaying enantioselective binding, respectively; Figure 2B and S2B).

**Figure 2.**
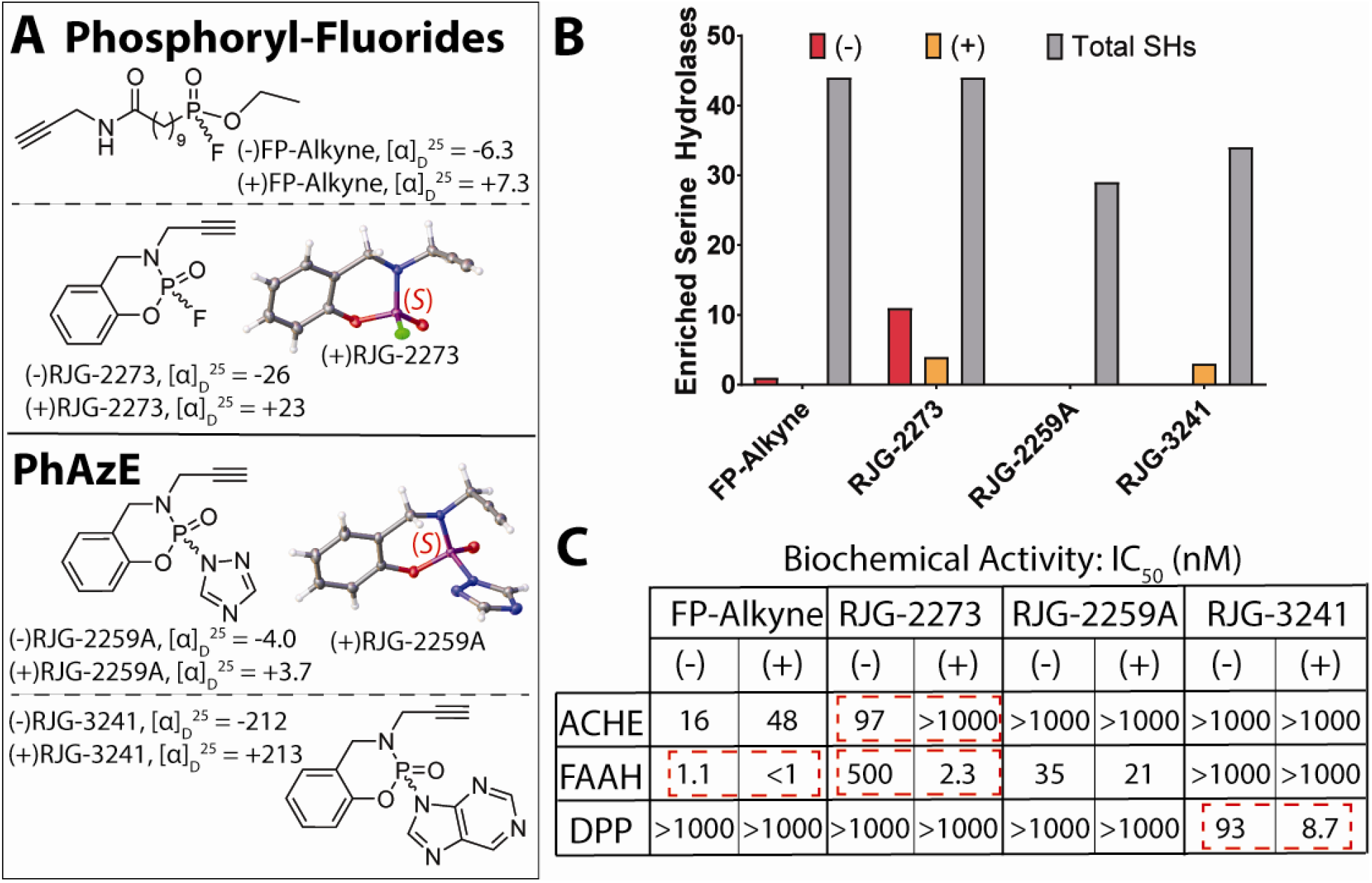
Enantioselective targeting of serine hydrolases with PhAzE. (**A**) Drawings of (−)- or (+)FP-Alkyne, RJG-2273, RJG-2259A, and RJG-3241 with respective specific rotation values or crystal structures. (**B**) Bar graph showing total and >50% enantioselectively enriched serine hydrolases from live THP1 monocytes treated with 20 µM (−)- or (+)-enantiomers. Data is representative of three independent biological replicates See Supporting Information Figure 2 for complete graph of enriched serine hydrolases. (**C**) Table of IC_50_ values of derived from biochemical assays. Live N2a cells were treated with respective enantiomers of compounds for 4 hours at 37 °C, then lysate was used to perform acetylcholinesterase (ACHE), fatty-acid amide hydrolase (FAAH), and dipeptidyl peptidase (DPP) biochemical assays. See Supporting Information for experimental details.

While the cyclic phosphoryl-fluoride strategy improved enantioselectivity, this probe retained the potent acetylcholinesterase (ACHE) inhibitory activity inherent to P(V) electrophiles (Figure 2C and S3E).^66-67^ We considered alternative leaving groups (LGs) as a means to temper the electrophile and mitigate covalent binding to ACHE. Azoles are effective and tunable nucleofuges to activate carbonyl^15-17^ and sulfonyl electrophiles for reactions at protein sites.^20, 24, 68-79^ Initially, we installed a 1,2,4-triazole in place of fluoride to produce RJG-2259A for mediating protein reactions via phosphorus-azole exchange (PhAzE) chemistry. We purified enantiomers and tested activity of the enantiomeric probes in THP1 monocytes and observed covalent binding to >20 serine hydrolases (Figure 2B), but negligible enantioselectivity (Figure S2C). Replacing the triazole with a purine and purifying the resulting enantiomers produced alkynyl probes (−)- and (+)RJG-3241, the latter of which enantioselectively enriched 3 serine hydrolases in THP1 cells (Figure 2B and S2D).

Next, we treated live Neuro-2A (N2a) mouse neuroblastoma cells with varying concentrations of (−)- or (+)-FP-Alkyne, -RJG-2273, -RJG-2259A, or -RJG-3241 to further assess enantioselective serine hydrolase binding and ACHE off-target activity in living cells. Importantly, PhAzE probes RJG-2259A and -3241 exhibited reduced inhibitory activity towards ACHE compared with phosphoryl-fluorides (Figure 2C and S3E). As expected, (−)- and (+)FP-Alkyne completely inhibited ACHE activity at 1 µM in live N2a cells (Figure 2C and S3E). Akin to the LC-MS/MS data, we found (−)RJG-2273 was a more potent inhibitor of ACHE biochemical activity compared to (+)RJG-2273, whereas both enantiomers of RJG-2259A and -3241 were weak inhibitors of ACHE (Figure 2B and S3E). In contrast, (+)RJG-2273 was a more potent inhibitor of FAAH biochemical activity, supporting the importance of P(V) chirality for recognition and covalent inactivation of serine hydrolases (Figure 2C and S3E).

We also observed subnanomolar, enantioselective inhibition of FAAH with (−)FP-Alkyne compared to the (+)FP-Alkyne (Figure 2C). We observed comparable inhibition of FAAH biochemical activity (Figure 2C and S3E) and fluorophosphonate-rhodamine (FP-Rh) labeling (Figure S3C) with (−)- and (+)RJG-2259A, suggesting RJG-2259A is a covalent inhibitor of serine hydrolases. In contrast, (+)RJG-3241 was a ∼10-fold more potent dipeptidyl peptidase (DPP) inhibitor compared to (−)RJG-3241, and we observed negligible inhibition with FP-Alkyne, RJG-2273, and RJG-2259A, thus highlighting the tunability of the PhAzE scaffold for enantioselective reactivity and specificity. Notably, several proteins were enantioselectively inhibited by (−)- or (+)FP-Alkyne and -RJG-2273 at lower concentrations as observed by gel-based competitive ABPP with FP-Rh (Figure S3A,B).

To initiate development of PhAzE ligands, we envisaged a structure-activity relationship (SAR) that would explore *N*-substitutions and azole LGs capable of potent inhibition of diverse serine hydrolase targets. The sp^3^-rich cyclopropyl moiety^80^ was installed to produced *N*-cyclopropyl or *N*-cyclopropylmethyl analogs RJG-3130-1 and -3045, respectively (Figure 3A). We treated live N2a cells with RJG-2273, RJG-3130-1, or RJG-3045 (1 µM compounds) followed by cell lysis, labeling of proteomes with FP-biotin, and TMT-based quantitative ABPP to assess inhibitory activity against the serine hydrolase family.^6^ FP-biotin labeled proteins were enriched by avidin chromatography, multiplexed using TMT, and quantified by liquid chromatography tandem mass spectrometry (LC-MS/MS, see Supporting Information for additional details).^64-65^ Phosphoryl-fluoride molecule RJG-2273 inhibited ACHE, whereas no inhibition was observed with PhAzE compounds RJG-3130-1 or RJG-3045 (Figure 3B). Both RJG-3130-1 or RJG-3045 showed comparable inhibitory activity against DPP8, APEH, and DPP9 (competition ratio or CR <0.5, 50% inactivation; Figure 3B and Figure S4). A dose response study in live N2a cells revealed RJG-3130-1 was about a 10-fold more potent inhibitor of DPPs and APEH compared to RJG-3045 (IC_50_ ≈ 8.5 nM for DPPs and APEH, Figure 3C, D and Figure S5).

**Figure 3.**
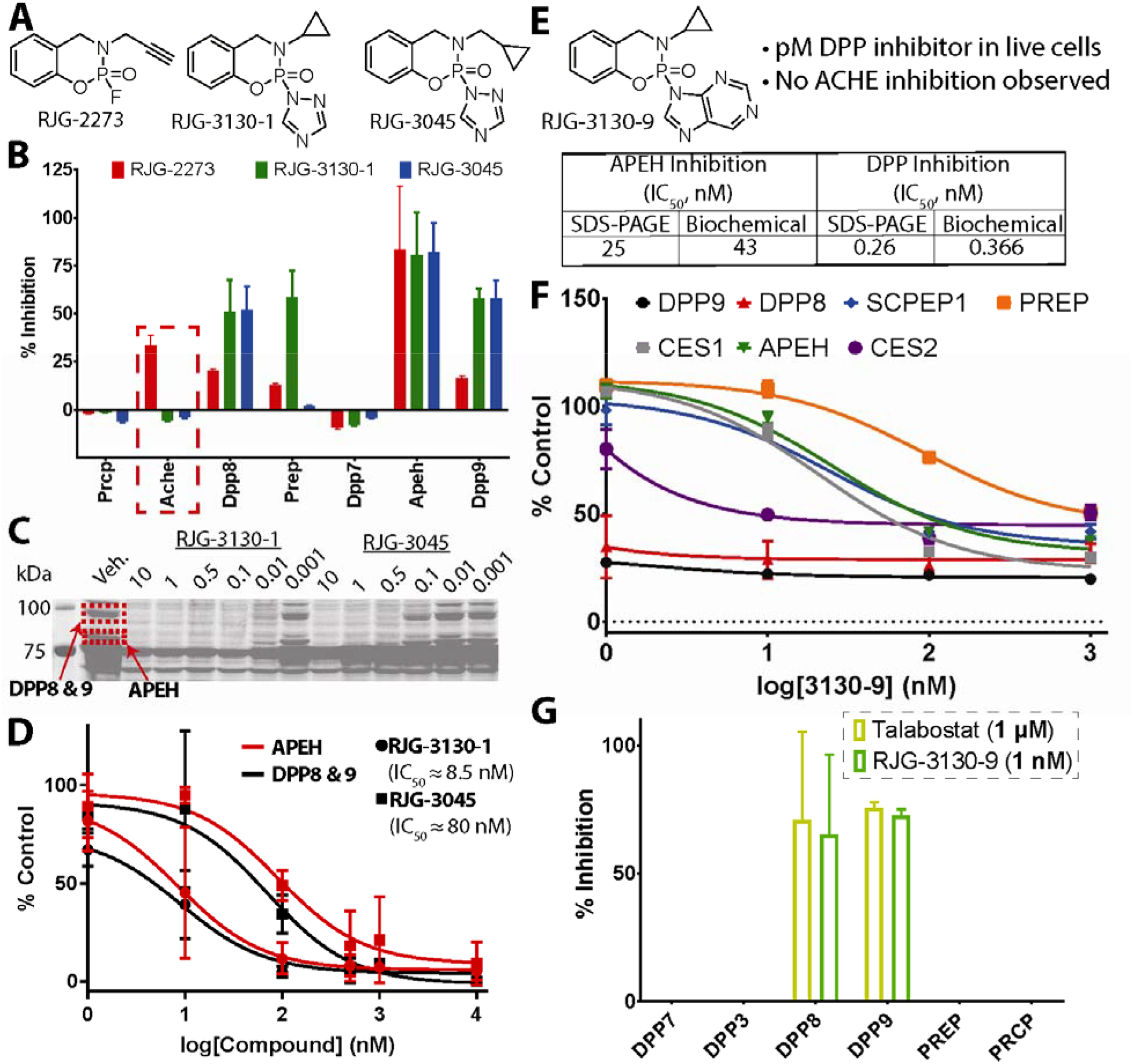
Developing PhAzE chiral ligands for ultrapotent enzyme inactivation. (**A**) Phosphoryl-fluoride molecule RJG-2273 and non-alkynyl ligands RJG-3130-1 and RJG-3045. (**B**) Graph of inhibited proteins quantified using Tandem Mass-Tag (TMT) LC-MS/MS. Live N2a cells were treated with 1µM RJG-2273, RJG-3130-1, or RJG-3045. The red dashed box highlights inhibition of acetylcholinesterase (ACHE) with RJG-2273, but not PhAzE ligands. Data is representative of three independent biological replicates. See Supporting Information Figure S4 for complete list of serine hydrolases. (**C**) SDS-PAGE based competition experiment using FP-Rh as a probe and a dose response of PhAzE ligands in live N2a cells. See Supporting Information Figure 5 for full length gel. (**D**) Graph of dose response curves derived from dividing raw fluorescence values of treatments over vehicle from Figure 3C. (**E**) Drawing of lead DPP inhibitor RJG-3130-9 and a table of IC_50_ values for APEH and DPPs obtained from dose response curves derived from dividing raw fluorescence values of treatments over vehicle and biochemical assays (See Supporting Information Figure 9 for full length gel and graphs). (**F**) Dose response curves of inhibited proteins derived from competition ratios using TMT abundances, *e*.*g*., competition ratio (3130-9/Vehicle), of proteins inhibited by RJG-3130-9 in live THP1 monocytes. (**G**) Graph comparing inhibition of DPP8 and 9, but not DPPs 2/7, 3, PREP, or PRCP, with 1 µM talabostat or 1 nM RJG-3130-9. % Inhibition was calculated using the competition ratio (CR) of (compound / vehicle). See Supporting Information Figure 9 for volcano plots.

RJG-3130-1 was selected for further development given its superior activity against serine peptidases. We synthesized several PhAzE molecules based on the RJG-3130 scaffold but further diversified by various azole LGs. PhAzE ligands were tested directly in live N2a cells to assess serine hydrolase inactivation as determined by FP-Rh-mediated, gel-based competitive ABPP and blockade of biochemical activity using substrate assays (1-1000 nM, 4 hours; Figure S6 and S7). We found RJG-3130-9 was a subnanomolar inhibitor of DPP probe labeling with apparent selectivity for DPPs over APEH compared to RJG-3130-8; neither compound showed activity towards ACHE (Figure S7). Compared with existing APEH (AA74-1^15^) and DPP8/9 (talabostat^81-83^) inhibitors, RJG-3130-9 was clearly a scaffold more optimized for DPP inactivation (Figure 3E and S7B). Note that two isomers of RJG-3130-9 were isolated after purification; the first isomer, presumed to be the *N*7-regioisomer, degraded quickly and was therefore not tested (Figure S8).

Widening the dose range revealed RJG-3130-9 is a picomolar inhibitor (∼300 pM) of the ∼100 kDa band matching the molecular weight of DPP8/9 in live cells (Figure 3E and S9A,B). Considering the existence of at least seven known DPPs^83-84^, we next performed TMT-based competitive ABPP to assess DPP and serine hydrolase family-wide selectivity of compounds. Treatment of N2a cells with RJG-3130-9 resulted in significant blockade of DPP8 and 9 probe labeling across all concentrations tested (1 – 1000 nM, Figure 3F and S10B-E). Off-target activity (e.g., APEH, ACOT2, CES) of RJG-3130-9 was only observed at higher concentrations and absent at lower concentrations that retained potent DPP8/9 inactivation (>50% blockade at 1 nM RJG-3130-9, Figure S10E). Notably, RJG-3130-9 was able to achieve a similar degree of DPP8 and 9 inhibition at a 1000X lower concentration compared to talabostat in live cells (Figure 3G).

Next, we investigated chiral purification to determine whether PhAzE ligands display enantioselective binding activity. We initiated efforts with chiral purification of RJG-3130-1 to test if the RJG-3130 scaffold was advantageous compared to alkynyl probe RJG-2259A (Figure 2). Although enantiomeric excess (e.e.) was >98% after chiral purification, (−)- and (+)RJG-3130-1 began to racemize during concentration to dryness (Figure 4A and S11A,B). We attempted chiral purification of RJG-3130-3 containing an imidazole LG that is more basic and presumably more stable because of reduced electrophilicity (Figure S6B); however, almost complete racemization occurred upon concentration (Figure 4A and S11C,D). RJG-3130-9, which is modified with a bulkier purine LG, was readily purified by chiral supercritical fluid chromatography (SFC) and retained e.e. after concentrating to dryness (Figure 4A and S11E,F). We solved the crystal structure of (+)RJG-3130-9 by microcrystal electron diffraction (microED) and observed (*S*)-configuration at the P(V) electrophile and the *N*9-substituted regioisomer (Figure 4B). In addition, the *N*20 nitrogen is directed at the benzene ring at an angle of 124°, and *N*20 is ∼2.8 Å from the bond between carbons C5-C6 and ∼3.6 Å from C5, which is within range of a *n* – π* interaction and is near the summed van der Waals radii of 3.44 Å between N and C (Figure 4B).^85-86^

**Figure 4.**
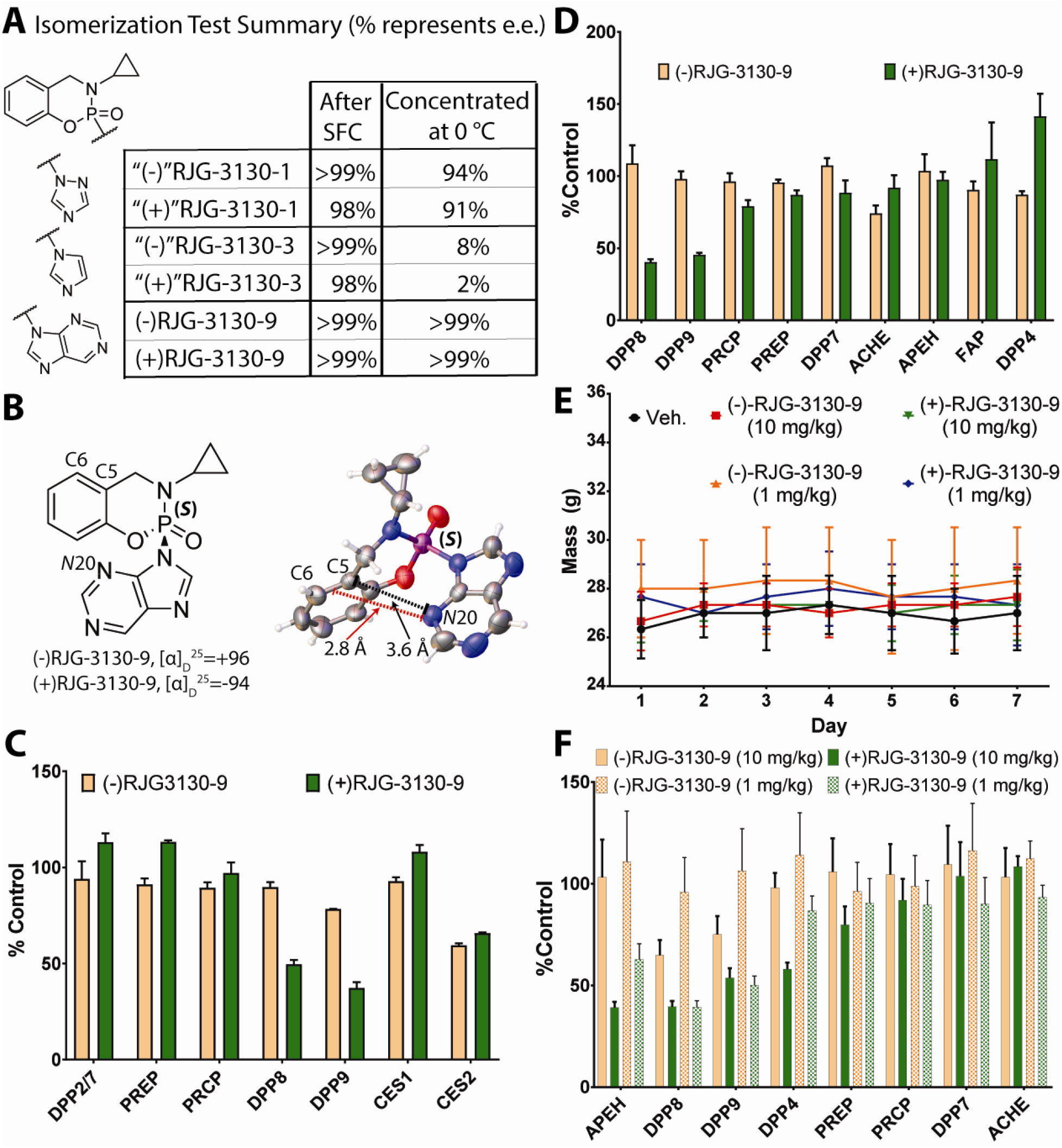
Enantioselective targeting of proteins *in vivo* with brain-penetrant PhAzE ligands. (**A**) RJG-3130-1, -3, and -9 were synthesized as racemates and respective enantiomers purified by supercritical fluid chromatography. Enantiomers of RJG-3130-1 and -3 isomerized upon concentration, whereas RJG-3130-9 retained enantiomeric excess (e.e.). (**B**) Drawing of (*S*)-(+)RJG-3130-9 and a crystal structure characterized by microED. The distance between *N*4 and C5 is 3.59 Å (black dashed line) and *N*4 and the bond between C5 and C6 is 2.89 Å (red dashed line). (**C**) Bar graph representing DPPs identified in a 16-plex TMT LC-MS/MS experiment using samples from live THP1 monocytes treated with vehicle or 1 nM (−)- or (+)RJG-3130-9. See Supporting Information Figure 13 for full dose response graphs. Data is representative of three independent biological replicates. (**D**) Bar graph of serine hydrolases enriched from brain tissue of mice treated with vehicle or 1 mg/kg of (−)- or (+)RJG-3130-9 for 4 hours. Samples were combined with TMT 6-plex and analyzed by LC-MS/MS. See Supporting Information Figure 14 for complete graph of enriched serine hydrolases. Data is representative of three independent biological replicates. (**E**) Average weights of mice over a seven-day treatment with vehicle, 10 or 1 mg/kg (−)RJG-3130-9, or 10 or 1 mg/kg (+)RJG-3130-9. See Supporting Information Figure 15 for additional graphs. (**F**) Bar graph of serine hydrolases enriched from brain tissue of mice treated with vehicle, 10 mg/kg, or 1 mg/kg of (−)- or (+)RJG-3130-9 for 7 days. Samples were combined with TMT 16-plex and analyzed by LC-MS/MS. Data is representative of three independent biological replicates. See Supporting Information Figure 16 for complete graph of enriched serine hydrolases.

We analyzed inhibition of DPPs with (−)- or (+)RJG-3130-9 in THP1 soluble lysate initially by gel-based competitive ABPP. (+)RJG-3130-9 displayed ultrapotent, time-dependent blockade of DPP probe labeling, supporting an irreversible covalent inactivation mechanism (Figure S12A, B). We compared inhibitory activity of (−)- and (+)RJG-3130-9 at 1 nM and found initial evidence for enantioselectivity at all time points tested (Figure S12C). Next, we evaluated whether the enantioselective blockade of probe labeling translated into DPP biochemical inactivation. (+)RJG-3130-9 displayed dose and time-dependent inhibition of DPPs achieving an estimated potency value (IC_50_) of 160 pM (4-hour treatment). (−)RJG-3130-9, in contrast, exhibited moderately potent inhibition with a ∼7-15-fold reduction in potency across the different treatment times tested (Figure S12D-F).

Next, we treated THP1 cells with (−)- or (+)RJG-3130-9 at varying concentrations to test whether the observed enantioselective activity against DPPs *in vitro* was recapitulated in live cells. Akin to *in vitro* findings, we detected >50% inhibition of DPP8/9 with (+)RJG-3130-9 compared to <25% inhibition with (−)RJG-3130-9 at 1 nM as measured by TMT-based competitive ABPP (Figure 4C and S13D, E). Additional DPPs identified were not significantly inhibited at 1 nM (Figure 4C). At higher concentrations of (−)- and (+)RJG-3130-9, we observed PREP, CES1, and CES2 off-target activity, which was not enantioselective and largely eliminated at lower concentrations where (+)RJG-3130-9 retained potent DPP8/9 blockade (Figure 4C, S13C,F,G).

Given the promising cellular activity of PhAzE chiral ligands, we pursued assessment of *in vivo* activity with (−) and (+)RJG-3130-9. To the best of our knowledge, enantioselective covalent ligands published to date and stereogenic P(V) electrophiles – excluding nefarious intent with fluorophosphonates – have not been tested for enantioselective covalent modification of proteins *in vivo*. Mice were treated with vehicle, (−)-, or (+)RJG-3130-9 for 4 hours (1 mg/kg) or 7 days (10 or 1 mg/kg). Tissue was harvested and samples were multiplexed for quantitation of protein inhibition after 4 hours or 7 days of treatment *via* 6- or 16-plex TMT-based competitive ABPP, respectively.

Strikingly, we discovered only DPP8/9 were significantly inhibited by >50% in the brains of (+)RJG-3130-9-treated mice, whereas negligible inhibition was observed with (−)RJG-3130-9 across the 48 serine hydrolases quantified (Figure 4D and S14). These findings support the ability of PhAzE ligands to cross the blood-brain barrier and enantioselectively engage target proteins. Importantly, minimal-to-no ACHE inhibition was observed after acute (4 hours) or chronic dosing consisting of a 7-day treatment regime with 1 or 10 mg/kg (−)- or (+)RJG-3130-9 (Figure 4D,F). We did not observe changes in body weight of mice treated with RJG-3130-9 enantiomers, providing initial evidence of a promising safety profile for PhAzE *in vivo* (Figure 4E and S15). Finally, we compared the competition binding profile of 10 and 1 mg/kg (−)- or (+)RJG-3130-9 after 7 days of treatment. At the higher dose (10 mg/kg), we detected only APEH and BCHE as additional competed targets of (+)RJG-3130-9 (Figure 4F and S16A). At lower concentrations (1 mg/kg), DPP8/9 remained the only significantly and enantioselectively competed targets of (+)RJG-3130-9 (>50% competition, Figure 4F and S16B).

## DISCUSSION

Despite their utility in chemical biology, fluorophosphonates and related P(V) electrophiles remain challenging to deploy more broadly due to their inherent toxicity and origins as lethal nerve agents, leaving no clear path for translational applications. If toxicity could be mitigated, stereogenic P(V) hubs would serve as enabling starting points for synthetic organic chemistry and drug discovery because of their potent bioavailability and inherent chirality. The latter feature is highly desirable for drug development^30-31, 49^ but often relies on bulky chiral substitutions adjacent or distal to achiral electrophiles,^34-39, 58, 87^ with only a few examples of chirality at the electrophile.^40^ Here, we developed a stereogenic P(V) electrophile dubbed PhAzE for safe and enantioselective protein inactivation *in vivo*, repositioning these agents from deadly poisons to valuable leads for therapeutic discovery.

We discovered that chiral phosphoryl-fluorides including FP-based agents could achieve enantioselective targeting of serine hydrolases but largely retained potent ACHE binding activity (Figure 2). The switch from an acyclic^6^ to cyclic P(V) electrophile^66^ improved enantioselectivity and reduced ACHE inactivation (Figure 2C and S3E). These promising findings along with the relative ease for accessing the tunable and constrained stereogenic P(V) electrophile in RJG-2273 prompted development of PhAzE analogs bearing azole LGs. Azoles are effective LGs for tunable activation of electrophilic protein modification^17-20, 24, 68, 88-93^ providing a testable path to reduce toxicity of the P(V) electrophile by selectivity optimization. We tested this hypothesis by synthesizing stereogenic PhAzE probes and discovered that the purine azole mitigated ACHE off-target activity while retaining sufficient enantioselective binding activity for downstream ligand discovery efforts (Figure 2C, S2, and S3).

The initial PhAzE design using a triazole LG ((−)- and (+)RJG-2259A) showed cellular activity but low enantioselectivity, prompting exploration of additional azoles. The natural metabolite purine emerged as an effective LG for enantioselective protein modification. We speculate that the bulkier nature of purines compared with the triazole could be contributing to improved enantioselectivity; future studies expanding on the SAR of PhAzE will further support this hypothesis. Nonetheless, we were encouraged to see a significant improvement in enantioselectively enriched serine hydrolases and differential, potent inhibition of DPP biochemical activity with RJG-3241 that was not observed with the phosphoryl-fluoride probes FP-Alkyne or RJG-2273 (Figure 2C and S3E).

Given the enantioselective activity of RJG-3241, we transitioned from probe to ligand optimization to assess whether PhAzE could achieve enantioselective inactivation across the serine hydrolase superfamily. Our SAR campaign characterized RJG-3130-9 as a picomolar DPP8/9 inhibitor with good selectivity over APEH and no inhibition of ACHE in live cells (Figure 3E, S7 and S9). Comparison of RJG-3130-9 with the existing DPP8/9 inhibitor talabostat revealed the PhAzE ligand could achieve similar target engagement at ∼1,000-fold lower concentrations than this clinical candidate (Figure 3G).

Although chiral purification of phosphoryl-fluorides was straightforward, we had to overcome several challenges with chiral purification of PhAzE. Partial or complete racemization of RJG-3130-1 (1,2,4-triazole LG) or RJG-3130-3 (imidazole LG) was observed upon concentration of fractions after chiral purification, respectively, a surprising result given the relative ease in purifying enantiomers of PhAzE probe RJG-2259A (Figure 2). Ultimately, we observed that (−) and (+)RJG-3130-9 retained >99% e.e. after concentration; these chiral ligands exhibited time-dependent, enantioselective inhibition of DPP8/9 in THP1 monocytes and soluble lysate (Figure 4C and S12, respectively). Crystallography revealed (+)RJG-3130-9 exists in a (*S*)-configuration, the purine is substituted at *N*9, and this regioisomer is stabilized by an intramolecular *n* – π interaction (Figure 4B).^86^ Similar to enantioselective enrichment of serine hydrolases with (+)RJG-3241, (+)RJG-3130-9 enantioselectively inhibited DPP8 and 9 by >50% compared to negligible inhibition with (−)RJG-3130-9 at the same concentration (1 nM compounds, Figure 4C and S13).

Finally, we demonstrated covalent and enantioselective inhibition *in vivo* with a chiral P(V) electrophile. The DPP8/9 selectivity and enantioselectivity of (+)RJG-3130-9 detected in cells was recapitulated in brain proteomes from treated mice. Specifically, (+)RJG-3130-9 (1 mg/kg, intraperitoneal or i.p.) inhibited FP-biotin probe labeling of DPP8/9 by >50% with minimal to no inhibition of the >40 serine hydrolases detected in brain. Notably, ACHE was not inhibited in mice treated under acute (4 hours) and more chronic exposure (7 days) at 1 or 10 mg/kg i.p. treatment (Figure 4D,F, S14, and S16). We did not detect overt changes in the weight of mice in any PhAzE treatment group (Figure 4E and S15). The lack of ACHE inhibitory activity combined with stable body weight under prolonged treatment provides early but critical evidence of *in vivo* safety for the PhAzE compound class. These findings represent, to the best of our knowledge, the first case of enantioselective inhibition of proteins *in vivo* with a P(V) electrophile that was not deadly.

By extension, the improved safety profile of PhAzE chemistry paves the way for designing stereogenic PhAzE molecules that are readily available for synthetic organic chemistry. The availability of phosphoryl-chlorides is lacking compared to sulfonyl-chloride congeners used for various synthetic chemistry applications including the synthesis of sulfonyl-fluorides that are now widely used in synthetic and medicinal chemistry.^94^ Phosphorus-fluoride exchange (PFEx) click chemistry was recently published as a versatile and efficient P(V) electrophilic hub for synthesizing phosphoryl moieties.^66^ Synthesis of PFEx molecules requires precautions such as harsh quenching conditions and plasticware when using fluorides that are toxic and known to etch/damage glassware, respectively.^95^ Similar to results we presented above, PFEx molecules are potent inhibitors of ACHE activity that require scientists to take precautions to prevent accidental exposure,^66^ thus limiting their potential for commercial availability and general accessibility. We envision that PhAzE chemistry presented herein has the potential to be translated to synthetic organic chemistry applications with minimal-to-no additional requirements to achieve chemical transformations similar to PFEx.^66^

## CONCLUSIONS

In summary, we re-engineered fluorophosphonates by substituting the fluoride for azole leaving groups to produce potent, enantioselective, and *in vivo*-active PhAzE ligands for covalent ligand and potential therapeutic discovery. We optimized PhAzE to achieve picomolar and enantioselective inhibition of serine hydrolases in live cells. Moreover, (*S*)-(+)RJG-3130-9 enantioselectively inhibited DPP8/9 *in vivo*, marking these findings as, to the best of our knowledge, the first disclosure of enantioselective and non-deadly covalent inhibition *in vivo* with a P(V) electrophile.

## Acknowledgements

We thank all members of the Hsu lab for review of this manuscript. We thank Peimin Fan and Yonghong Li (Pharmaron Inc.) for small molecule synthesis, chiral purification, and microED.

## Contributions

R.J.G. and K.-L.H conceived the study. R.J.G. synthesized and characterized all PhAzE compounds, as well as RJG-2273, and supervised synthesis and characterization of FP-biotin and chiral compounds. R.J.G. performed all experiments including tissue culture and cellular treatments, live animal treatments and tissue harvesting, biochemical assays, gel-ABPP, LC-MS/MS ABPP to generate proteomic data, and analysis of proteomic data. M.L.W. and O.L.M. designed the biotin enrichment protocol and LC-MS/MS methods. S.V. performed X-ray crystallography. K.-L.H. supervised the study. All authors edited and approved the paper.

## Competing Interests

K.-L.H. is a founder and scientific advisory board member of Hyku Biosciences. A patent application has been filed on the work presented in this manuscript.

## References

(1) Labaška, M.; Gál, M.; Mackuľak, T.; et al. Neutralizing the threat: A comprehensive review of chemical warfare agent decontamination strategies. J. Environ. Chem. Eng. 2024, 12 (6), 114243.

(2) Everts, S. The Nazi origins of deadly nerve gases. C&EN Global Enterprise 2016, 94 (41), 26–28.

(3) Voros, C.; Dias, J.; Timperley, C. M.; et al. The risk associated with organophosphorus nerve agents: from their discovery to their unavoidable threat, current medical countermeasures and perspectives. Chem.-Biol. Interact. 2024, 395, 110973.

(4) History, Looking back helps us look forward, OPCW. https://www.opcw.org/about-us/history x(accessed 12th September 2025).

(5) Convention on the Prohibition of the Development, Production, Stockpiling and Use of Chemical Weapons and on their Destruction, OPCW (n.d.). https://www.opcw.org/chemical-weapons-convention x(accessed.

(6) Liu, Y.; Patricelli, M. P.; Cravatt, B. F. Activity-based protein profiling: The serine hydrolases. Proc. Natl. Acad. Sci. 1999, 96 (26), 14694–14699.

(7) Long, J. Z.; Cravatt, B. F. The Metabolic Serine Hydrolases and Their Functions in Mammalian Physiology and Disease. Chem. Rev. 2011, 111 (10), 6022–6063.

(8) Bachovchin, D. A.; Koblan, L. W.; Wu, W.; et al. A high-throughput, multiplexed assay for superfamily-wide profiling of enzyme activity. Nat. Chem. Biol. 2014, 10 (8), 656–663.

(9) Egmond, N. v.; Straub, V. M.; Stelt, M. v. d. Targeting Endocannabinoid Signaling: FAAH and MAG Lipase Inhibitors. Annu. Rev. Pharmacol. Toxicol. 2021, 61 (1), 441–463.

(10) Long, J. Z.; Li, W.; Booker, L.; et al. Selective blockade of 2-arachidonoylglycerol hydrolysis produces cannabinoid behavioral effects. Nat. Chem. Biol. 2009, 5 (1), 37–44.

(11) Blankman, J. L.; Cravatt, B. F. Chemical Probes of Endocannabinoid Metabolism. Pharmacol. Rev. 2013, 65 (2), 849–871.

(12) Niphakis, M. J.; Cognetta, A. B.; Chang, J. W.; et al. Evaluation of NHS Carbamates as a Potent and Selective Class of Endocannabinoid Hydrolase Inhibitors. ACS Chem. Neurosci. 2013, 4 (9), 1322–1332.

(13) Chang, Jae W.; Niphakis Micah J.; Lum Kenneth M.; et al. Highly Selective Inhibitors of Monoacylglycerol Lipase Bearing a Reactive Group that Is Bioisosteric with Endocannabinoid Substrates. Chem. Biol. 2012, 19 (5), 579–588.

(14) Cisar, J. S.; Weber, O. D.; Clapper, J. R.; et al. Identification of ABX-1431, a Selective Inhibitor of Monoacylglycerol Lipase and Clinical Candidate for Treatment of Neurological Disorders. J. Med. Chem. 2018, 61 (20), 9062–9084.

(15) Adibekian, A.; Martin, B. R.; Wang, C.; et al. Click-generated triazole ureas as ultrapotent in vivo– active serine hydrolase inhibitors. Nat. Chem. Biol. 2011, 7 (7), 469–478.

(16) Adibekian, A.; Martin, B. R.; Chang, J. W.; et al. Confirming Target Engagement for Reversible Inhibitors in Vivo by Kinetically Tuned Activity-Based Probes. J. Am. Chem. Soc. 2012, 134 (25), 10345–10348.

(17) Hsu, K.-L.; Tsuboi, K.; Adibekian, A.; et al. DAGLβ inhibition perturbs a lipid network involved in macrophage inflammatory responses. Nat. Chem. Biol. 2012, 8 (12), 999–1007.

(18) Hsu, K.-L.; Tsuboi, K.; Whitby, L. R.; et al. Development and Optimization of Piperidyl-1,2,3-Triazole Ureas as Selective Chemical Probes of Endocannabinoid Biosynthesis. J. Med. Chem. 2013, 56 (21), 8257–8269.

(19) Hsu, K.-L.; Tsuboi, K.; Chang, J. W.; et al. Discovery and Optimization of Piperidyl-1,2,3-Triazole Ureas as Potent, Selective, and in Vivo-Active Inhibitors of α/β-Hydrolase Domain Containing 6 (ABHD6). J. Med. Chem. 2013, 56 (21), 8270–8279.

(20) Grams, R. J.; Hsu, K.-L. Reactive chemistry for covalent probe and therapeutic development. Trends Pharmacol. Sci. 2022.

(21) Narayanan, A.; Jones, L. H. Sulfonyl fluorides as privileged warheads in chemical biology. Chem. Sci. 2015, 6 (5), 2650-2659, 10.1039/C5SC00408J.

(22) Jones, L. H. Advances in sulfonyl exchange chemical biology: expanding druggable target space. Chem. Sci. 2025, 16 (23), 10119-10140, 10.1039/D5SC02647D.

(23) Boike, L.; Henning, N. J.; Nomura, D. K. Advances in covalent drug discovery. Nat. Rev. Drug Discov. 2022, 21 (12), 881–898.

(24) Kim, G.; Grams, R. J.; Hsu, K.-L. Advancing Covalent Ligand and Drug Discovery beyond Cysteine. Chem. Rev. 2025, 125 (14), 6653–6684.

(25) Jones, L. H. Chapter Four - Design of next-generation covalent inhibitors: Targeting residues beyond cysteine. In Annu. Rep. Med. Chem., Ward, R. A., Grimster, N. P. Eds.; Vol. 56; Academic Press, 2021; pp 95–134.

(26) Gehringer, M.; Laufer, S. A. Emerging and Re-Emerging Warheads for Targeted Covalent Inhibitors: Applications in Medicinal Chemistry and Chemical Biology. J. Med. Chem. 2019, 62 (12), 5673–5724.

(27) Niphakis, M. J.; Cravatt, B. F. Ligand discovery by activity-based protein profiling. Cell Chem. Biol. 2024, 31 (9), 1636–1651.

(28) Canon, J.; Rex, K.; Saiki, A. Y.; et al. The clinical KRAS(G12C) inhibitor AMG 510 drives anti-tumour immunity. Nature 2019, 575 (7781), 217–223.

(29) Backus, K. M.; Correia, B. E.; Lum, K. M.; et al. Proteome-wide covalent ligand discovery in native biological systems. Nature 2016, 534 (7608), 570–574.

(30) Agranat, I.; Caner, H.; Caldwell, J. Putting chirality to work: the strategy of chiral switches. Nat. Rev. Drug Discov. 2002, 1 (10), 753–768.

(31) Nguyen, L. A.; He, H.; Pham-Huy, C. Chiral drugs: an overview. Int. J. Biomed. Sci.. 2006, 2, 85–100.

(32) Sui, J.; Zhang, J.; Ching, C. B.; Chen, W. N. Expanding proteomics into the analysis of chiral drugs. Mol. BioSyst. 2009, 5 (6), 603-608, 10.1039/B903858B.

(33) Sanna, M. G.; Wang, S.-K.; Gonzalez-Cabrera, P. J.; et al. Enhancement of capillary leakage and restoration of lymphocyte egress by a chiral S1P1 antagonist in vivo. Nat. Chem. Biol. 2006, 2 (8), 434–441.

(34) Rosen, H. T.; Li, K.; Stieger, C. E.; et al. Sulfinyl Aziridines as Stereoselective Covalent Destabilizing Degraders of the Oncogenic Transcription Factor MYC. Angew. Chem. Int. Ed. 2025, 64 (48), e202508518.

(35) Zhang, Y.; Liu, Z.; Hirschi, M.; et al. An allosteric cyclin E-CDK2 site mapped by paralog hopping with covalent probes. Nat. Chem. Biol. 2025, 21 (3), 420–431.

(36) Njomen, E.; Hayward, R. E.; DeMeester, K. E.; et al. Multi-tiered chemical proteomic maps of tryptoline acrylamide–protein interactions in cancer cells. Nat. Chem. 2024, 16 (10), 1592–1604.

(37) Wang, Y.; Dix, M. M.; Bianco, G.; et al. Expedited mapping of the ligandable proteome using fully functionalized enantiomeric probe pairs. Nat. Chem. 2019, 11 (12), 1113–1123.

(38) Liu, Z.; Remsberg, J. R.; Li, H.; et al. Proteomic Ligandability Maps of Spirocycle Acrylamide Stereoprobes Identify Covalent ERCC3 Degraders. J. Am. Chem. Soc. 2024, 146 (15), 10393–10406.

(39) Chen, Y.; Craven, G. B.; Kamber, R. A.; et al. Direct mapping of ligandable tyrosines and lysines in cells with chiral sulfonyl fluoride probes. Nat. Chem. 2023, 15 (11), 1616–1625.

(40) Huang, H.-s.; Yuan, Y.; Wang, W.; et al. Enantioselective Synthesis of Chiral Sulfonimidoyl Fluorides Facilitates Stereospecific SuFEx Click Chemistry. Angew. Chem. Int. Ed. 2025, 64 (4), e202415873.

(41) Knouse, K. W.; Flood, D. T.; Vantourout, J. C.; et al. Nature Chose Phosphates and Chemists Should Too: How Emerging P(V) Methods Can Augment Existing Strategies. ACS Cent. Sci. 2021, 7 (9), 1473–1485.

(42) Sewald, L.; Tabak, W. W. A.; Fehr, L.; et al. Sulphostin-inspired N-phosphonopiperidones as selective covalent DPP8 and DPP9 inhibitors. Nat. Commun. 2025, 16 (1), 3208.

(43) Akiyama, T.; Abe, M.; Harada, S.; et al. Sulphostin, a Potent Inhibitor for Dipeptidyl Peptidase IV from Streptomyces sp. MK251-43F3. J. Antibiot. 2001, 54, 744–746.

(44) Abe, M.; Akiyama, T.; Umezawa, Y.; et al. Synthesis and biological activity of sulphostin analogues, novel dipeptidyl peptidase IV inhibitors. Bioorg. Med. Chem. 2005, 13 (3), 785–797.

(45) Abe, M.; Akiyama, T.; Nakamura, H.; et al. First Synthesis and Determination of the Absolute Configuration of Sulphostin, a Novel Inhibitor of Dipeptidyl Peptidase IV. J. Nat. Prod. 2004, 67 (6), 999–1004.

(46) Kokic, G.; Hillen, H. S.; Tegunov, D.; et al. Mechanism of SARS-CoV-2 polymerase stalling by remdesivir. Nat. Commun. 2021, 12 (1), 279.

(47) Warren, T. K.; Jordan, R.; Lo, M. K.; et al. Therapeutic efficacy of the small molecule GS-5734 against Ebola virus in rhesus monkeys. Nature 2016, 531 (7594), 381–385.

(48) Hu, H.; Mady Traore, M. D.; Li, R.; et al. Optimization of the Prodrug Moiety of Remdesivir to Improve Lung Exposure/Selectivity and Enhance Anti-SARS-CoV-2 Activity. J. Med. Chem. 2022, 65 (18), 12044–12054.

(49) Knouse, K. W.; deGruyter, J. N.; Schmidt, M. A.; et al. Unlocking P(V): Reagents for chiral phosphorothioate synthesis. Science 2018, 361 (6408), 1234–1238.

(50) Nassir, M.; Gherardi, L.; Redman, R. L.; et al. An Improved P(V) Thio-Oligonucleotide Synthesis Platform. Org. Lett. 2025, 27 (1), 97–102.

(51) Obexer, R.; Nassir, M.; Moody, E. R.; et al. Modern approaches to therapeutic oligonucleotide manufacturing. Science 2024, 384 (6692), eadl4015.

(52) Huang, Y.; Knouse, K. W.; Qiu, S.; et al. A P(V) platform for oligonucleotide synthesis. Science 2021, 373 (6560), 1265–1270.

(53) Flood, D. T.; Knouse, K. W.; Vantourout, J. C.; et al. Synthetic Elaboration of Native DNA by RASS (SENDR). ACS Cent. Sci. 2020, 6 (10), 1789–1799.

(54) Xu, D.; Rivas-Bascón, N.; Padial, N. M.; et al. Enantiodivergent Formation of C–P Bonds: Synthesis of P-Chiral Phosphines and Methylphosphonate Oligonucleotides. J. Am. Chem. Soc. 2020, 142 (12), 5785–5792.

(55) Zhang, H.-J.; Ociepa, M.; Nassir, M.; et al. Stereocontrolled access to thioisosteres of nucleoside di- and triphosphates. Nat. Chem. 2024, 16 (2), 249–258.

(56) Ociepa, M.; Knouse, K. W.; He, D.; et al. Mild and Chemoselective Phosphorylation of Alcohols Using a Ψ-Reagent. Org. Lett. 2021, 23 (24), 9337–9342.

(57) Vantourout, J. C.; Adusumalli, S. R.; Knouse, K. W.; et al. Serine-Selective Bioconjugation. J. Am. Chem. Soc. 2020, 142 (41), 17236–17242.

(58) Sharma, H. A.; Bielecki, M.; Holm, M. A.; et al. Proteomic Ligandability Maps of Phosphorus(V) Stereoprobes Identify Covalent TLCD1 Inhibitors. J. Am. Chem. Soc. 2025, 147 (18), 15554–15566.

(59) Rostovtsev, V. V.; Green, L. G.; Fokin, V. V.; Sharpless, K. B. A Stepwise Huisgen Cycloaddition Process: Copper(I)-Catalyzed Regioselective “Ligation” of Azides and Terminal Alkynes. Angew. Chem. Int. Ed. 2002, 41 (14), 2596–2599.

(60) Meldal, M.; Tornøe, C. W. Cu-Catalyzed Azide–Alkyne Cycloaddition. Chem. Rev. 2008, 108 (8), 2952–3015.

(61) Speers, A. E.; Adam, G. C.; Cravatt, B. F. Activity-Based Protein Profiling in Vivo Using a Copper(I)-Catalyzed Azide-Alkyne [3 + 2] Cycloaddition. J. Am. Chem. Soc. 2003, 125 (16), 4686–4687.

(62) Speers, A. E.; Cravatt, B. F. Profiling Enzyme Activities In Vivo Using Click Chemistry Methods. Chemistry & Biology 2004, 11 (4), 535–546.

(63) Speers, A. E.; Cravatt, B. F. Chemical Strategies for Activity-Based Proteomics. ChemBioChem 2004, 5 (1), 41–47.

(64) Rauniyar, N.; Yates, J. R. Isobaric Labeling-Based Relative Quantification in Shotgun Proteomics. J. Proteome Res. 2014, 13 (12), 5293–5309.

(65) O’Connell, J. D.; Paulo, J. A.; O’Brien, J. J.; Gygi, S. P. Proteome-Wide Evaluation of Two Common Protein Quantification Methods. J. Proteome Res. 2018, 17 (5), 1934–1942.

(66) Sun, S.; Homer, J. A.; Smedley, C. J.; et al. Phosphorus fluoride exchange: Multidimensional catalytic click chemistry from phosphorus connective hubs. Chem 2023, 9 (8), 2128–2143.

(67) Chappell, W. P.; Schur, N.; Vogel, J. A.; et al. Poison to promise: The resurgence of organophosphorus fluoride chemistry. Chem 2024, 10 (6), 1644–1654.

(68) Hahm, H. S.; Toroitich, E. K.; Borne, A. L.; et al. Global targeting of functional tyrosines using sulfur-triazole exchange chemistry. Nat. Chem. Biol. 2020, 16 (2), 150–159.

(69) Toroitich, E. K.; Ciancone, A. M.; Hahm, H. S.; et al. Discovery of a Cell-Active SuTEx Ligand of Prostaglandin Reductase 2. ChemBioChem 2021, 22 (12), 2134–2139.

(70) Borne, A. L.; Brulet, J. W.; Yuan, K.; Hsu, K.-L. Development and biological applications of sulfur– triazole exchange (SuTEx) chemistry. RSC Chem. Biol. 2021, 2 (2), 322-337, 10.1039/D0CB00180E.

(71) Brulet, J. W.; Borne, A. L.; Yuan, K.; et al. Liganding Functional Tyrosine Sites on Proteins Using Sulfur–Triazole Exchange Chemistry. J. Am. Chem. Soc. 2020, 142 (18), 8270–8280.

(72) Huang, T.; Hosseinibarkooie, S.; Borne, A. L.; et al. Chemoproteomic profiling of kinases in live cells using electrophilic sulfonyl triazole probes. Chem. Sci. 2021, 12 (9), 3295-3307, 10.1039/D0SC06623K.

(73) McCloud, R. L.; Yuan, K.; Mahoney, K. E.; et al. Direct Target Site Identification of a Sulfonyl– Triazole Covalent Kinase Probe by LC-MS Chemical Proteomics. Anal. Chem. 2021.

(74) Justin Grams, R.; Yuan, K.; Founds, M. W.; et al. Imidazoles are Tunable Nucleofuges for Developing Tyrosine-Reactive Electrophiles. ChemBioChem 2024, 25 (16), e202400382.

(75) Ciancone, A. M.; Seo, K. W.; Chen, M.; et al. Global Discovery of Covalent Modulators of Ribonucleoprotein Granules. J. Am. Chem. Soc. 2023, 145 (20), 11056–11066.

(76) Ciancone, A. M.; Hosseinibarkooie, S.; Bai, D. L.; et al. Global profiling identifies a stress-responsive tyrosine site on EDC3 regulating biomolecular condensate formation. Cell Chem. Biol. 2022, 29 (12), 1709-1720.e1707.

(77) Founds, M. W.; Murtagh, O. L.; Grams, R. J.; et al. Human PTGR2 Inactivation Alters Eicosanoid Metabolism and Cytokine Response of Inflammatory Macrophages. ACS Chem. Biol. 2025, 20 (6), 1426–1434.

(78) Grams, R. J.; Wolfe, W. J.; Seal, R. J.; et al. Discovery and Optimization of a Covalent AKR1C3 Inhibitor. J. Med. Chem. 2025, 68 (9), 9465–9478.

(79) Cruite, J. T.; Nowak, R. P.; Donovan, K. A.; et al. Covalent Stapling of the Cereblon Sensor Loop Histidine Using Sulfur-Heterocycle Exchange. ACS Med. Chem. Lett. 2023, 14 (11), 1576–1581.

(80) Talele, T. T. The “Cyclopropyl Fragment” is a Versatile Player that Frequently Appears in Preclinical/Clinical Drug Molecules. J. Med. Chem. 2016, 59 (19), 8712–8756.

(81) Okondo, M. C.; Johnson, D. C.; Sridharan, R.; et al. DPP8 and DPP9 inhibition induces pro-caspase-1-dependent monocyte and macrophage pyroptosis. Nat. Chem. Biol. 2017, 13 (1), 46–53.

(82) Okondo, M. C.; Rao, S. D.; Taabazuing, C. Y.; et al. Inhibition of Dpp8/9 Activates the Nlrp1b Inflammasome. Cell Chem. Biol. 2018, 25 (3), 262-267.e265.

(83) Lankas, G. R.; Leiting, B.; Roy, R. S.; et al. Dipeptidyl Peptidase IV Inhibition for the Treatment of Type 2 Diabetes: Potential Importance of Selectivity Over Dipeptidyl Peptidases 8 and 9. Diabetes 2005, 54 (10), 2988–2994.

(84) Waumans, Y.; Baerts, L.; Kehoe, K.; et al. The Dipeptidyl Peptidase Family, Prolyl Oligopeptidase and Prolyl Carboxypeptidase in the Immune System and Inflammatory Disease, including Atherosclerosis. Front. Immunol. 2015, Volume 6 - 2015, Review.

(85) Dance, I. Distance criteria for crystal packing analysis of supramolecular motifs. New J. Chem. 2003, 27 (1), 22-27, 10.1039/B206867B.

(86) Newberry, R. W.; Raines, R. T. The n→π* Interaction. Acc. Chem. Res. 2017, 50 (8), 1838–1846.

(87) Ogasawara, D.; Konrad, D. B.; Tan, Z. Y.; et al. Chemical tools to expand the ligandable proteome: Diversity-oriented synthesis-based photoreactive stereoprobes. Cell Chem. Biol. 2024, 31 (12), 2138-2155.e2132.

(88) Chen, M.; Shin, M.; Ware, T. B.; et al. Endocannabinoid biosynthetic enzymes regulate pain response via LKB1–AMPK signaling. Proc. Natl. Acad. Sci. 2023, 120 (52), e2304900120.

(89) Wilkerson, J. L.; Ghosh, S.; Bagdas, D.; et al. Diacylglycerol lipase β inhibition reverses nociceptive behaviour in mouse models of inflammatory and neuropathic pain. Br. J. Pharmacol. 2016, 173 (10), 1678–1692.

(90) Buczynski, M. W.; Herman, M. A.; Hsu, K.-L.; et al. Diacylglycerol lipase disinhibits VTA dopamine neurons during chronic nicotine exposure. Proc. Natl. Acad. Sci. 2016, 113 (4), 1086–1091.

(91) Cao, J. K.; Viray, K.; Shin, M.; et al. ABHD6 Inhibition Rescues a Sex-Dependent Deficit in Motor Coordination in The HdhQ200/200 Mouse Model of Huntington’s Disease. J. Neurol. Neurol. Disord. 2021, 7 (1), 106.

(92) Manterola, A.; Bernal-Chico, A.; Cipriani, R.; et al. Re-examining the potential of targeting ABHD6 in multiple sclerosis: Efficacy of systemic and peripherally restricted inhibitors in experimental autoimmune encephalomyelitis. Neuropharmacology 2018, 141, 181–191.

(93) Manterola, A.; Bernal-Chico, A.; Cipriani, R.; et al. Deregulation of the endocannabinoid system and therapeutic potential of ABHD6 blockade in the cuprizone model of demyelination. Biochem. Pharmacol. 2018, 157, 189–201.

(94) Dong, J.; Krasnova, L.; Finn, M. G.; Sharpless, K. B. Sulfur(VI) Fluoride Exchange (SuFEx): Another Good Reaction for Click Chemistry. Angew. Chem. Int. Ed. 2014, 53 (36), 9430–9448.

(95) Homer, J. A.; Sun, S.; Koelln, R. A.; Moses, J. E. Protocol for producing phosphoramidate using phosphorus fluoride exchange click chemistry. STAR Protoc. 2024, 5 (1), 102824.

