## Supplementary Information for "Re-Engineering P(V) Chemical Warfare: Harnessing Stereogenic Phosphorus-Azoles for Protein Ligand Discovery *In Vivo*"

**This PDF file includes:**

**Supporting Figures**

**Materials and Methods**

**NMR Spectra**

**X-Ray Crystal Structure Data Tables**

**References**

#### Supporting Figures

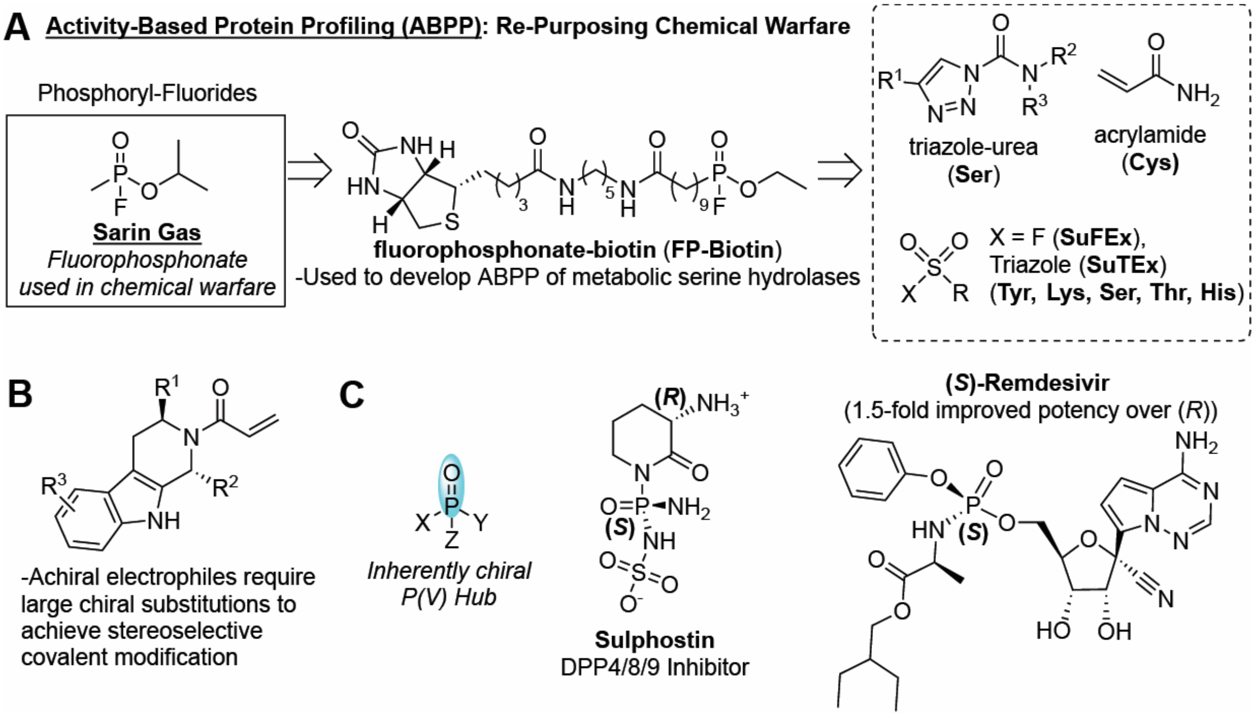


**Figure S1**. (**A**) Re-purposing of chemical warfare (sarin gas) to profile the metabolic serine hydrolases using fluorophosphonate-biotin (FP-biotin) that provided a platform for development of novel electrophiles targeting other nucleophilic amino acids. (**B**) Markush drawing of sterically hindered tryptoline acrylamides developed by the Cravatt lab to characterized stereoselective covalent modification of cysteines. (**C**) Drawing of a chiral phosphoryl moiety and representative examples sulphostin, a DPP4/8/9 targeting natural product, and (*S*)-remdesivir, an FDA-approved antiviral pro-drug.

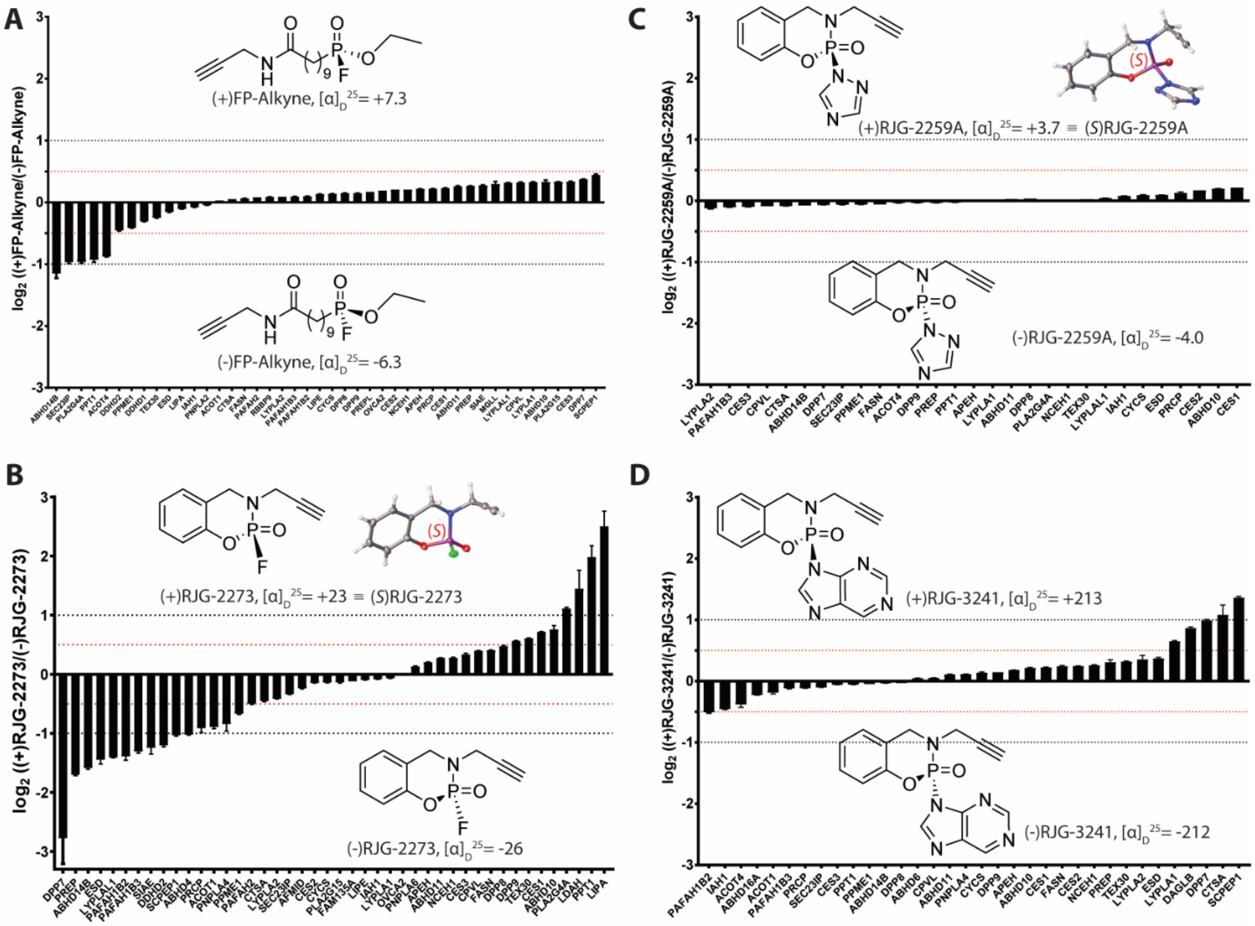

**Figure S2**. Bar graphs showing ratios for enantioselectively enriched serine hydrolases using (**A**) (+)- or (-)FP-Alkyne, (**B**) (+)- or (-)RJG-2273, (**C**) (+)- or (-)RJG-2259A, or (**D**) (+)- or (-)RJG-3241. Images of crystal structures for (+)RJG-2273 and (+)RJG-2259A are shown in (**B**) and (**C**), respectively, revealing (*S*)-configuration at phosphorus. The red and black dashed lines represent >25% and >50% enantioselective enrichment by (+) or (-)-enantiomers, respectively. Data represents three independent biological replicates.

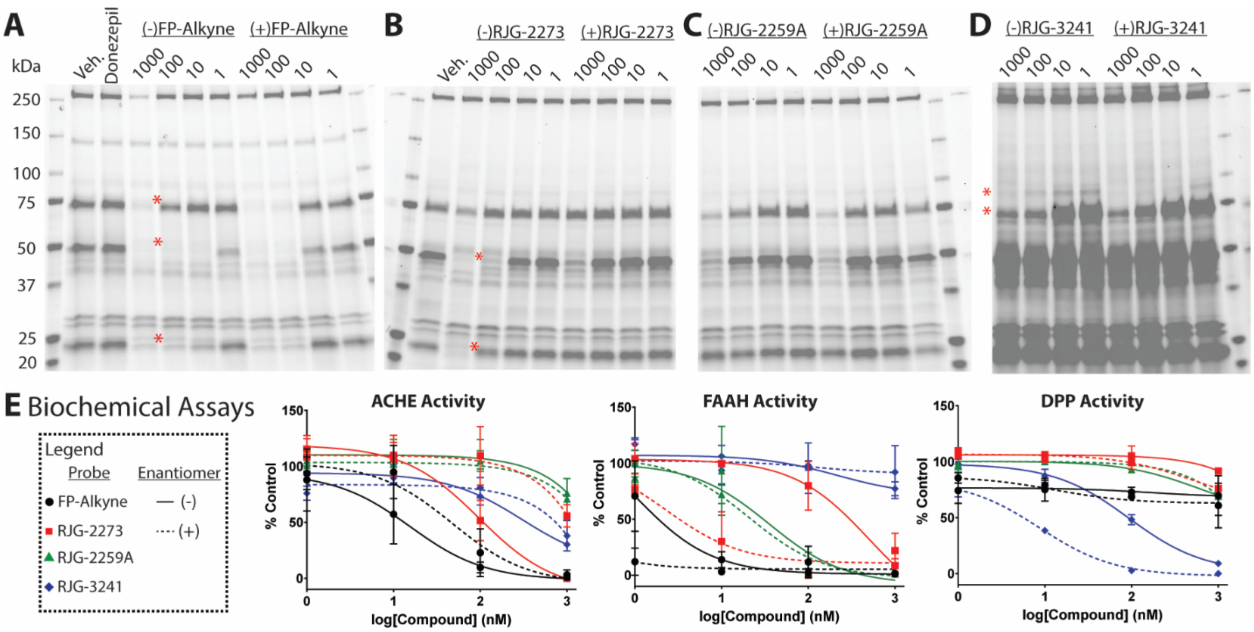


**Figure S3**. Full length SDS-PAGE gel showing dose response (1 – 1000 nM) after treating live N2a cells with (**A**) donepezil (1000 nM), (-) or (+)FP-Alkyne, (**B**) (-) or (+)RJG-2273, (**C**) (-) or (+)RJG-2259A, or (**D**) (-) or (+)RJG-3241. Red asterisks highlight proteins enantioselectively competed (-) or (+) probes. (**E**) Legend for graphs showing inhibition of ACHE, FAAH, or DPP biochemical activity using same lysate used to generate SDS-PAGE images in (**A-D**). Data represents two independent biological replicates.

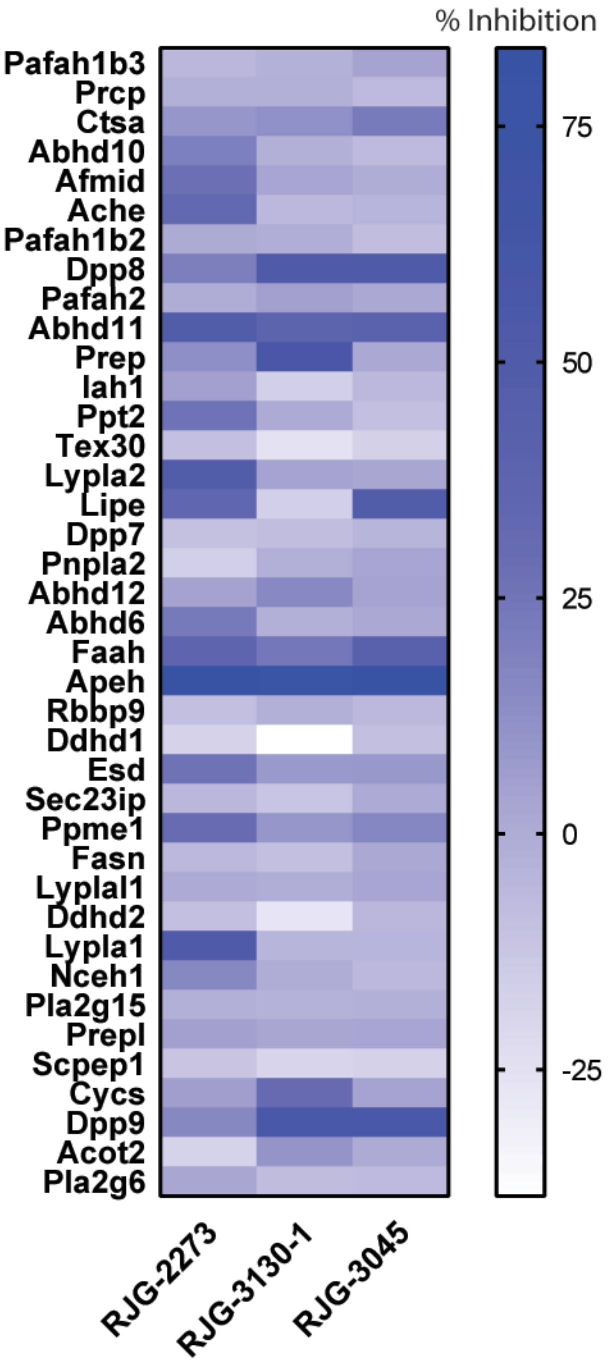


**Figure S4**. Heat map of serine hydrolases competed by RJG-2273, RJG-3130-1, or RJG-3045. In brief, N2a cells were treated with 1 µM compound in serum-free DMEM (0.1% DMSO final) for 4 hours at 37 °C. Data is representative of three independent biological replicates.

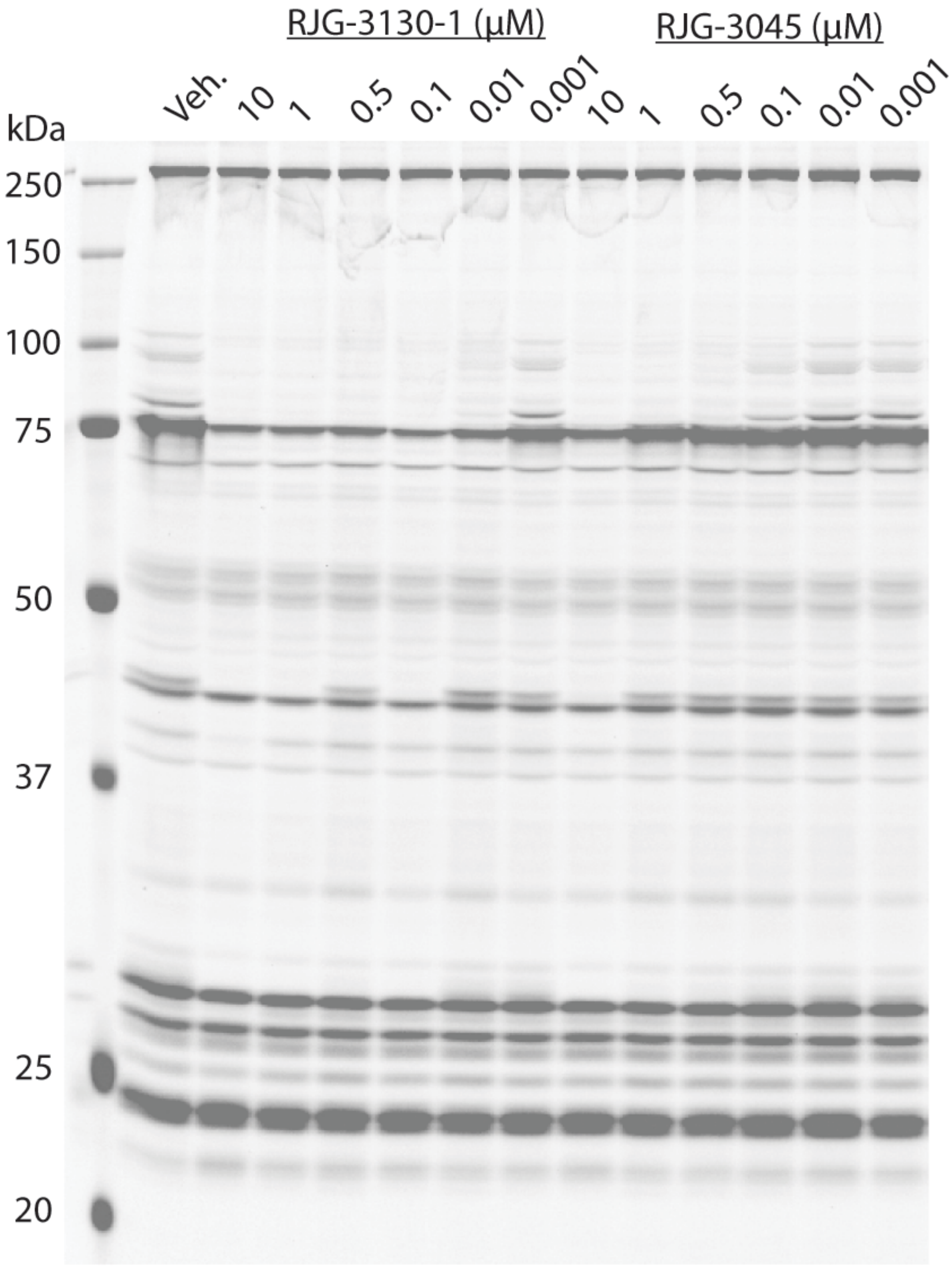


**Figure S5**. Full length SDS-PAGE gel showing dose response of RJG-3130-1 and RJG-3045. Data is representative of two independent biological replicates.

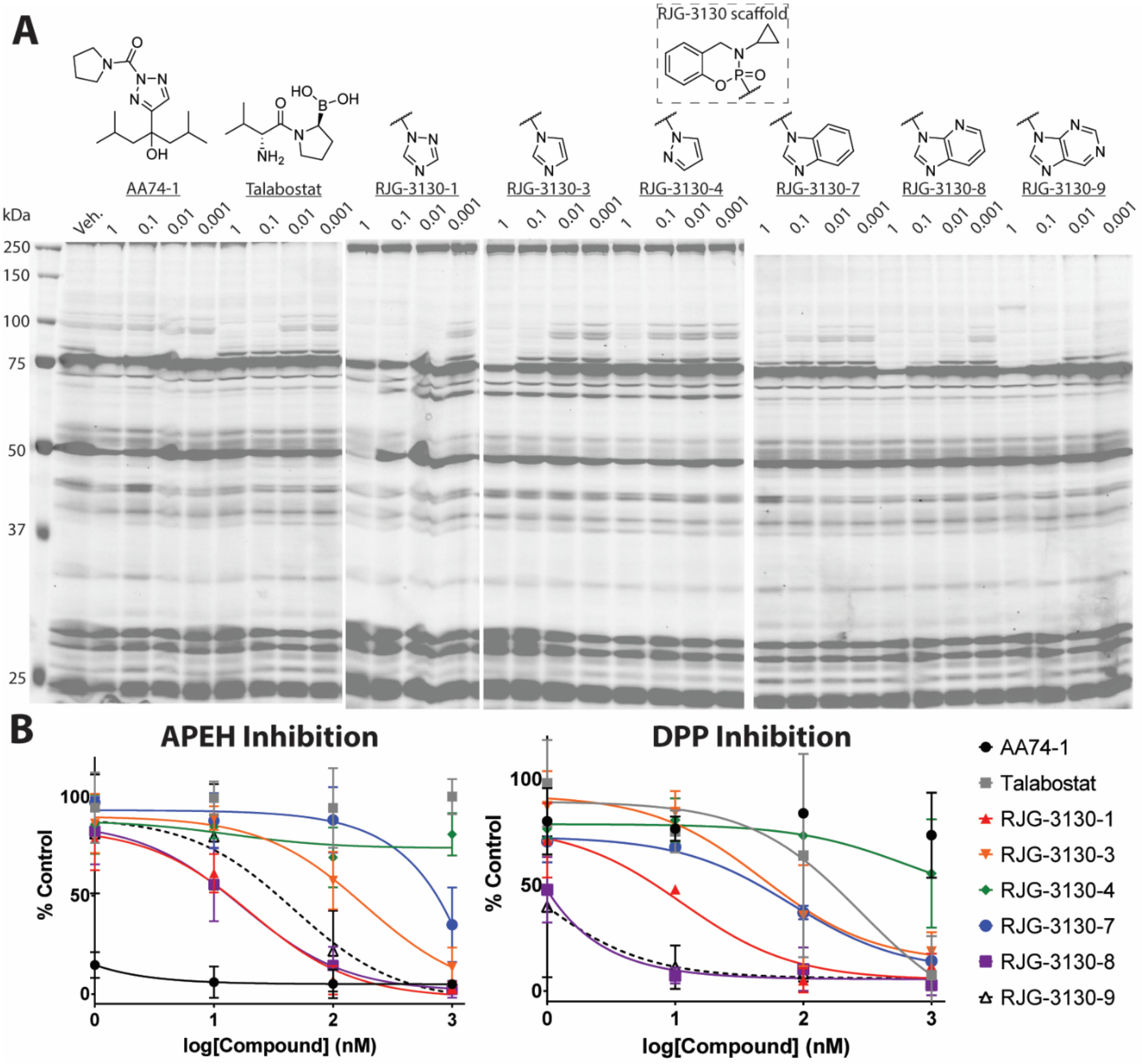


**Figure S6**. (**A**) Full length SDS-PAGE gels showing dose response of AA74-1 (APEH inhibitor control), talabostat (DPP8 & 9 inhibitor control), RJG-3130-1 (triazole leaving group (LG)), RJG-3130-3 (imidazole LG), RJG-3130-4 (pyrazole LG), RJG-3130-7 (benzimidazole LG), RJG-3130-8 (4-azabenzimidazole LG), and RJG-3130-9 (purine LG). (**B**) Graphs quantifying inhibition of FP-Rh probe labeling *via* competition with controls or PhAzE ligands ($IC_{50}$ values are listed in Figure S6). AA74-1 is an ultrapotent, irreversible covalent APEH ligand and talabostat is a reversible covalent DPP4/8/9 ligand. Data is representative of 2-3 independent biological replicates.

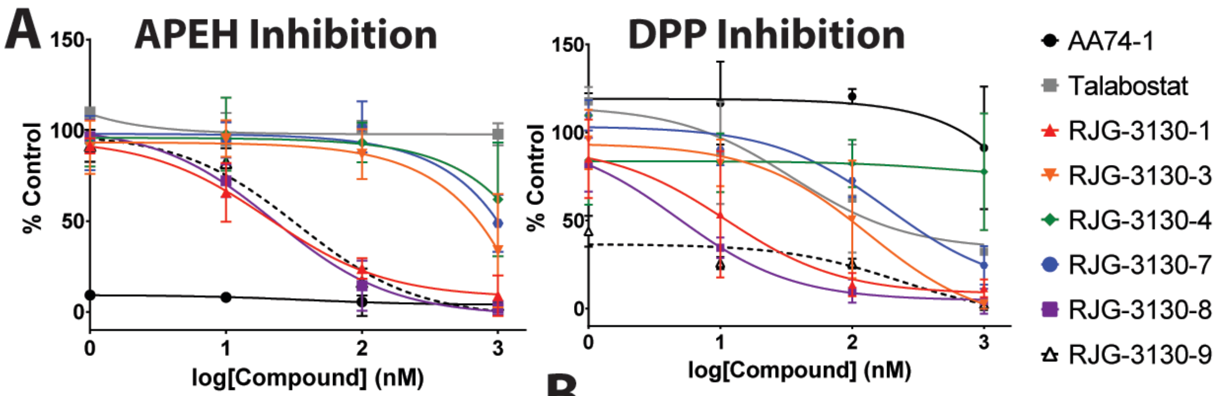


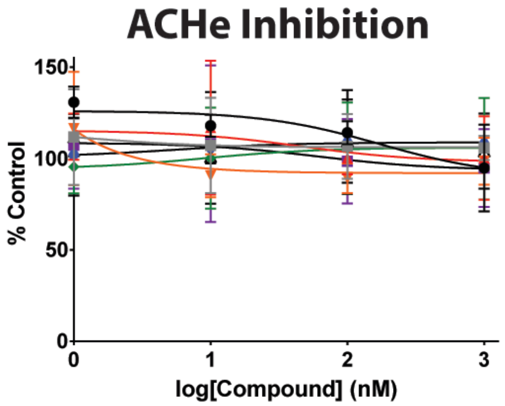


**B**

| Compound Name | APEH Inhibition ($IC_{50}$, nM) | | DPP Inhibition ($IC_{50}$, nM) | |
|---|---|---|---|---|
| | SDS-PAGE | Biochemical | SDS-PAGE | Biochemical |
| AA74-1 | <1 | <1 | >1000 | >1000 |
| Talabostat | >1000 | >1000 | 28 | 35 |
| RJG-3130-1 | 20 | 21 | 11 | 11 |
| RJG-3130-3 | 190 | 541 | 48 | 134 |
| RJG-3130-4 | >1000 | >1000 | >1000 | 569 |
| RJG-3130-7 | >1000 | >1000 | 80 | 285 |
| RJG-3130-8 | 17 | 23 | <1 | 5.2 |
| RJG-3130-9 | 47 | 34 | <1 | <1 |

**Figure S7**. (**A**) Graphs of APEH, DPP, and ACHE assays utilizing the same lysate to perform FP-Rh competition experiment in Figure S5. (**B**) Table showing $IC_{50}$ values quantified using nonlinear regression curves for SDS-PAGE (Figure S5B) and biochemical assays (Figure S6A). Data is representative of 2-3 independent biological replicates.

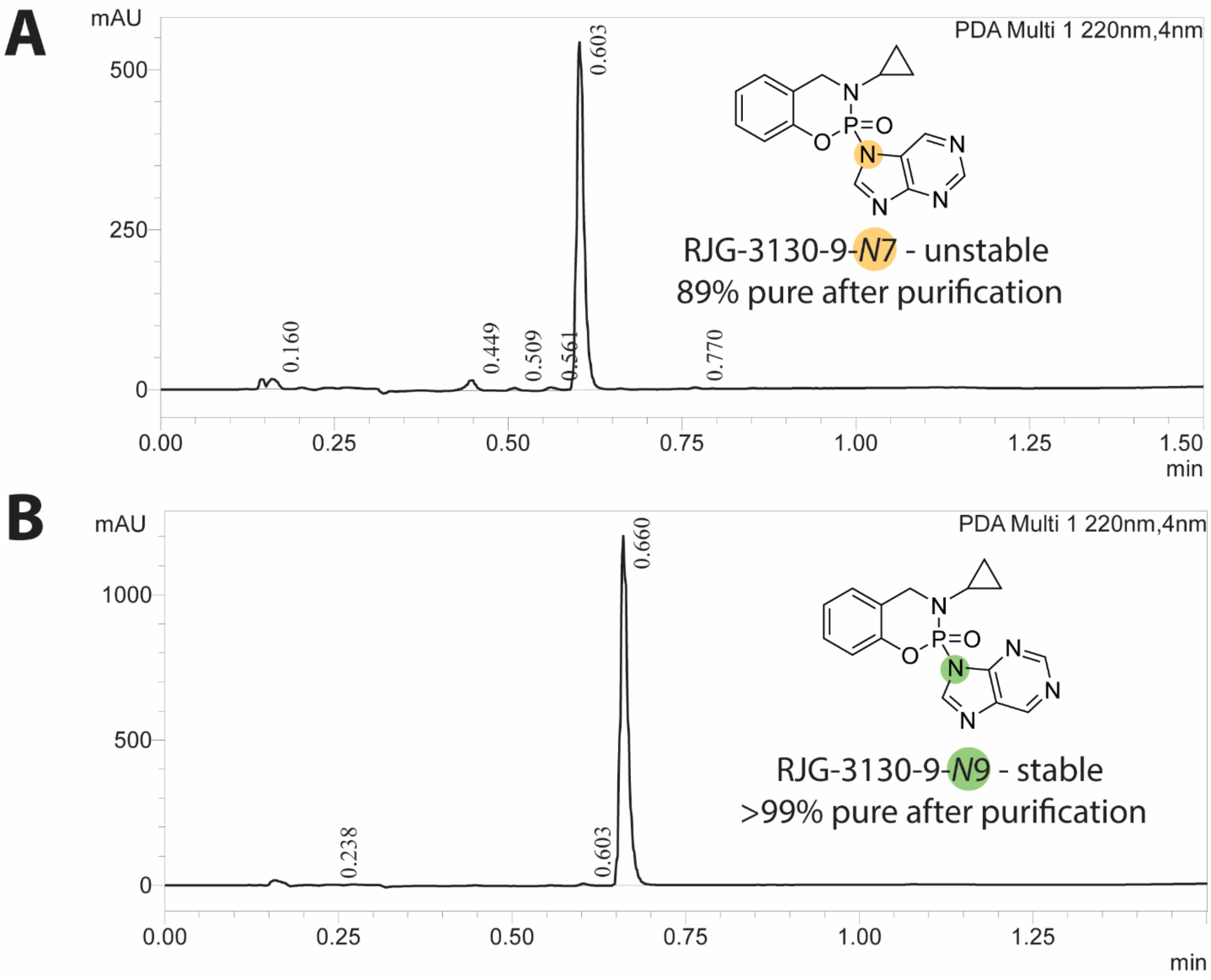


**Figure S8**. HPLC chromatograms after purification of the presumed (**A**) *N*7- and (**B**) *N*9-regioisomers of RJG-3130-9. A 1.5 min. gradient of 5%-95%B on a Shimadzu LCMS2020 equipped with a 30*3.0mm,3µm L-column 3 was used to determine purity, where mobile phase A is 5 mM $NH_4HCO_3$ and mobile phase B is acetonitrile.

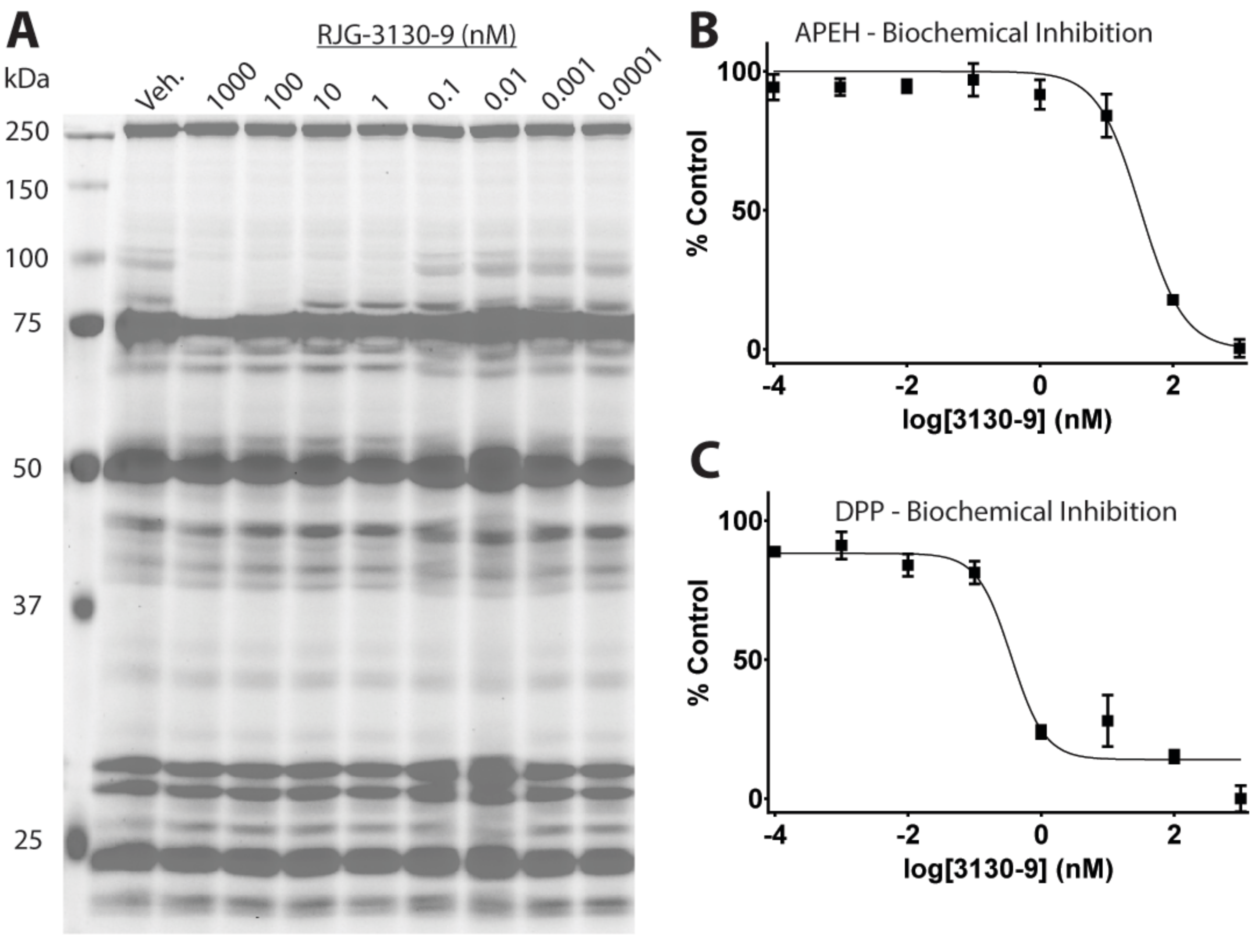


**Figure S9**. (**A**) Full length SDS-PAGE gel showing dose response of RJG-3130-9 in live N2a cells. Dose response graphs showing inhibition of APEH (**B**) and DPP (**C**) as determined by biochemical assays.

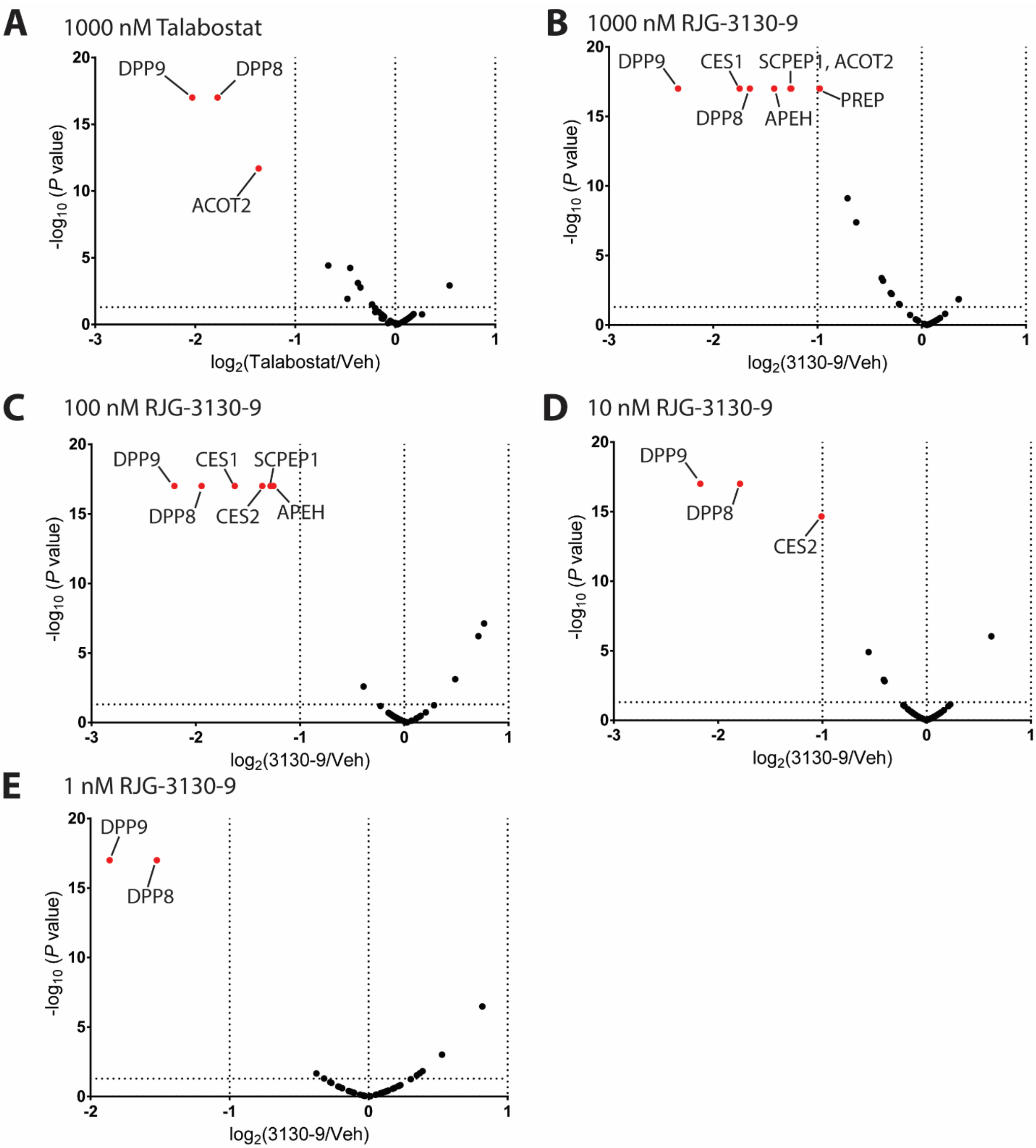


**Figure S10**. Volcano plots depicting inhibition of FP-biotin probe labeling after treating live THP1 cells with (**A**) 1000 nM talabostat or 1000, 100, 10, or 1 nM RJG-3130-9 (**B**-**E**, respectively). Proteins inhibited by >50% are highlighted in red.

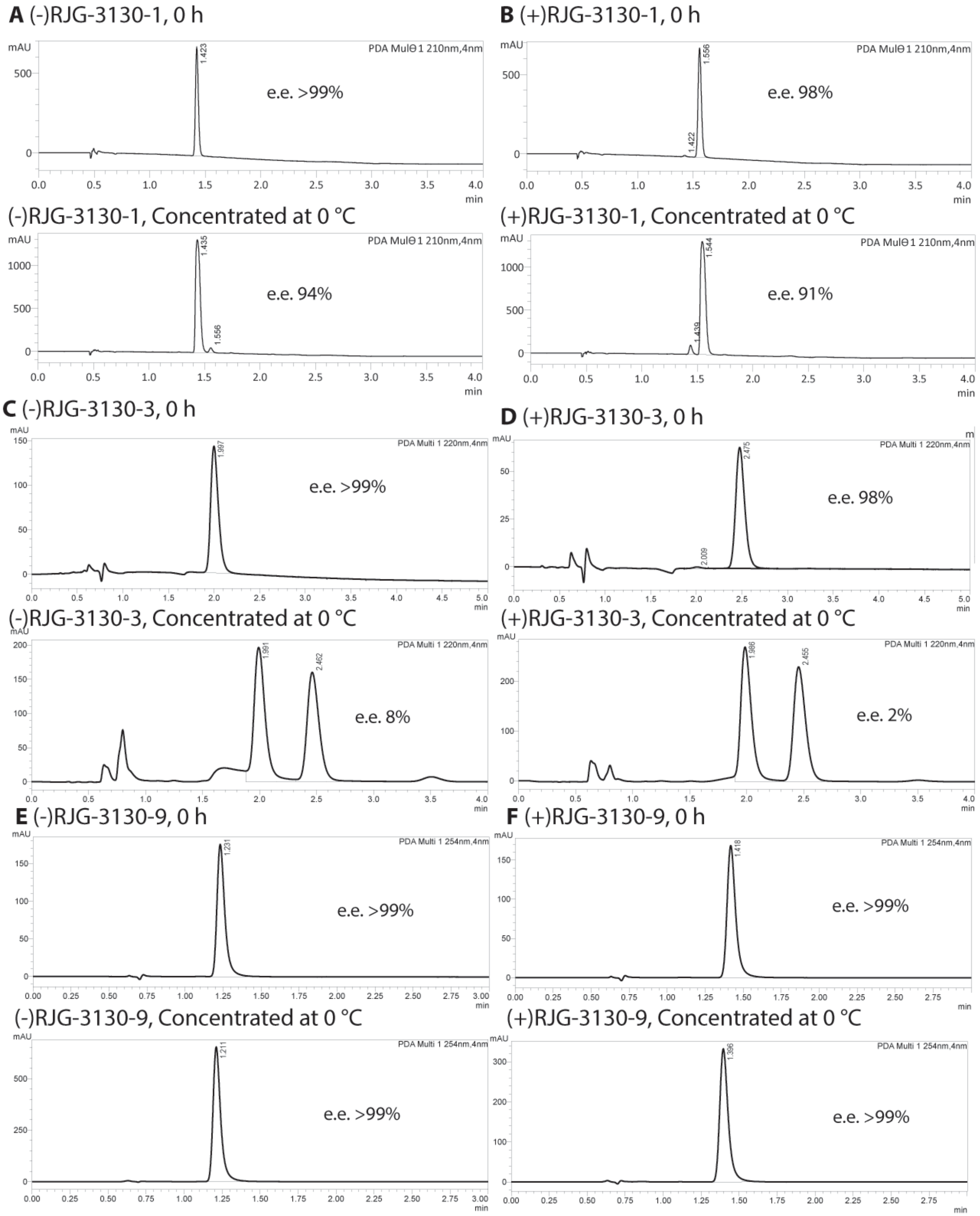


**Figure S11**. SFC chromatograms characterizing e.e. of (**A**) (-)- or (**B**) (+)-RJG-3130-1, (**C**) (-)- or (**D**) (+-RJG-3130-3, or (**E**) (-)- or (**F**) (+)-RJG-3130-9 after 0 h (in solution after purification) and after concentrating solution at 0 °C to obtain dry compound.

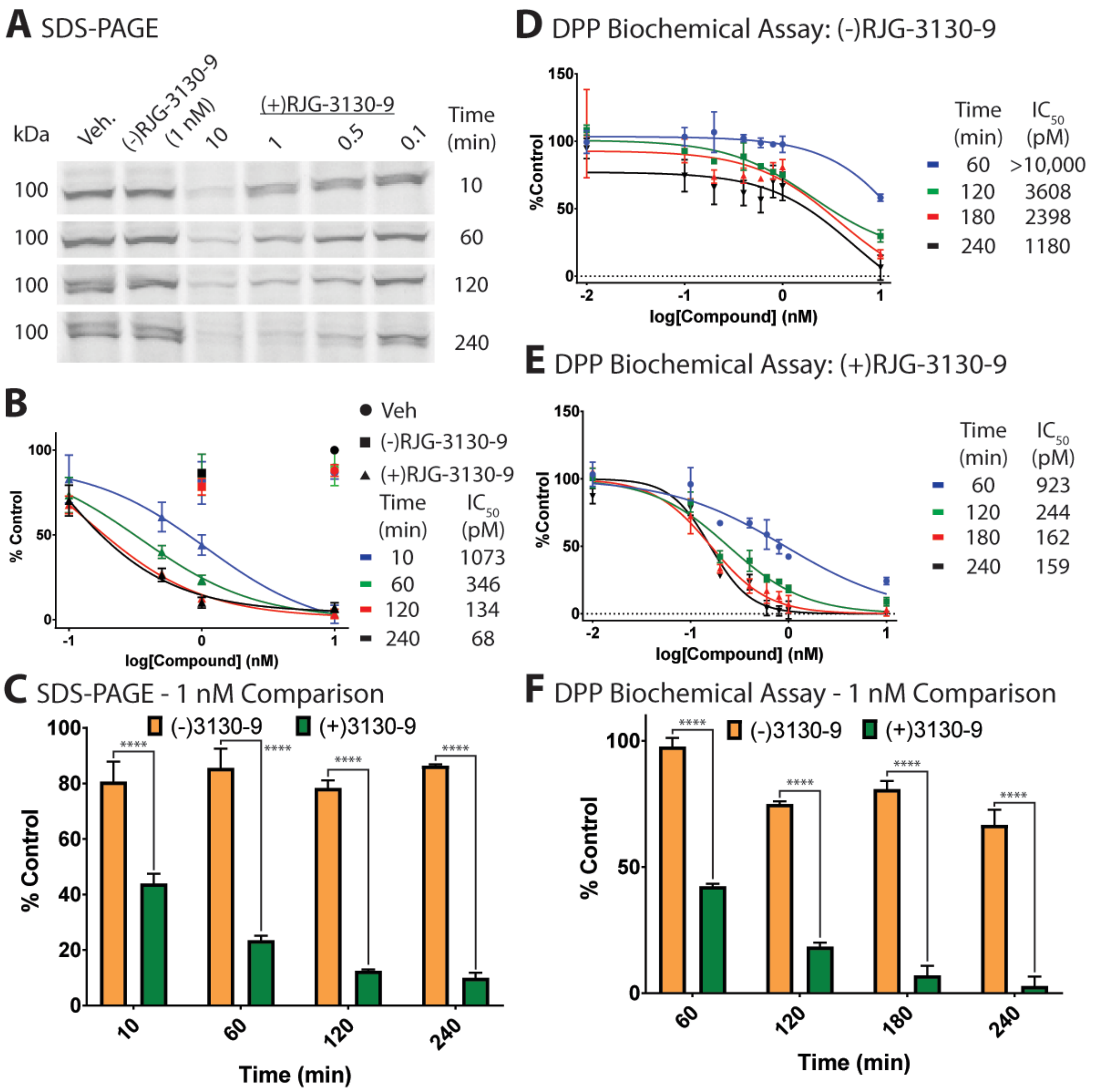


**Figure S12**. *In vitro* DPP inhibition to support irreversible covalent inhibition. (**A**) Competition-based SDS-PAGE experiment to determine potency of covalent inhibition of (-)- or (+)RJG-3130-9. (**B**) Graph of dose response curves derived from comparing RJG-3130-9 and vehicle fluorescence values from Figure 11A. (**C**) Bar graph showing inhibition of DPPs after THP1 soluble lysate with 1 nM (-)- or (+)RJG-3130-9 for 10 – 240 minutes. (**D**) Graph showing dose response curves of a DPP biochemical assay after THP1 soluble lysate was treated with vehicle or (-)RJG-3130-9 for 60 – 240 minutes. (**E**) Graph showing dose response curves of a DPP biochemical assay after THP1 soluble lysate was treated with vehicle or (+)RJG-3130-9 for 60 – 240 minutes. (**F**) Bar graph showing inhibition of DPPs after THP1 soluble lysate with 1 nM (-)- or (+)RJG-3130-9 for 60 – 240 minutes. Data is representative of three independent biological replicates.

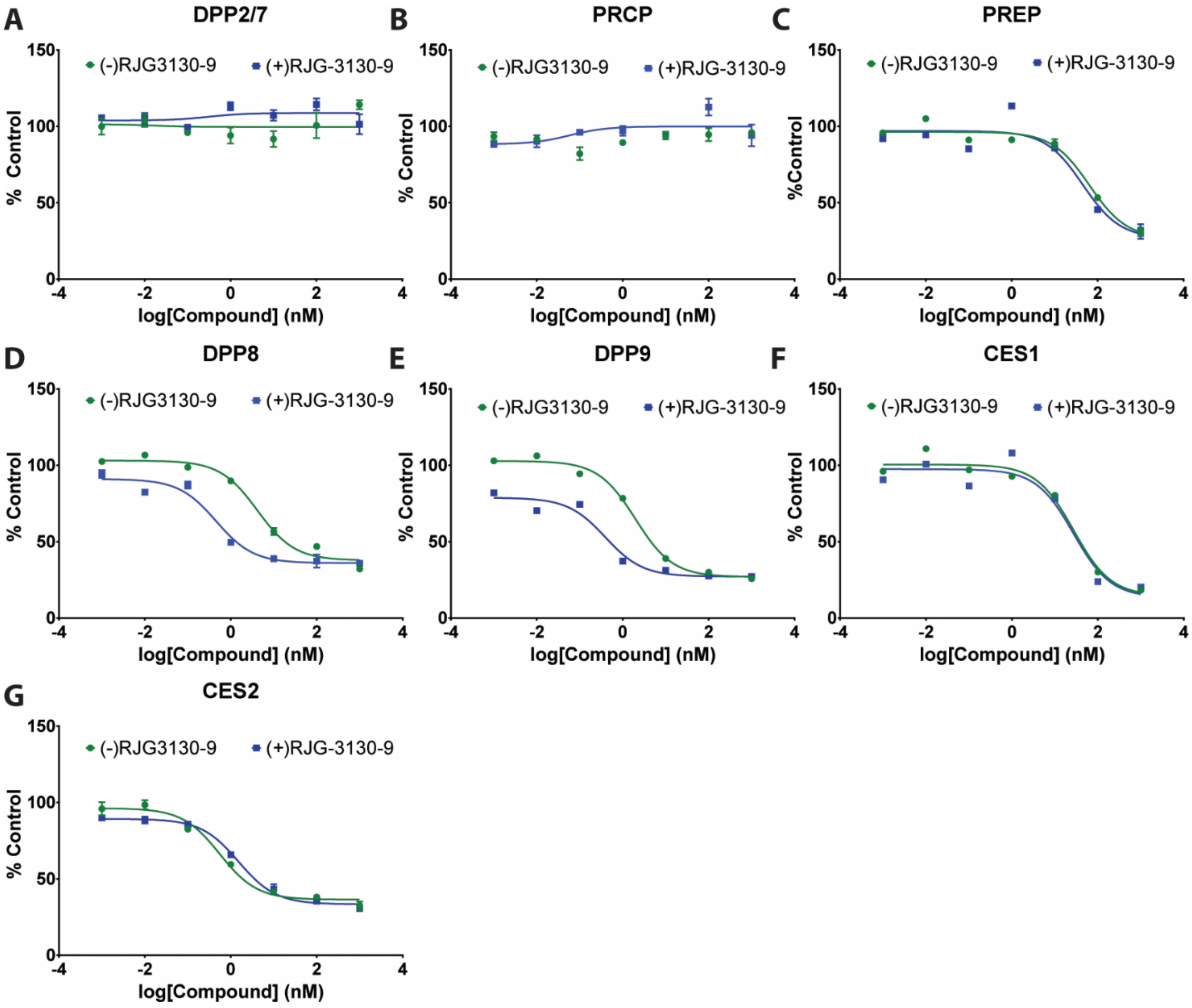


**Figure S13**. Inhibition of DPP (**A** – **E**) and CES (**F** – **G**) proteins in live THP1 cells. In brief, live THP1 cells were treated with vehicle or a dose response of (-)- or (+)RJG-3130-9 (*e.g.*, 1000, 100, 10, 1, 0.1, 0.01, or 0.001 nM) for 4 hours at 37 °C. LC-MS/MS samples were prepared as described below using TMTpro 16-plex for multiplexing and quantitation. Data is representative of three independent biological replicates.

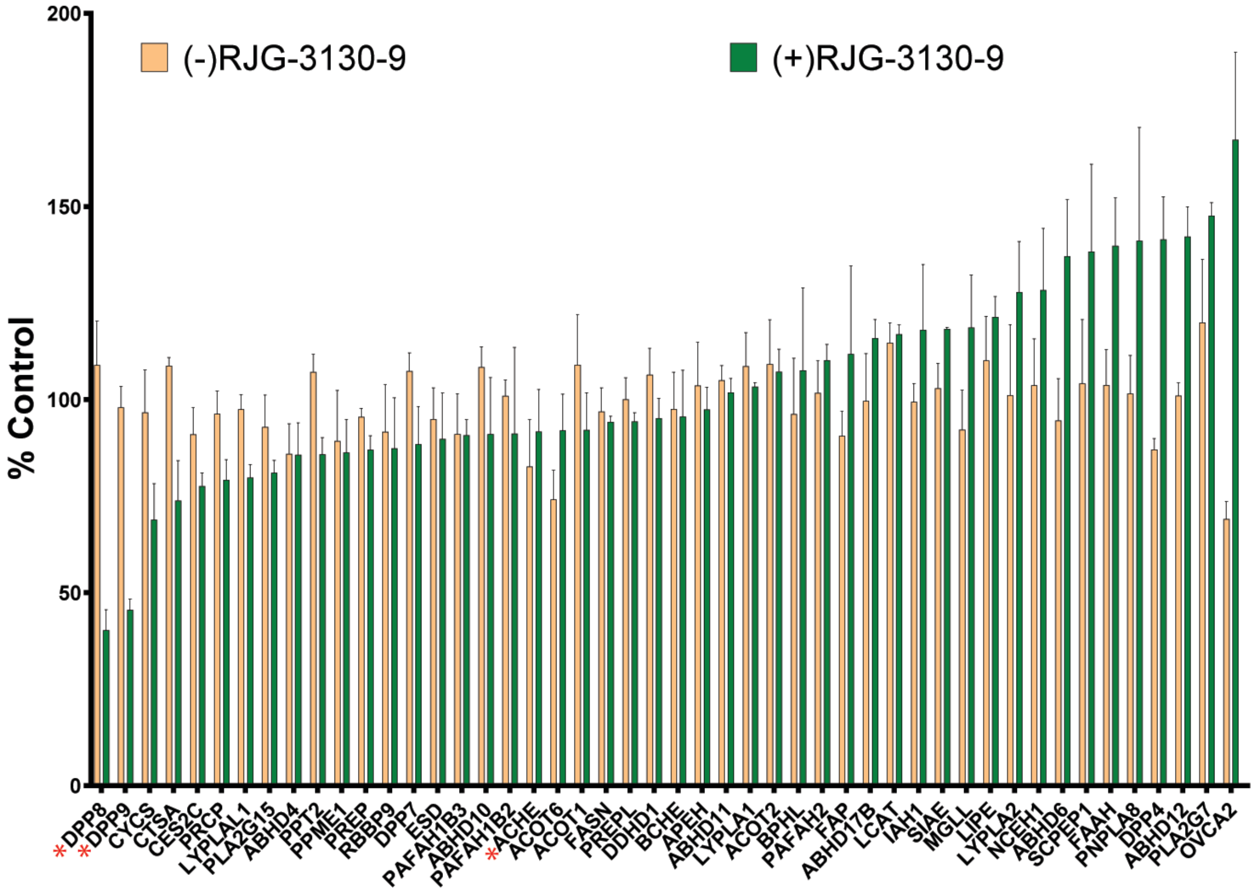


**Figure S14**. Bar graph of serine hydrolases enriched from brain tissue of mice treated with vehicle or 1 mg/kg of (-)- or (+)RJG-3130-9 for 4 hours. Samples were combined with TMT 6-plex and analyzed by LC-MS/MS. The red asterisks highlight >50% inhibition of FP-biotin probe labeling of DPP8/9 by (+)RJG-3130-9 compared to no inhibition observed with (-)RJG-3130-9, and <20% inhibition of ACHE. Data is representative of two independent biological replicates.

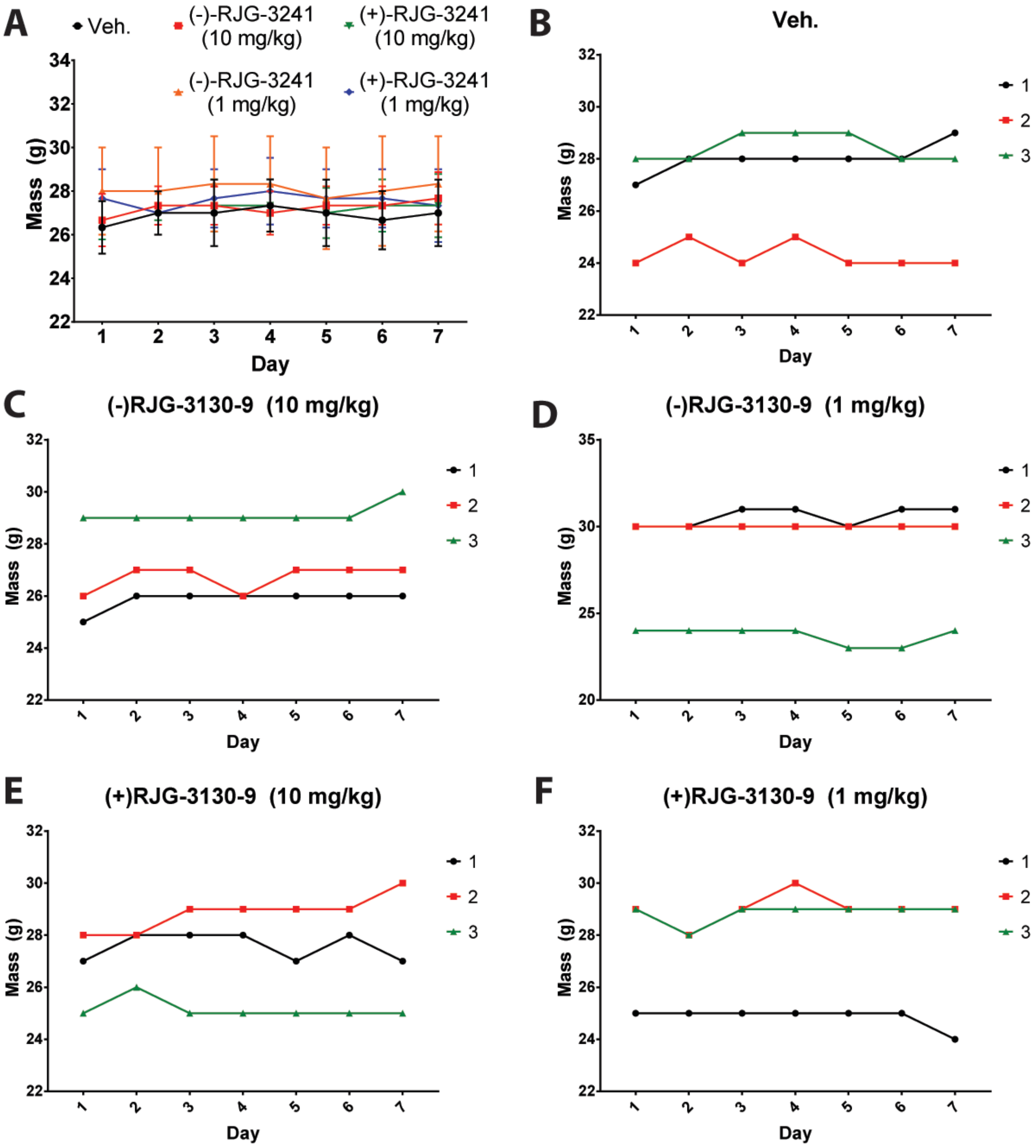


**Figure S15**. Average (**A**) or individual mouse weights over seven day treatment with (**B**) vehicle, (**C**) 10 mg/kg (-)RJG-3130-9, (**D**) 1 mg/kg (-)RJG-3130-9, (**E**) 10 mg/kg (+)RJG-3130-9, or (**F**) 1 mg/kg (+)RJG-3130-9.

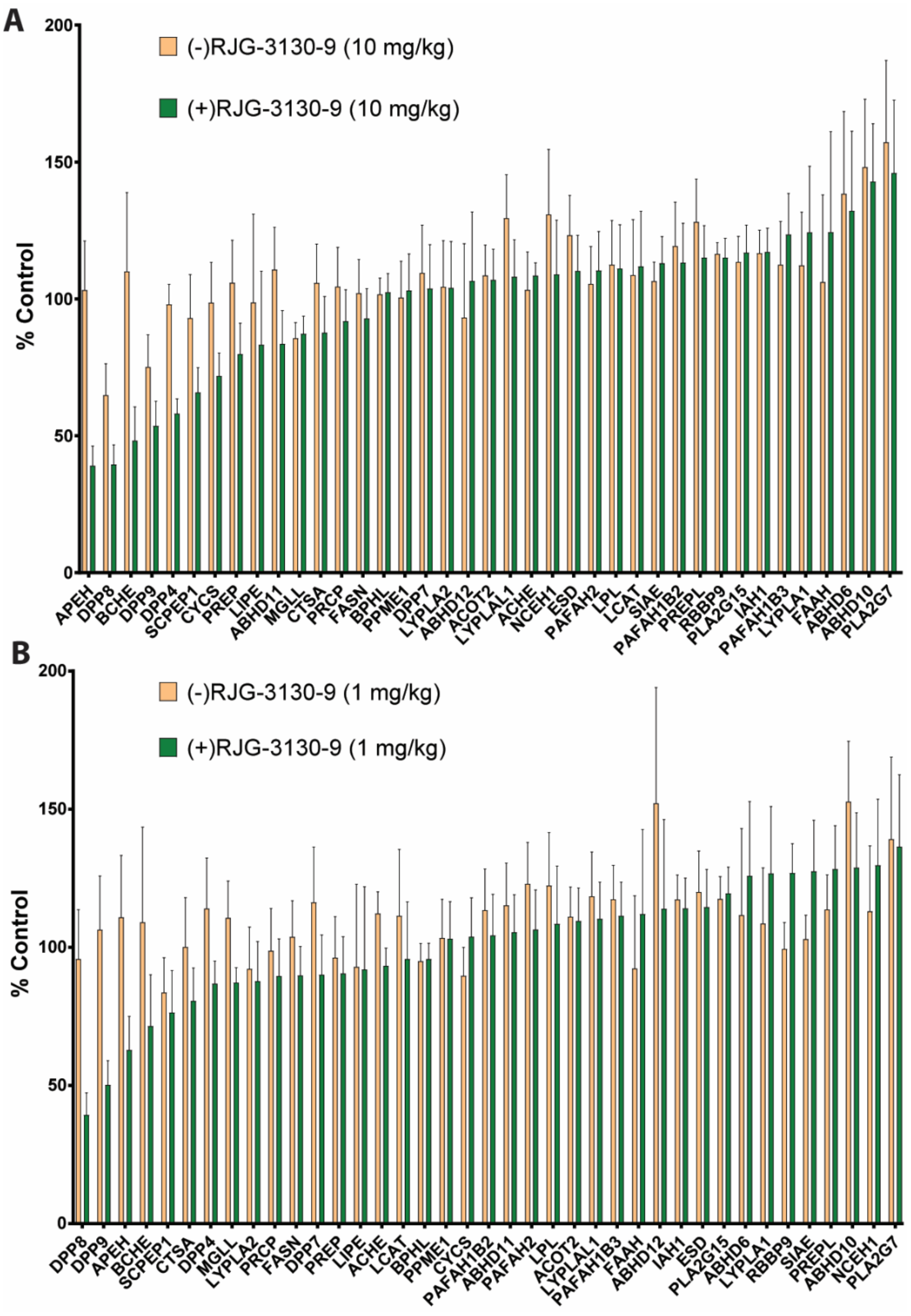


**Figure S16**. Bar graph of serine hydrolases enriched from brain tissue of mice treated with vehicle, (**A**) 10 mg/kg, or (**B**) 1 mg/kg of (-)- or (+)RJG-3130-9 for 7 days. Samples were combined with TMT 16-plex and analyzed by LC-MS/MS. Data is representative of three independent biological replicates.

#### Biological Methods

##### Cell culture

Cell lines were cultured at 37 °C with 5% $CO_2$ with manufacturer recommended media supplemented with 10% fetal bovine serum (US Source, Omega Scientific) and 1% L-glutamine (Fisher Scientific): THP1: RPMI + 1% MEM non-essential amino acids (Gibco); N2a: DMEM. Cells were collected or treated for experimental use when they reached ~90% confluency. The media was aspirated, cells washed with cold PBS (2×) and scraped from plates. The cells were pelleted by centrifugation at 400g for 5 min, snap-frozen using liquid nitrogen and stored at −80 °C until further use.

##### Mice

All animal experiments were approved by the Animal Resources Center of the University of Texas at Austin. All animals were allowed free access to a standard chow diet and water. The animals were housed according to the IACUC policy on social housing of animals. In all mouse experiments, age-matched and weight-matched mice were randomized to each group to eliminate age and weight differences. Mice were house in an animal facility of the University of Texas at Austin under constant environmental conditions (RT, 21 ± 1 °C; relative humidity, 40–70%; a 12 h light–dark cycle). Mice were administered intraperitoneally with 100 µL of vehicle using a 29 G needle (sterile saline : PEG40 Castor oil : 100% ethanol solution = 18 : 1 : 1) Mice weight was recorded daily to monitor potential drug toxicity.

##### Gel-Based Chemical Proteomic Assay

For *in situ* experiments, live cells were washed with serum-free media then treated with compound (1000x dilution into serum-free media, final [DMSO] = 0.1%) for 4 hours at 37 °C. Cell pellets were lysed in PBS by sonication and fractionated (100,000xg, 45 min, 4 °C) to generate soluble and insoluble fractions. Protein concentrations were determined using the Bio-Rad DC protein assay and adjusted to 1 mg $ml^{-1}$ in PBS. For *in vitro* experiments, proteome samples (50 µl aliquots) were treated with PhAzE ligands at the indicated concentrations (1 µl, 50× stock in DMSO) and incubated at 37 °C for respective times. Lysate was labeled with fluorophosphonate-rhodamine (FP-Rh) for 1 hour at room temperature then quenched with 17 µl of 4x SDS–PAGE loading buffer and ß-mercaptoethanol (ßME). Quenched samples were analyzed by SDS–PAGE and in-gel fluorescence scanning.

##### Live cell evaluation of phosphoryl-fluoride or PhAzE probes

Cells grown to ~90% confluency in 10 cm plates were treated with DMSO vehicle or compound (10 µl of 1,000× DMSO stock) in 10 mL serum-free media for the indicated concentrations and times at 37 °C with 5% $CO_2$. For THP1 monocytes, cells were pelleted and washed twice with serum-free RPMI. About 40-50x$10^6$ THP1 cells in 25 mL serum-free RPMI were transferred to 75 $cm^2$, then 25 mL containing 2x compound stock was added and the experiments incubated at 37

°C for 4 hours. After treatment, cells were washed with cold PBS twice then lysed in 1 mL cold 1X PBS. For LC–MS studies, protein concentrations were normalized to 2 mg $ml^{-1}$ and 500 µl (1 mg final protein amount) was used for sample preparation as detailed below.

**Preparation of proteomes for tandem mass tag LC-MS/MS chemical proteomics**

For alkyne-probe treated samples, copper-catalyzed azide-alkyne [3+2] cycloaddition (CuAAC) and chloroform/methanol extractions were used to remove click reagents as previously described.[1-2] For FP-biotin enrichment, lysate was treated with 5 µM FP-biotin for 1 hour at room temperature followed by chloroform/methanol extractions to remove excess reagent. Following resuspension in 6M Urea/ 25mM AmBic, proteins were reduced by dithiothreitol and alkylated with iodoacetamide as previously described.[1] Samples were diluted in 5 mL 1X PBS + 140 µL 10% SDS-PAGE. Probe-labeled proteins were enriched with avidin agarose beads by rotation for 1 hour at room temperature, followed by washing (3X with 10 mL 1X PBS, 3X with 10 mL LC/MS $H_2O$) to remove unbound protein. Avidin beads were suspended in 200 µL 25 mM ammonium bicarbonate and proteins digested overnight with Tryp-Lys-C in 25mM ammonium bicarbonate (2 µg, 37 °C). Supernatant was transferred to a biospin column and isolated *via* centrifugation (1,400 g x 3 min). The beads were washed with another 200 µL of 25 mM ammonium bicarbonate buffer then dried *via* SpeedVac. Dried peptides were re-suspended in 50 µL 0.1% FA in water and de-salted *via* STAGE tip. Peptides were eluted with 4:1 MeCN:$H_2O$ (0.1% TFA) and dried *via* SpeedVac.

**Biotin Enriched Peptide TMT labeling**

Dried peptides were re-suspended in 10 µL 200 mM EPPS, pH 8.5 then transferred to TMTsixplex™ reagents at a 2:1 w/w ratio of label to peptide. The final solution contains 15 µL (2:1 EPPS:MeCN). The reaction was incubated for 1 hour at room temperature with shaking at 500 rpm. The TMT reaction was quenched with 6% hydroxylamine (15 min, RT) and channels were mixed. Samples were dried using a SpeedVac and reconstituted in 50 µL 0.1% TFA in water and de-salted *via* STAGE tip. Peptides were eluted with 4:1 MeCN:$H_2O$ (0.1% TFA) and dried *via* SpeedVac. Dry, TMT-labeled samples were re-suspended in 25 µL 0.1% TFA in water (Mobile Phase A) and stored at -80 °C until analysis.

**LC–MS/MS analysis of samples**

Peptides were analyzed with nano-electrospray ionization-liquid chromatography-mass spectrometry (LC-MS/MS) on a Vanquish Neo UHPLC (ThermoFisher Scientific) coupled to an Orbitrap Exploris mass spectrometer. Peptides were separated with a 120 min gradient by reverse phase LC using 3 µm C18 (20 cm) as follows: (A: 0.1% formic acid/H2O; B: 80% ACN, 0.1% formic acid in H2O): 0–2 min 4% B, 450 nL/min; 2–78 min 25% B, 300 nL/min; 78–123 min 45% B, 300 nL/min; 123–125 min 99% B, 450 nL/min; 125-127 min 99% B, 450 nL/min; 127–130 min 1% B, 450 nL/min; 130–140 min 1% B, 450 nL/min. Data was acquired using a top 30 ddMS2 method. MS1 spectra were acquired at 120K resolution with a maximum injection time of 20 ms and MS2 spectra were taken with an AGC of 25K, quadrupole isolation width of 0.7m/z, HCD collision energy of 36, Orbitrap Resolution of 15K (TMT6-plex) or 45K (TMT16-plex), and max IT of 75 ms.

**LC-MS/MS data analysis for TMT-tagged SuTEx modified peptides**

Peptide identification and quantification was accomplished with Proteome Discoverer 3.0 and a PMI-Byonic node. MS2 spectra were searched against the human protein database (Uniprot, download date 01/17/2024) using the following parameters: ≤2 missed cleavages, 10 ppm precursor mass tolerance, 20 ppm fragment mass tolerance, and 1% protein false discovery rate. Modifications considered methionine oxidation (+15.9949, M, variable), cysteine carbamidomethylation (+57.021464, C, fixed), and TMT modification (TMT6plex (+229.1629) or TMTpro (+304.2071), N-term, K, variable). Peptides used for quantification met the following quality control criteria: PMI-Byonic Score ≥300, delta ppm err. 5, co-isolation interference threshold ≤ 50%, reporter ion S/N ≥10, and were present in n=2 biological replicates. Volcano plots were generated by grouping PSMs using the peptide isoform node with cutoffs of $Log_2$ FC≥0.5, and p-value ≤0.05.

**ACHE Activity Assay**

This procedure was adapted from published protocols.[3-4] N2a cells were treated with compound in serum-free DMEM for 4 hours at 37 °C. Cells were washed and harvested in cold 1X PBS. Soluble cell proteomes were diluted to 1.0 mg $ml^{-1}$ in assay buffer (100 mM $NaH_2PO_4$, pH 7.0) and 50 µL was added to each well of a 96 well plate. Then, 20 µL of a substrate solution containing 4.94 mM DTNB and 7.47 mM acetyl thiocholine in assay buffer was added to each well. The reaction was incubated at 37 °C for 60 minutes. Absorbance was measured at 415 nm on a BMG Labtech CLARIOstar plate reader.

**APEH Activity Assay**

This procedure was adapted from a published protocol.[5] N2a cells were treated with compound in serum-free DMEM for 4 hours at 37 °C. Cells were washed and harvested in cold 1X PBS. Soluble cell proteomes were diluted to 1.0 mg $ml^{-1}$ in assay buffer (100 mM $NaH_2PO_4$, pH 7.0) and 50 µL was added to each well of a 96 well plate. Then, 25 µL of a substrate solution containing 300 µM Ac-ala-pna (Arctom Scientific, AAB-AA00CHNT) in assay buffer was added to each well. The reaction was incubated at 37 °C for 60 minutes. Absorbance was measured at 405 nm on a BMG Labtech CLARIOstar plate reader.

**DPP Activity Assay**

This procedure was adapted from a published protocol.[6] N2a cells were treated with compound in serum-free DMEM for 4 hours at 37 °C. Cells were washed and harvested in cold 1X PBS. Soluble cell proteomes were diluted to 1.0 mg $ml^{-1}$ in assay buffer (100 mM $NaH_2PO_4$, pH 7.0) and 50 µL was added to each well of a 96 well plate. Then, 25 µL of a substrate solution containing 300 µM gly-pro-pna·TsOH (Chem-Impex, 03365) in assay buffer was added to each well. The reaction was incubated at 37 °C for 60 minutes. Absorbance was measured at 405 nm on a BMG Labtech CLARIOstar plate reader.

**Chemical Synthesis and Characterization**

All chemicals used were reagent grade and used as supplied, except where noted. Dichloromethane ($CH_2Cl_2$) and methanol (MeOH) were used without any further purification steps. Analytical thin layer chromatography (TLC) was performed on Merck silica gel 60 F254 plates (0.25 mm). Flash column chromatography was carried out using forced flow of the indicated solvent on Silica Gel 60 (230-400 mesh) purchased from Fisher Scientific. Compounds were visualized by UV-irradiation. $^1H$ and $^{13}C$ NMR spectra were recorded on a Varian Inova 500 (500 MHz), 600 (600 MHz), or Bruker Avance III 800 (800 MHz) spectrometers in $CDCl_3$, Acetone-$d_6$, or DMSO-$d_6$ with chemical shifts referenced to internal standards ($CDCl_3$: 7.26 ppm $^1H$, 77.16 ppm $^{13}C$ ; $(CD_3)_2CO$: 2.05 ppm $^1H$, 29.84 and 206.26 ppm $^{13}C$; $(CD_3)_2CO$: 2.50ppm $^1H$, 40.00 ppm $^{13}C$) unless stated otherwise. Splitting patterns are indicated as s, singlet; d, doublet; t, triplet; q, quartet; m, multiplet for 1 H-NMR data. NMR chemical shifts (δ) are reported in ppm and coupling constants (J) are reported in Hz. High resolution mass spectral (HRMS) data were obtained by an Agilent 6545B LC/Q-TOF (Agilent Technologies, Santa Clara, CA, USA). Chiral resolution was performed on a SFC150AP, SFC150 MGM, or a Gilson 281 HPLC equipped with a chiral column (See characterization below for column specifications). Optical rotation was acquired using an Anton Paar MCP4100 polarimeter with a quartz cell length of 100 mm and a wavelength of 589 nm.

**Synthesis and Experimental Data for Cyclic Phosphoramidotriazoles**

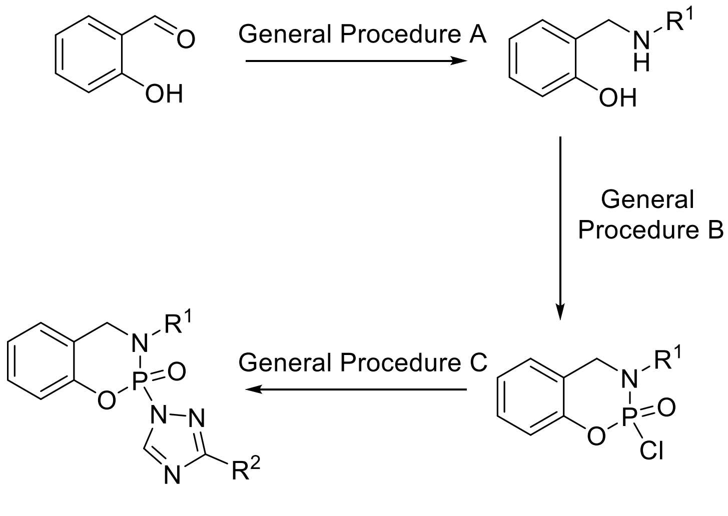


**General Procedure A[7]**

To a solution of amine (1.0 equiv) in MeOH (0.8 M) at room temperature was added sodium sulfate, salicylaldehyde (1.0 equiv) and *N*,*N*-diisopropylethylamine (1.0 equiv). The reaction was stirred at room temperature for 16 hours. Once complete, the reaction was diluted with MeOH to 0.4 M and cooled to 0 °C. Sodium borohydride (1.2 equiv) was slowly added. The resulting mixture slowly warmed to RT overnight. The reaction was diluted with water and $K_2CO_3$ added. The solution was stirred for 1 hour at room temperature. The aqueous layer was extracted three times with ethyl acetate. The combined organic layers were washed with brine, dried over sodium sulfate, then concentrated *in vacuo*. The crude material was purified by flash chromatography eluting with ethyl acetate-hexane to afford the desired aminophenol.

**General Procedure B**[7]

A solution of aminophenol (1.0 equiv) and triethylamine (2.0 equiv) in dry $CH_2Cl_2$ (0.2 M) was cooled to – 78 °C. After cooling for 15 minutes, phosphorous oxychloride (1.00 equiv, 1 M in $CH_2Cl_2$) was slowly added. The reaction was slowly warmed to room temperature overnight. The reaction was concentrated *in vacuo* and the crude material was purified by flash chromatography eluting with ethyl acetate-hexanes to afford the desired cyclic phosphoramidochloridate.

**General Procedure C:**

To a solution of phosphoramidochloridate (1.00 equiv) in $CH_2Cl_2$ (0.2 M) was added azole (1.00 equiv) and *N*,*N*-diisopropylethylamine (1.00 equiv). The reaction was stirred overnight. The reaction was concentrated *in vacou* and the crude material purified by flash chromatography with ethyl acetate-hexanes to afford the desired cyclic phosphoramidoazole.

**Characterization of Phosphoramidoazole (PhAzE) Probes**

**ethyl (10-oxo-10-(prop-2-yn-1-ylamino)decyl)phosphonofluoridate (FP-alkyne)**

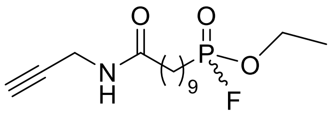

Prepared according to published methods and characterization data matches previously published data.[8]

**(-)FP-alkyne: $[\alpha]_D^{25}$** = -6 (0.1 g/mL, $CHCl_3$)

**(+)FP-alkyne: $[\alpha]_D^{25}$** = +7 (0.1 g/mL, $CHCl_3$)

***Determination of enantiomeric excess***

Enantiomeric excess was determined using CHIRALPAK AY-3 (100x4.6mm 3.0μm) column eluting with 50% B (Mobile Phases: A: supercritical $CO_2$; B: 50% MeCN in EtOH) over 4 minutes, 3.0 mL/min, (-)FP-alkyne: 1.785 min. (>99% e.e.), (+)FP-alkyne: 1.861 min. (98% e.e.).

(-)FP-Alkyne

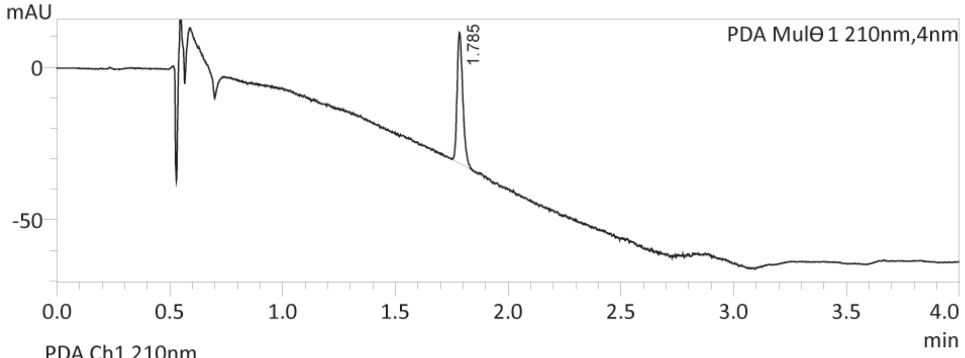


PDA Ch1 210nm

| Peak# | Ret. Time | Height | Width at 50% Height | Area | Area% | Resolution(JP) |
|---|---|---|---|---|---|---|
| 1 | 1.785 | 41981 | 0.027 | 75251 | 100.000 | -- |
| Total | | | | 75251 | 100.000 | |

(+)FP-Alkyne

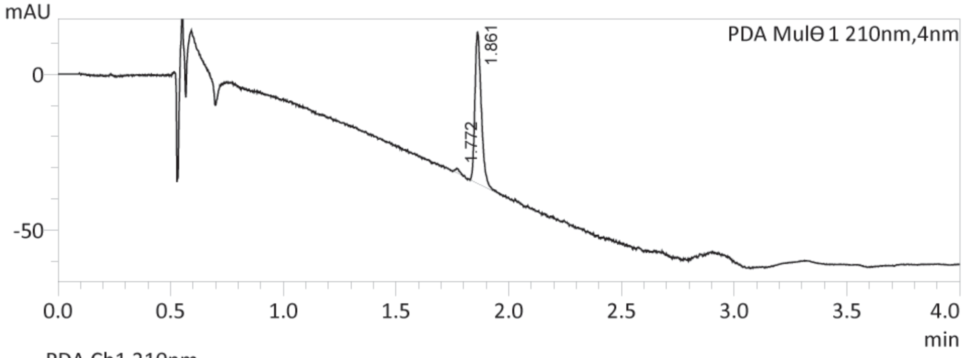


PDA Ch1 210nm

| Peak# | Ret. Time | Height | Width at 50% Height | Area | Area% | Resolution(JP) |
|---|---|---|---|---|---|---|
| 1 | 1.772 | 1370 | 0.020 | 1755 | 1.805 | -- |
| 2 | 1.861 | 47998 | 0.030 | 95475 | 98.195 | 2.087 |
| Total | | | | 97229 | 100.000 | |

**2-fluoro-3-(prop-2-yn-1-yl)-3,4-dihydrobenzo[e][1,3,2]oxazaphosphinine 2-oxide (RJG-2273)**

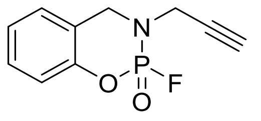


Prepared according to published methods.[7] White solid (86%, 400 mg) **$^1$H NMR** (600 MHz, cdcl$_3$) δ 7.30 (t, *J* = 7.5 Hz, 1H), 7.18 – 7.13 (m, 2H), 7.07 (d, *J* = 8.2 Hz, 1H), 4.61 (dd, *J* = 15.0, 4.1 Hz, 1H), 4.34 (ddd, *J* = 21.4, 15.1, 2.0 Hz, 1H), 4.25 (ddd, *J* = 17.9, 7.8, 2.5 Hz, 2H), 3.98 (dddd, *J* = 18.1, 15.8, 4.5, 2.5 Hz, 1H), 2.36 (t, *J* = 2.5 Hz, 1H). **$^{19}$F NMR** (564 MHz, cdcl$_3$) δ -70.33 (d, *J* = 1018.4 Hz). **$^{31}$P NMR** (243 MHz, cdcl$_3$) δ -6.15 (ddt, *J* = 1019.0, 15.2, 7.8 Hz). $^{13}$C NMR (201 MHz, CDCl$_3$) δ 149.77 (d, *J* = 8.1 Hz), 129.41, 127.13, 125.03, 121.17 (d, *J* = 8.3 Hz), 119.17 (d, *J* = 9.5 Hz), 74.06, 48.70, 37.64 (d, *J* = 5.1 Hz). ESI-TOF (HRMS) m/z [M+H]$^+$ 226.0428, Found 226.0476.
**(-)RJG-2273: [α]$_D$$^{25}$** = -26 (0.1 g/mL, CHCl$_3$)
**(+)RJG-2273: [α]$_D$$^{25}$** = +23 (0.1 g/mL, CHCl$_3$)

***Determination of enantiomeric excess***

Enantiomeric excess was determined using CHIRALPAK IH-3 (100x4.6mm 3.0µm) column eluting with 50% B (Mobile Phases: A: supercritical CO$_2$; B: 50% MeCN in IPA) over 4 minutes, 3.0 mL/min, (-)RJG-2273: 1.120 min. (>99% e.e.), (+)RJG-2273: 1.543 min. (>% e.e.).

(- )RJG-2273

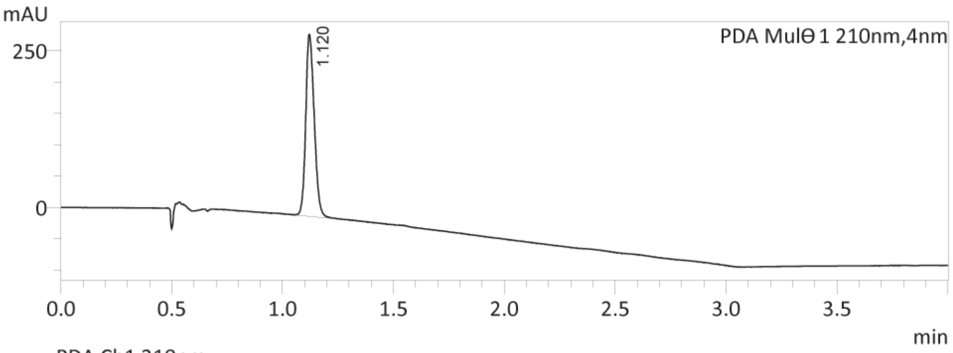


PDA Ch1 210nm

| Peak# | Ret. Time | Height | Width at 50% Height | Area | Area% | Resolution(JP) |
|---|---|---|---|---|---|---|
| 1 | 1.120 | 288527 | 0.041 | 774104 | 99.651 | -- |
| 2 | 1.546 | 1317 | 0.034 | 2714 | 0.349 | 6.697 |
| Total | | | | 776817 | 100.000 | |

(+)RJG-2273

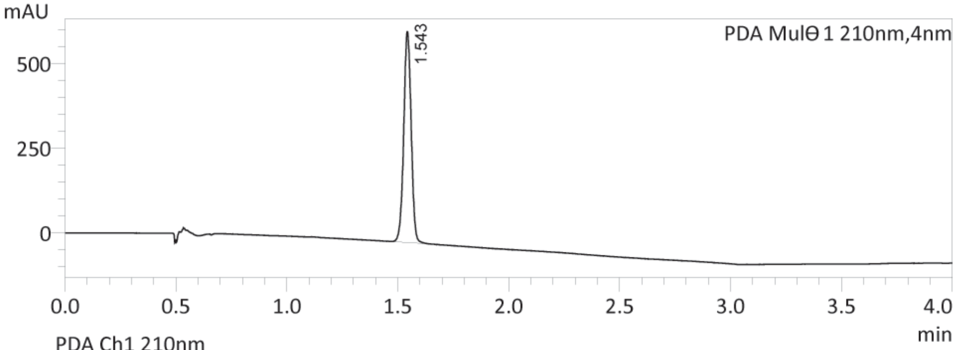


PDA Ch1 210nm

| Peak# | Ret. Time | Height | Width at 50% Height | Area | Area% | Resolution(JP) |
|---|---|---|---|---|---|---|
| 1 | 1.117 | 1024 | 0.042 | 2698 | 0.185 | -- |
| 2 | 1.543 | 620702 | 0.036 | 1455989 | 99.815 | 6.436 |
| Total | | | | 1458688 | 100.000 | |

**3-(prop-2-yn-1-yl)-2-(1*H*-1,2,4-triazol-1-yl)-3,4-dihydrobenzo[e][1,3,2]oxazaphosphinine 2-oxide (RJG-2259A)**

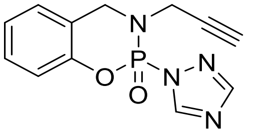


White solid (78%, 88 mg) **$^{1}$H NMR** (600 MHz, cdcl$_3$) δ 8.73 (d, *J* = 0.9 Hz, 1H), 8.04 (d, *J* = 1.9 Hz, 1H), 7.29 (dddt, *J* = 7.6, 5.9, 2.4, 1.3 Hz, 1H), 7.21 – 7.16 (m, 2H), 7.05 (dd, *J* = 8.0, 1.0 Hz, 1H), 4.95 (ddd, *J* = 14.6, 4.0, 1.1 Hz, 1H), 4.32 (dd, *J* = 23.3, 14.6 Hz, 1H), 4.20 (ddd, *J* = 18.0, 6.6, 2.5 Hz, 1H), 3.92 (td, *J* = 17.7, 2.5 Hz, 1H), 2.14 (t, *J* = 2.5 Hz, 1H). **$^{31}$P NMR** (243 MHz, cdcl$_3$) δ -9.53. **$^{13}$C NMR** (201 MHz, cdcl$_3$) δ 154.9 (d, *J* = 17.5 Hz), 149.7 (d, *J* = 8.4 Hz), 149.3 (d, *J* = 10.5 Hz), 129.4, 129.0 (d, *J* = 15.2 Hz), 127.1, 125.2, 124.7 (d, *J* = 3.5 Hz), 121.9 (d, *J* = 7.2 Hz), 118.7 (d, *J* = 9.2 Hz), 76.2 (d, *J* = 4.1 Hz), 74.1, 73.7 (d, *J* = 3.7 Hz), 49.6, 48.5 (d, *J* = 22.4 Hz), 37.5 (d, *J* = 4.7 Hz). ESI-TOF (HRMS) m/z $[M+H]^+$ 275.0692, Found 275.0695.

**(-)RJG-2259A: $[\alpha]_D^{25}$** = -4 (0.1 g/mL, DMF)

**(+)RJG-2259A: $[\alpha]_D^{25}$** = +4 (0.1 g/mL, DMF)

***Determination of enantiomeric excess***

Enantiomeric excess was determined using Amylose-C Neo (100x4.6mm 3.0μm) column eluting with 50% B (Mobile Phases: A: supercritical $CO_2$; B: 50% MeCN in IPA) over 4 minutes, 3.0 mL/min, (-)RJG-2259A: 1.739 min. (>99% e.e.), (+)RJG-2259A: 2.045 min. (>99% e.e.).

(- )RJG-2259A

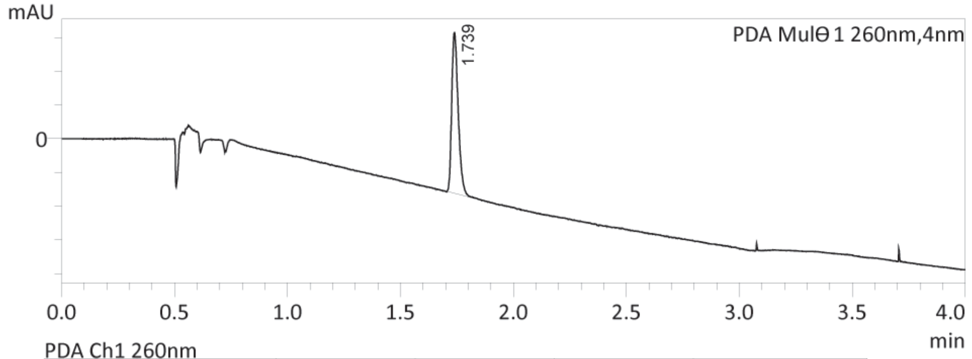


PDA Ch1 260nm

| Peak# | Ret. Time | Height | Area | Area% | Resolution(JP) |
|---|---|---|---|---|---|
| 1 | 1.739 | 94894 | 189753 | 100.000 | -- |
| Total | | | 189753 | 100.000 | |

(+)RJG-2259A

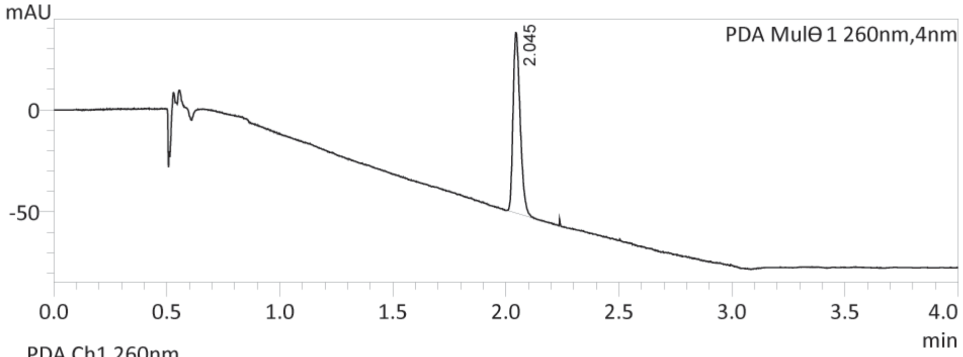


PDA Ch1 260nm

| Peak# | Ret. Time | Height | Area | Area% | Resolution(JP) |
|---|---|---|---|---|---|
| 1 | 2.045 | 87542 | 192779 | 100.000 | -- |
| Total | | | 192779 | 100.000 | |

**3-(prop-2-yn-1-yl)-2-(9H-purin-9-yl)-3,4-dihydrobenzo[e][1,3,2]oxazaphosphinine 2-oxide (RJG-3241)**

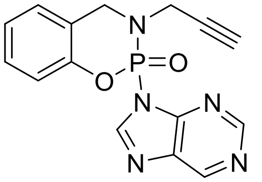

White solid (59%, 320 mg) **$^{1}$H NMR** (500 MHz, cdcl$_3$) δ 9.17 (d, *J* = 1.5 Hz, 1H), 8.89 (s, 1H), 8.56 (s, 1H), 7.34 – 7.29 (m, 1H), 7.26 – 7.20 (m, 2H), 7.09 (d, *J* = 8.2 Hz, 1H), 5.41 (dd, *J* = 14.4, 4.0 Hz, 1H), 4.33 – 4.22 (m, 2H), 3.96 (ddd, *J* = 18.0, 15.5, 2.4 Hz, 1H), 1.96 (t, *J* = 2.5 Hz, 1H). **$^{31}$P NMR** (202 MHz, cdcl$_3$) δ -9.63 – -9.99 (m). **$^{13}$C NMR** (201 MHz, cdcl$_3$) δ 153.47, 153.39, 149.74 (d, *J* = 8.4 Hz), 149.00 (d, *J* = 3.2 Hz), 147.35, 135.26 (d, *J* = 11.3 Hz), 129.37, 127.01, 125.36, 123.55 (d, *J* = 7.2 Hz), 118.75 (d, *J* = 8.3 Hz), 73.79, 49.80 (d, *J* = 2.5 Hz), 37.81 (d, *J* = 6.0 Hz). ESI-TOF (HRMS) m/z $[M+H]^+$ 326.0801, Found 326.0796.

**(-)RJG-3241: $[\alpha]_D^{25}$** = -212(0.2 g/mL, $CHCl_3$)

**(+)RJG-3241: $[\alpha]_D^{25}$** = +213 (0.2 g/mL, $CHCl_3$)

***Determination of enantiomeric excess***

Enantiomeric excess was determined using CHIRALPAK IC-3 (50*4.6mm, 3µm IC30CC-SC002) column eluting with 60% B (Mobile Phases: A: *n*-hexane (0.1% DEA); B: 50% $CH_2Cl_2$ in IPA) over 4 minutes, 1.0 mL/min, (-)RJG-3241: 1.311 min. (>99% e.e.), (+)RJG-3241: 1.632 min. (>99% e.e.).

(-)RJG-3241

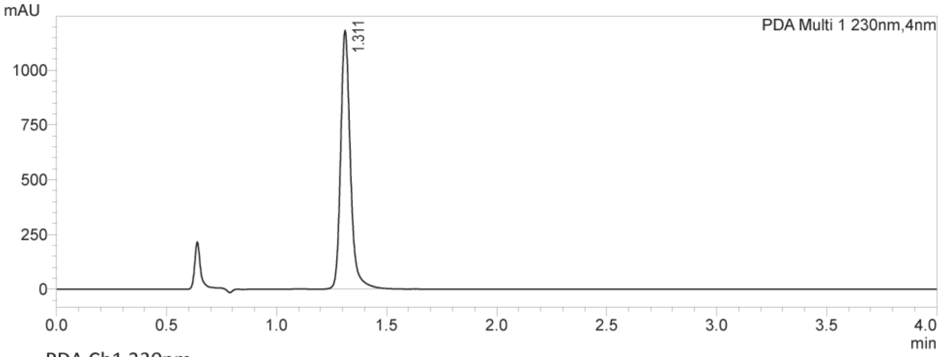


PDA Ch1 230nm

| Peak# | Ret. Time | Height | Area | Area% | Width at 50% Height | Resolution(USP) |
|---|---|---|---|---|---|---|
| 1 | 1.311 | 1155892 | 3618265 | 100.000 | 0.046 | -- |
| Total | | | 3618265 | 100.000 | | |

(+)RJG-3241

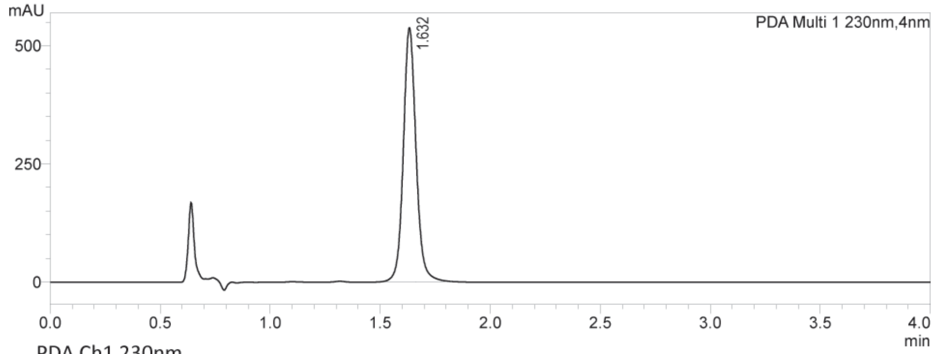


PDA Ch1 230nm

| Peak# | Ret. Time | Height | Area | Area% | Width at 50% Height | Resolution(USP) |
|---|---|---|---|---|---|---|
| 1 | 1.632 | 528818 | 2157967 | 100.000 | 0.060 | -- |
| Total | | | 2157967 | 100.000 | | |

**3-cyclopropyl-2-(1H-1,2,4-triazol-1-yl)-3,4-dihydrobenzo[e][1,3,2]oxazaphosphinine 2-oxide (RJG-3130-1)**

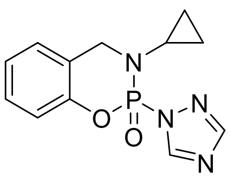

White solid (98%, 222 mg). **$^{1}$H NMR** (400 MHz, $CDCl_3$) δ 8.78 (d, *J* = 0.9 Hz, 1H), 8.06 (d, *J* = 1.9 Hz, 1H), 7.30 – 7.24 (m, 2H), 7.20 – 7.13 (m, 2H), 7.01 (dd, *J* = 8.1, 1.1 Hz, 1H), 4.82 (ddd, *J* = 14.6, 6.3, 1.1 Hz, 1H), 4.39 (dd, *J* = 21.3, 14.6 Hz, 1H), 2.34 (ttd, *J* = 6.8, 3.6, 1.4 Hz, 1H), 1.19 (dddd, *J* = 10.4, 6.7, 5.5, 3.6 Hz, 1H), 0.77 (dtd, *J* = 9.8, 6.8, 5.5 Hz, 1H), 0.67 (dtdd, *J* = 9.8, 6.7, 5.1, 1.6 Hz, 1H), 0.54 – 0.46 (m, 1H). **$^{31}$P NMR** (162 MHz, $CDCl_3$) δ -9.18 (dd, *J* = 21.9, 6.3 Hz). **$^{13}$C NMR** (201 MHz, $CDCl_3$) δ 129.3, 126.9, 125.1, 122.7 (d, *J* = 6.6 Hz), 118.5 (d, *J* = 8.2 Hz), 51.7 (d, *J* = 2.9 Hz), 29.6 (d, *J* = 4.0 Hz), 7.9, 5.9 (d, *J* = 6.6 Hz). ESI-TOF (HRMS) m/z $[M+H]^+$ 277.0849, Found 277.0856.

**3-(cyclopropylmethyl)-2-(1H-1,2,4-triazol-1-yl)-3,4-dihydrobenzo[e][1,3,2]oxazaphosphinine 2-oxide (RJG-3045)**

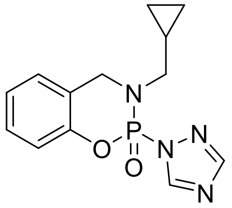

Off-white solid (80%, 95 mg) **$^{1}$H NMR** (400 MHz, $CDCl_3$) δ 8.72 (d, *J* = 0.9 Hz, 1H), 8.03 (d, *J* = 1.9 Hz, 1H), 7.29 – 7.23 (m, 1H), 7.20 – 7.12 (m, 2H), 7.01 (dd, *J* = 8.2, 1.1 Hz, 1H), 4.95 (ddt, *J* = 14.7, 4.7, 0.9 Hz, 1H), 4.38 (dd, *J* = 22.7, 14.7 Hz, 1H), 3.20 – 3.03 (m, 2H), 0.85 (dddd, *J* = 12.6, 7.7, 6.0, 2.9 Hz, 1H), 0.49 (dddd, *J* = 9.1, 8.1, 5.7, 4.6 Hz, 1H), 0.43 – 0.34 (m, 1H), 0.20 (ddt, *J* = 9.5, 5.6, 4.7 Hz, 1H), 0.06 (ddt, *J* = 9.5, 5.7, 4.7 Hz, 1H). **$^{31}$P NMR** (162 MHz, $CDCl_3$) δ -8.53 – -9.00 (m). **$^{13}$C NMR** (201 MHz, $CDCl_3$) δ 154.8 (d, *J* = 17.2 Hz), 149.7 (d, *J* = 8.2 Hz), 149.3 (d, *J* = 10.2 Hz), 129.2, 126.9, 124.9, 122.6 (d, *J* = 7.5 Hz), 118.6 (d, *J* = 8.7 Hz), 53.3 (d, *J* = 3.6 Hz), 49.7, 9.2 (d, *J* = 2.9 Hz), 4.2, 3.3. ESI-TOF (HRMS) m/z $[M+H]^+$ 291.1005, Found 291.1012.

**3-cyclopropyl-2-(1H-imidazol-1-yl)-3,4-dihydrobenzo[e][1,3,2]oxazaphosphinine 2-oxide (RJG-3130-3)**

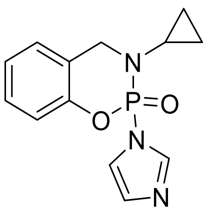

White solid (50%, 114 mg) **$^{1}$H NMR** (500 MHz, $CDCl_3$) δ 7.80 (t, *J* = 1.1 Hz, 1H), 7.33 – 7.28 (m, 1H), 7.23 – 7.16 (m, 2H), 7.10 (ddd, *J* = 2.5, 1.6, 0.9 Hz, 1H), 7.07 – 7.02 (m, 1H), 6.97 (q, *J* = 1.5 Hz, 1H), 4.55 (dd, *J* = 14.7, 12.7 Hz, 1H), 4.31 (dd, *J* = 16.4, 14.7 Hz, 1H), 2.48 (ttd, *J* = 6.9, 3.7, 1.5 Hz, 1H), 0.94 (dddd, *J* = 10.3, 6.6, 5.2, 3.6 Hz, 1H), 0.79 – 0.64 (m, 3H), 0.42 (dddd, *J* = 10.5, 6.6, 5.1, 3.6 Hz, 1H). **$^{31}$P NMR** (162 MHz, $CDCl_3$) δ -6.67 (t, *J* = 14.6 Hz). **$^{13}$C NMR** (201 MHz, $CDCl_3$) δ 149.4 (d, *J* = 8.7 Hz), 140.0 (d, *J* = 5.5 Hz), 131.6 (d, *J* = 13.2 Hz), 129.8, 126.9, 125.4, 123.8 (d, *J* = 6.3 Hz), 119.0 (dd, *J* = 7.1, 5.3 Hz), 50.9 (d, *J* = 3.7 Hz), 29.8 (d, *J* = 4.6 Hz), 7.1 (d, *J* = 4.6 Hz), 6.4 (d, *J* = 3.8 Hz). ESI-TOF (HRMS) m/z $[M+H]^+$ 276.0896, Found 276.0893.

**3-cyclopropyl-2-(1H-pyrazol-1-yl)-3,4-dihydrobenzo[e][1,3,2]oxazaphosphinine 2-oxide (RJG-3130-4)**

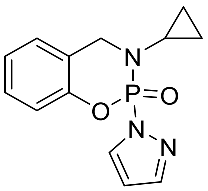

White solid (96%, 217 mg) **$^{1}$H NMR** (500 MHz, $CDCl_3$) δ 8.16 (d, *J* = 2.6 Hz, 1H), 7.74 (dd, *J* = 3.1, 1.6 Hz, 1H), 7.25 – 7.21 (m, 1H), 7.16 (ddd, *J* = 7.7, 1.9, 1.0 Hz, 1H), 7.12 (td, *J* = 7.5, 1.2 Hz, 1H), 7.00 (dd, *J* = 8.2, 1.1 Hz, 1H), 6.40 (td, *J* = 2.8, 1.5 Hz, 1H), 4.85 (dd, *J* = 14.5, 6.5 Hz, 1H), 4.35 (dd, *J* = 21.1, 14.5 Hz, 1H), 2.36 (ttd, *J* = 6.9, 3.6, 1.4 Hz, 1H), 1.15 (dddd, *J* = 10.5, 6.7, 5.4, 3.6 Hz, 1H), 0.72 (dtd, *J* = 9.7, 6.8, 5.4 Hz, 1H), 0.62 (dtdd, *J* = 9.7, 6.7, 5.0, 1.6 Hz, 1H), 0.52 – 0.42 (m, 1H). **$^{31}$P NMR** (162 MHz, $CDCl_3$) δ -6.56 (d, *J* = 21.3 Hz). **$^{13}$C NMR** (201 MHz, $CDCl_3$) δ 149.8 (d, *J* = 8.3 Hz), 145.4 (d, *J* = 14.9 Hz), 135.8 (d, *J* = 11.5 Hz), 129.0, 126.8, 124.6, 123.3 (d, *J* = 6.4 Hz), 118.6 (d, *J* = 8.2 Hz), 107.7 (d, *J* = 6.8 Hz), 51.8 (d, *J* = 2.5 Hz), 29.6 (d, *J* = 3.9 Hz), 7.7 (d, *J* = 2.2 Hz), 5.8 (d, *J* = 6.5 Hz). ESI-TOF (HRMS) m/z $[M+H]^+$ 276.0896, Found 276.0898.

**2-(1*H*-benzo[*d*]28yridine28-1-yl)-3-cyclopropyl-3,4-dihydrobenzo[e][1,3,2]oxazaphosphinine 2-oxide (RJG-3130-7)**

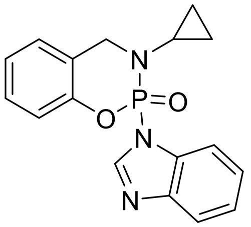

White solid (94%, 317 mg) $^{1}$**H NMR** (400 MHz, $CDCl_3$) δ 8.27 (s, 1H), 7.83 (d, *J* = 8.0 Hz, 1H), 7.41 – 7.33 (m, 2H), 7.30 (ddd, *J* = 14.3, 4.8, 2.6 Hz, 1H), 7.25 – 7.20 (m, 2H), 7.06 – 7.01 (m, 1H), 4.64 (dd, *J* = 14.8, 11.8 Hz, 1H), 4.46 (dd, *J* = 17.4, 14.8 Hz, 1H), 2.51 (ddq, *J* = 6.7, 5.4, 3.5 Hz, 1H), 0.92 (dq, *J* = 10.5, 5.2 Hz, 1H), 0.73 – 0.55 (m, 2H), 0.36 – 0.24 (m, 1H). $^{31}$**P NMR** (162 MHz, $CDCl_3$) δ -6.63 (dd, *J* = 17.8, 11.9 Hz). $^{13}$**C NMR** (126 MHz, $CDCl_3$) δ 149.60 (d, *J* = 8.5 Hz), 145.18, 145.06 (d, *J* = 6.7 Hz), 133.51 (d, *J* = 5.4 Hz), 129.88, 127.05, 125.32, 124.78, 124.09, 123.34 (d, *J* = 6.3 Hz), 113.03, 51.11 (d, *J* = 3.9 Hz), 32.02, 29.92 (d, *J* = 4.2 Hz), 29.16, 22.83, 14.26, 6.81 (d, *J* = 5.7 Hz), 6.29 (d, *J* = 2.8 Hz. ESI-TOF (HRMS) m/z $[M+H]^+$ 326.1053, Found 326.1047.

**3-cyclopropyl-2-(3H-imidazo[4,5-b]28yridine-3-yl)-3,4-dihydrobenzo[e][1,3,2]oxazaphosphinine 2-oxide (RJG-3130-8)**

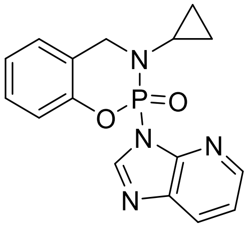

White solid (9%, 24 mg) $^{1}$**H NMR** (500 MHz, $CDCl_3$) δ 8.59 (s, 1H), 8.29 (dd, *J* = 4.8, 1.5 Hz, 1H), 8.10 (dt, *J* = 8.0, 1.6 Hz, 1H), 7.28 (dd, *J* = 4.5, 3.5 Hz, 1H), 7.23 (dd, *J* = 8.1, 1.7 Hz, 2H), 7.17 (td, *J* = 7.4, 1.1 Hz, 1H), 7.05 (dd, *J* = 8.1, 1.2 Hz, 1H), 5.45 (dd, *J* = 14.2, 4.5 Hz, 1H), 4.28 (dd, *J* = 24.4, 14.1 Hz, 1H), 4.13 (q, *J* = 7.1 Hz, 0H), 2.42 (ttd, *J* = 7.0, 3.7, 1.2 Hz, 1H), 2.05 (s, 0H), 1.27 (t, *J* = 7.1 Hz, 1H), 1.23 – 1.17 (m, 1H), 0.70 – 0.60 (m, 2H), 0.51 (dddd, *J* = 12.3, 6.2, 4.2, 1.2 Hz, 1H). $^{31}$**P NMR** (162 MHz, $CDCl_3$) δ -8.60 (d, *J* = 24.4 Hz). $^{13}$**C NMR** (201 MHz, $CDCl_3$) δ 149.80 (d, *J* = 7.9 Hz), 148.26 (d, *J* = 3.6 Hz), 146.95 (d, *J* = 5.8 Hz), 145.33, 136.80 (d, *J* = 11.7 Hz), 128.97, 128.54, 126.76, 124.84, 124.79, 119.78, 118.72 (d, *J* = 7.6 Hz), 51.62 (d, *J* = 4.7 Hz), 32.03, 29.70 (d, *J* = 5.4 Hz), 29.17, 22.84, 14.26, 7.46 (d, *J* = 2.2 Hz), 6.06 (d, *J* = 5.9 Hz. ESI-TOF (HRMS) m/z $[M+H]^+$ 327.1005, Found 327.1000.

**3-cyclopropyl-2-(9*H*-purin-9-yl)-3,4-dihydrobenzo[*e*][1,3,2]oxazaphosphinine 2-oxide (RJG-3130-9)**

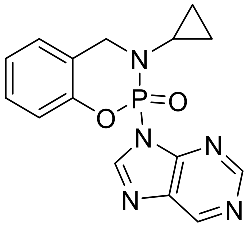

White solid (60%, 190 mg) $^{1}$**H NMR** (400 MHz, CDCl3) δ 9.19 (d, *J* = 1.5 Hz, 1H), 8.90 (s, 1H), 8.61 (s, 1H), 7.26 (s, 3H), 7.18 (td, *J* = 7.4, 1.2 Hz, 1H), 7.03 (dd, *J* = 8.0, 1.2 Hz, 1H), 5.37 (dd, *J* = 14.3, 5.0 Hz, 1H), 4.33 (dd, *J* = 23.5, 14.3 Hz, 1H), 2.43 (ttd, *J* = 6.9, 3.6, 1.3 Hz, 1H), 1.30 – 1.14 (m, 1H), 0.76 – 0.63 (m, 2H), 0.56 – 0.44 (m, 1H). $^{31}$**P NMR** (162 MHz, $CDCl_3$) δ -9.65 – -9.96 (m). $^{13}$**C NMR** (126 MHz, $CDCl_3$) δ 153.66, 153.05, 149.50, 149.39, 147.74 (d, *J* = 5.6 Hz), 135.38 (d, *J* = 10.9 Hz), 129.20, 126.85, 125.19, 124.32 (d, *J* = 6.7 Hz), 118.66 (d, *J* = 7.5 Hz), 51.54 (d, *J* = 4.9 Hz), 29.78 (d, *J* = 5.4 Hz), 7.54 (d, *J* = 2.5 Hz), 6.16 (d, *J* = 5.6 Hz. ESI-TOF (HRMS) m/z $[M+H]^+$ 328.0958, Found 328.0956.

**(-)RJG-3130-9: $[\alpha]_D^{25}$** = -94 (0.1 g/mL, $CHCl_3$)

**(+)RJG-3130-9: $[\alpha]_D^{25}$** = +96 (0.1 g/mL, $CHCl_3$)

***Determination of enantiomeric excess***

Enantiomeric excess was determined using a Shimadzu LC-20AD equipped with a CHIRALPAK IK-3 (50x4.6mm 3.0µm, IK30CB-CW002) column eluting with 50% B (Mobile Phases: A: n-hexane; B: 50% $CH_2Cl_2$ in IPA) over 3 minutes, 1.0 mL/min, (-)RJG-3130-9: 1.211 min. (>99% e.e.), (+)RJG-3130-9: 1.396 min. (>99% e.e.).

(-)RJG-3130-9

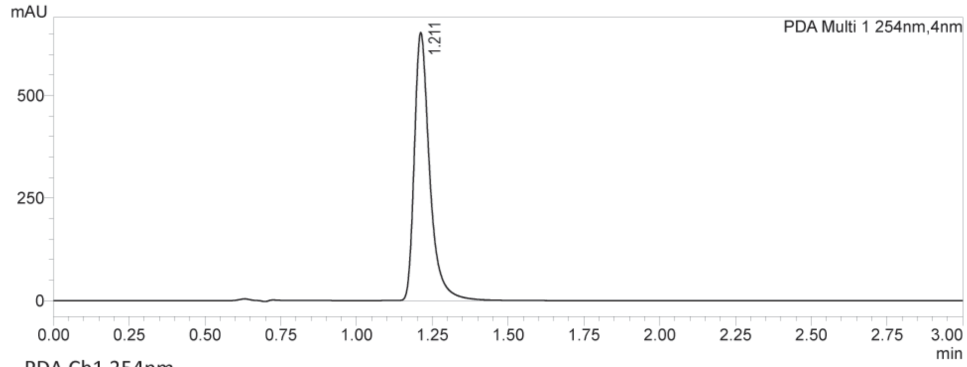


PDA Ch1 254nm

| Peak# | Ret. Time | Height | Area | Area% | Width at 50% Height | Resolution(USP) |
|---|---|---|---|---|---|---|
| 1 | 1.211 | 649068 | 2313154 | 100.000 | 0.051 | -- |
| Total | | | 2313154 | 100.000 | | |

(+)RJG-3130-9

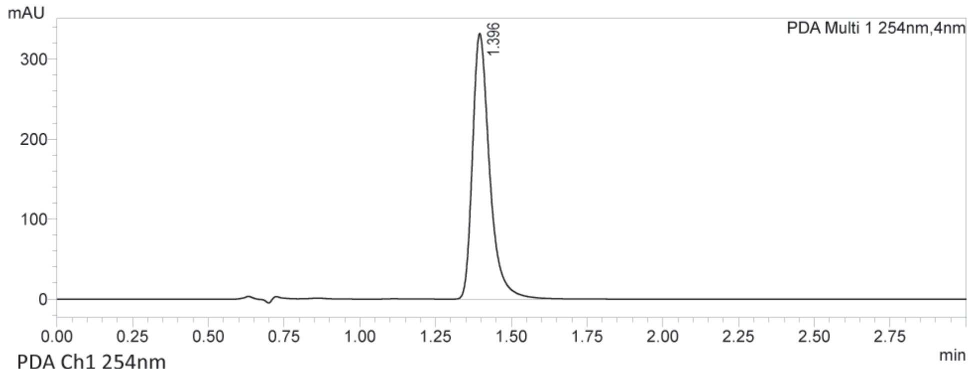


PDA Ch1 254nm

| Peak# | Ret. Time | Height | Area | Area% | Width at 50% Height | Resolution(USP) |
|---|---|---|---|---|---|---|
| 1 | 1.396 | 330396 | 1307828 | 100.000 | 0.058 | -- |
| Total | | | 1307828 | 100.000 | | |

### NMR Spectra

#### RJG-2273 $^{1}$H NMR (600 MHz, cdcl$_3$)

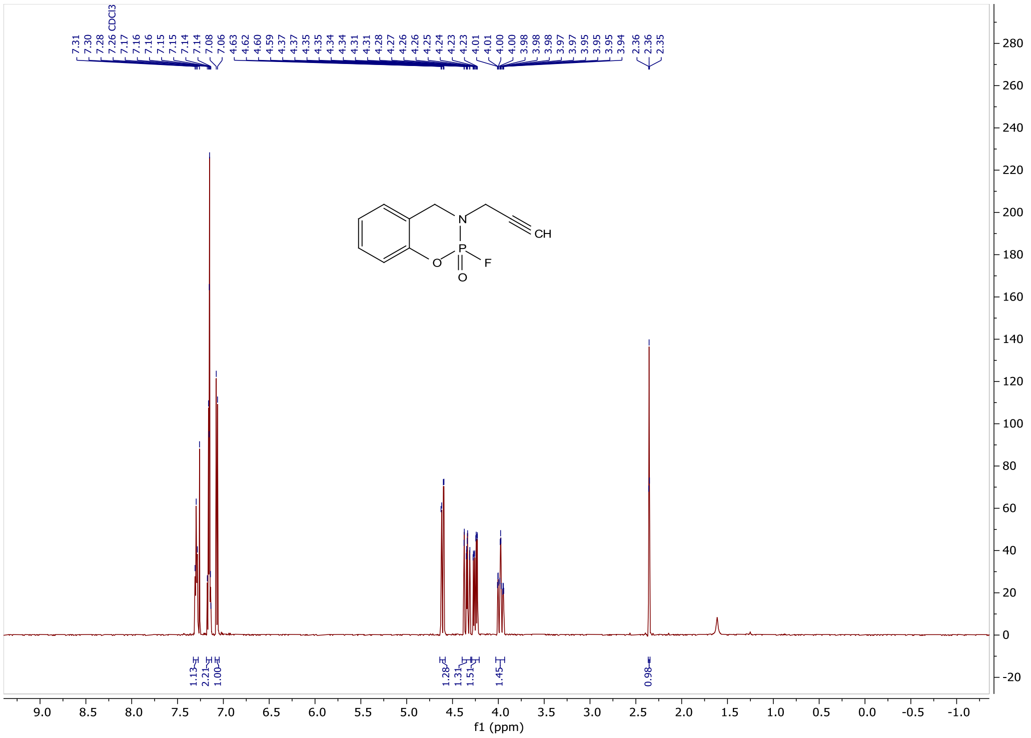

#### RJG-2273 $^{19}$F NMR (564 MHz, cdcl$_3$)

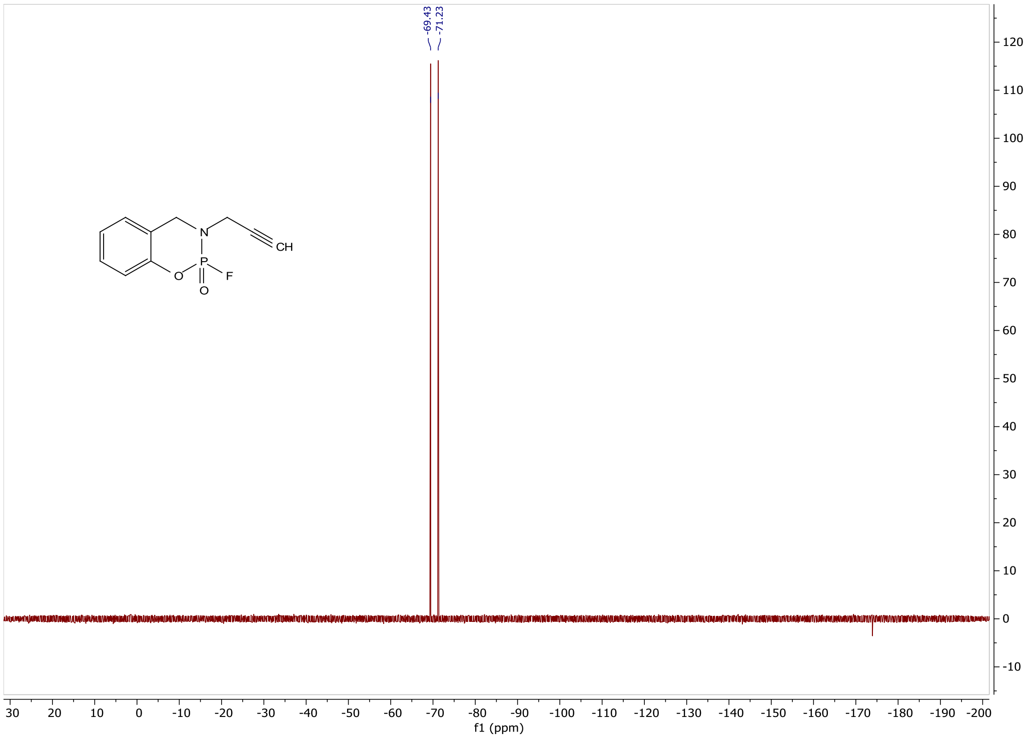

#### RJG-2273 $^{31}P$ NMR (243 MHz, cdcl$_3$)

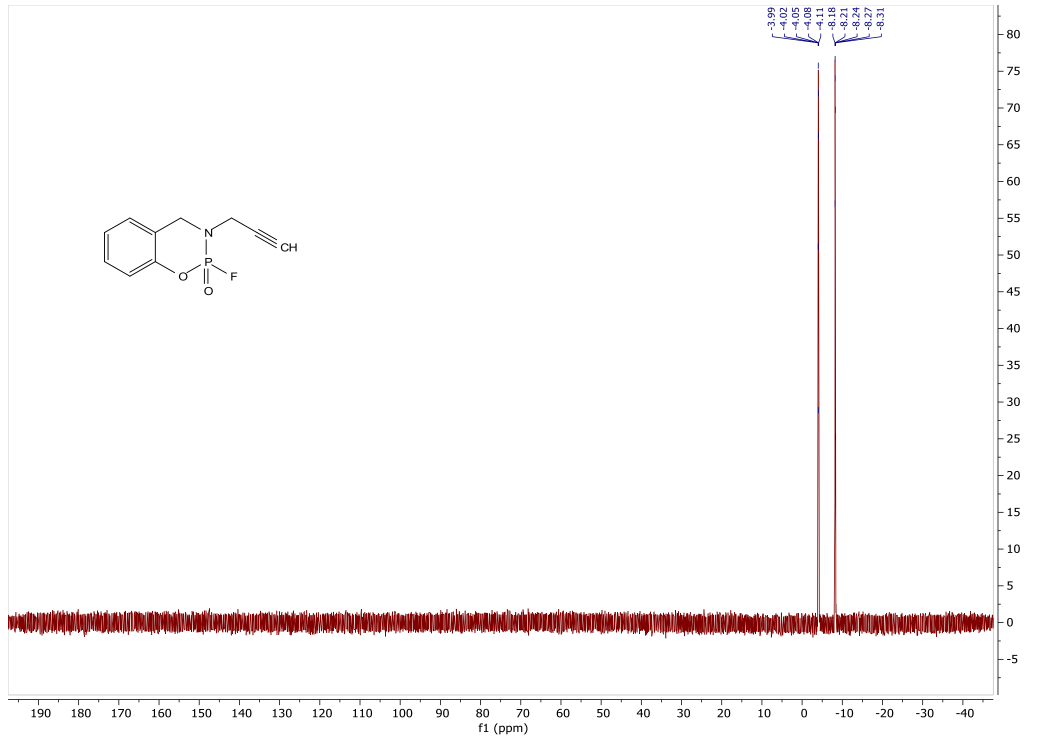

#### RJG-2273 $^{13}C$ NMR (201 MHz, cdcl$_3$)

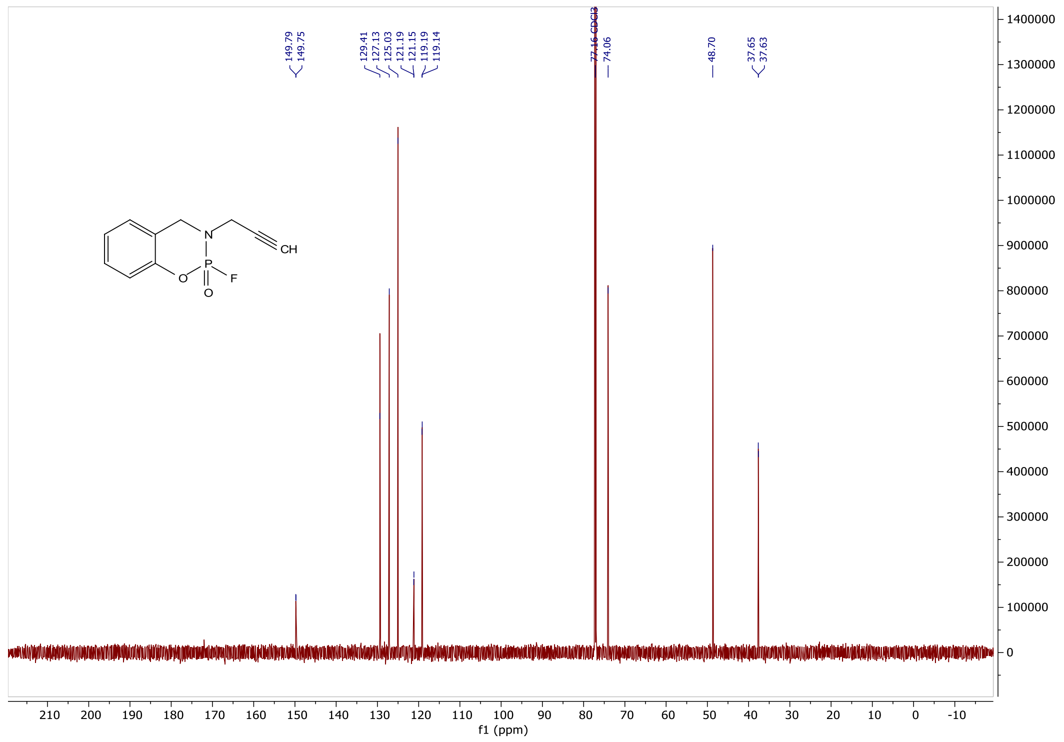

### RJG-2259A $^{1}H$ NMR (600 MHz, cdcl$_3$)

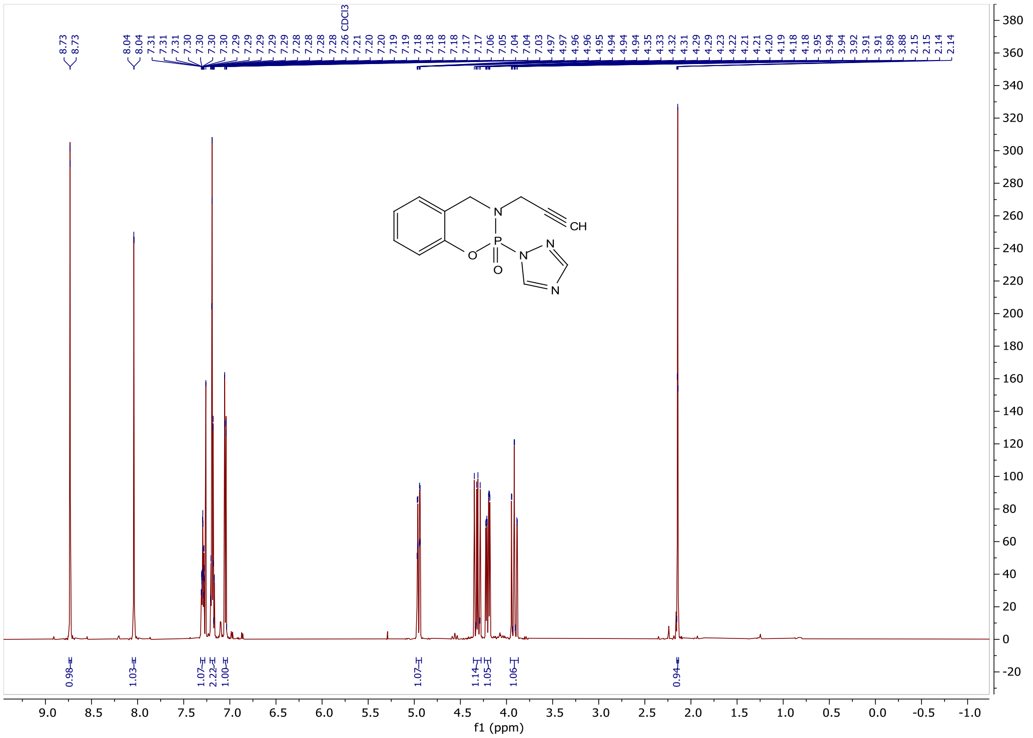


### RJG-2259A $^{31}P$ NMR (243 MHz, cdcl$_3$)

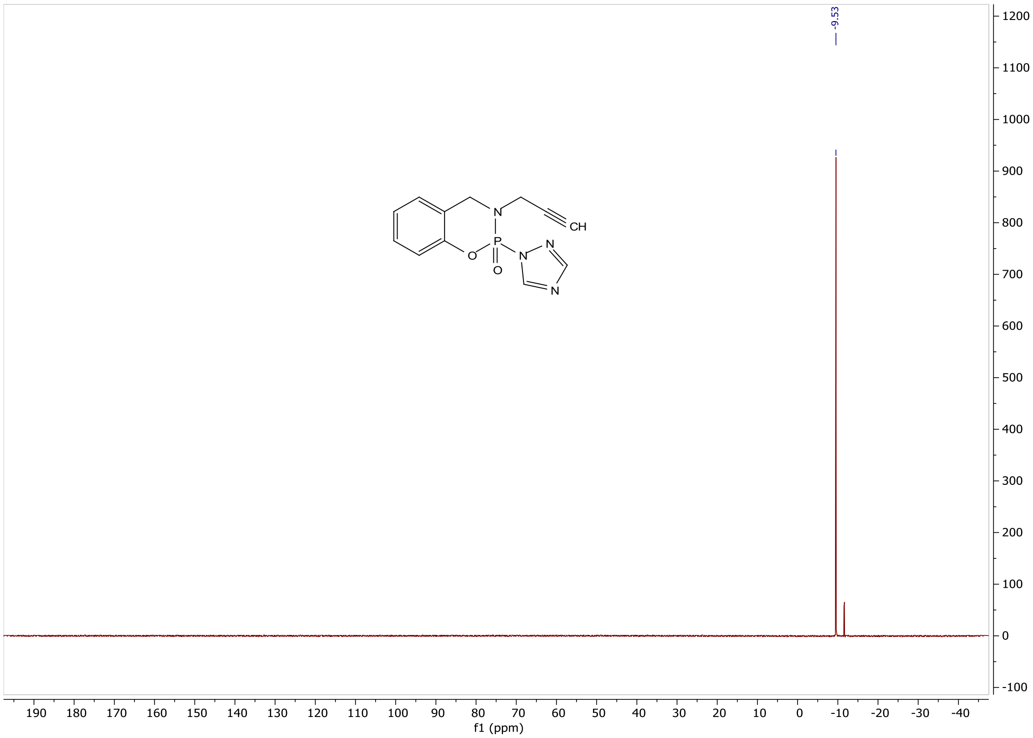

#### RJG-2259A $^{13}C$ NMR (201 MHz, cdcl$_3$)

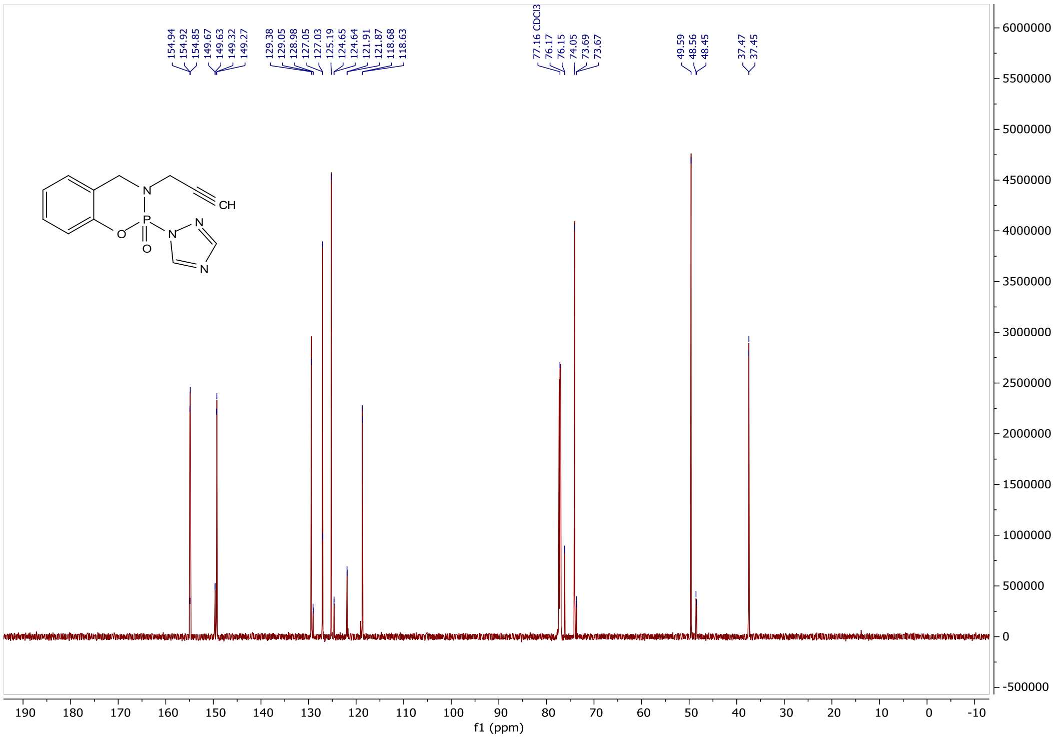

#### RJG-3241 $^{1}H$ NMR (500 MHz, cdcl$_3$)

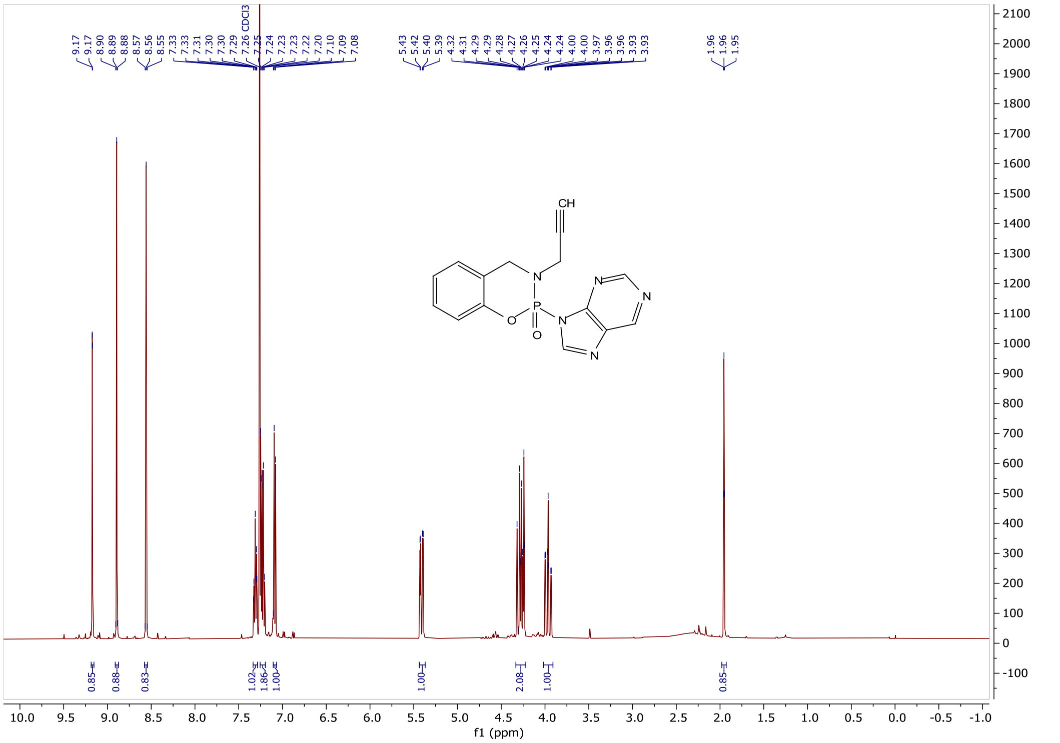

#### RJG-3241 $^{31}P$ NMR (202 MHz, cdcl$_3$)

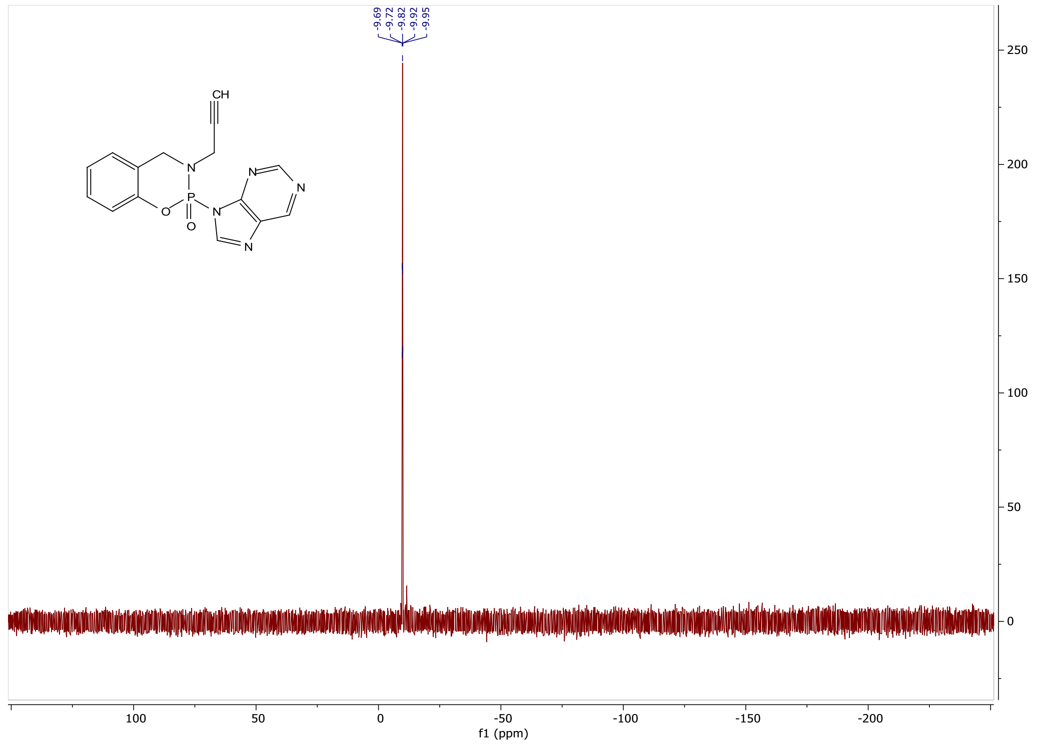

#### RJG-3241 $^{13}C$ NMR (126 MHz, cdcl$_3$)

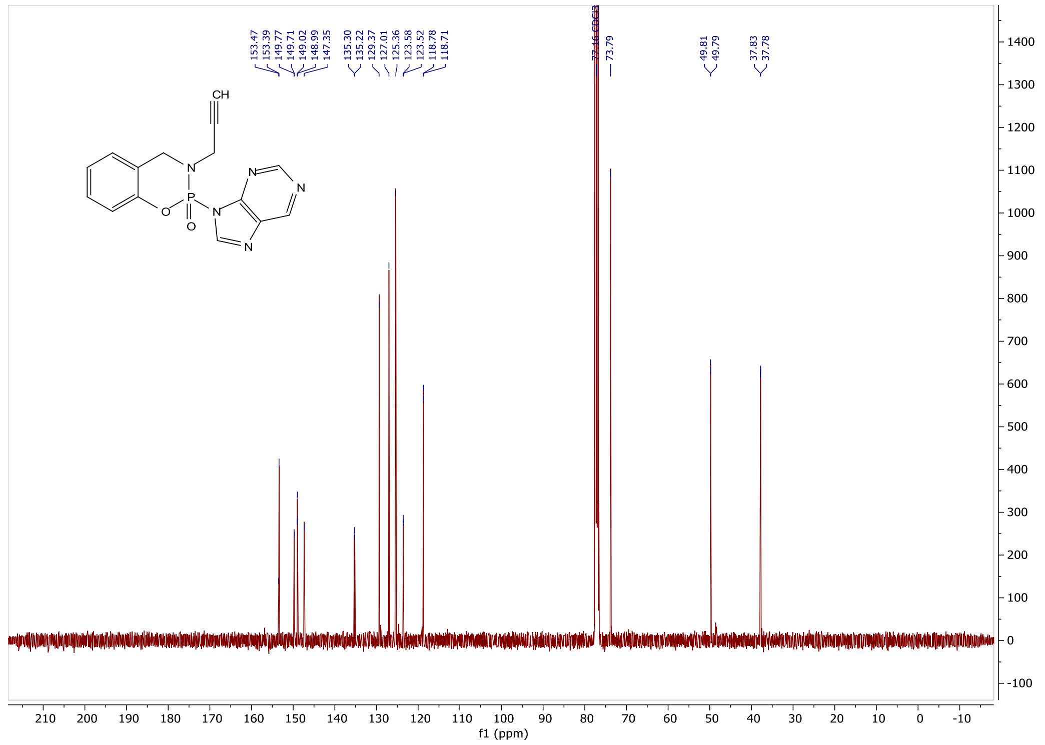

#### RJG-3130-1 $^{1}H$ NMR (400 MHz, cdcl$_3$)

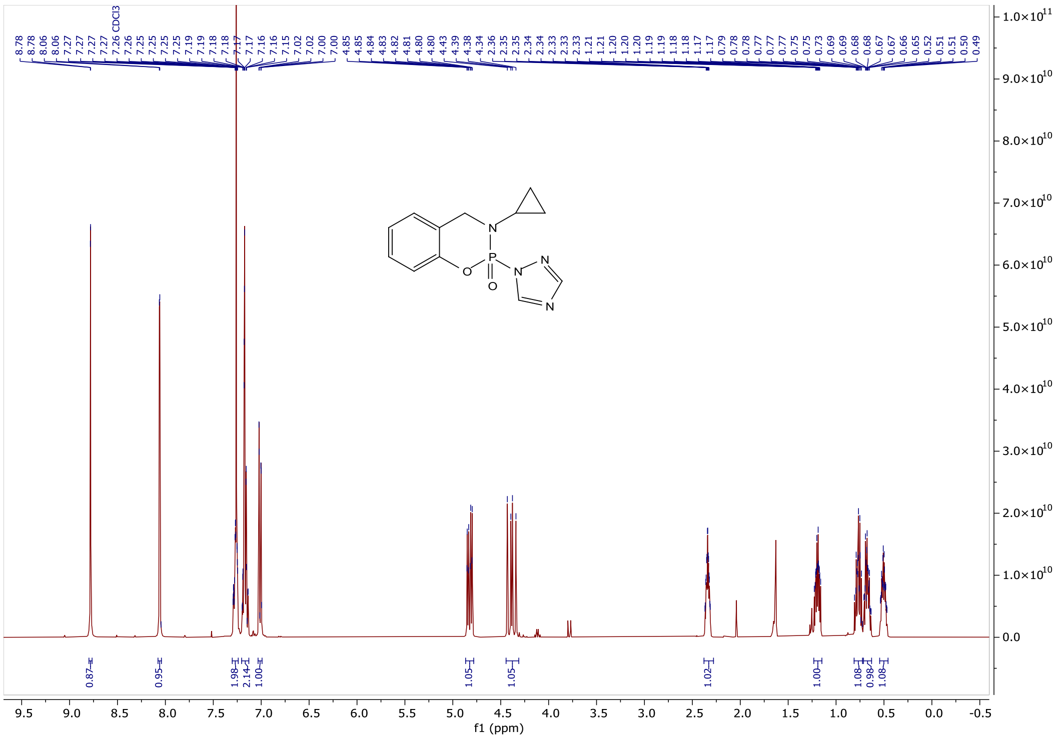

#### RJG-3130-1 $^{31}P$ NMR (162 MHz, cdcl$_3$)

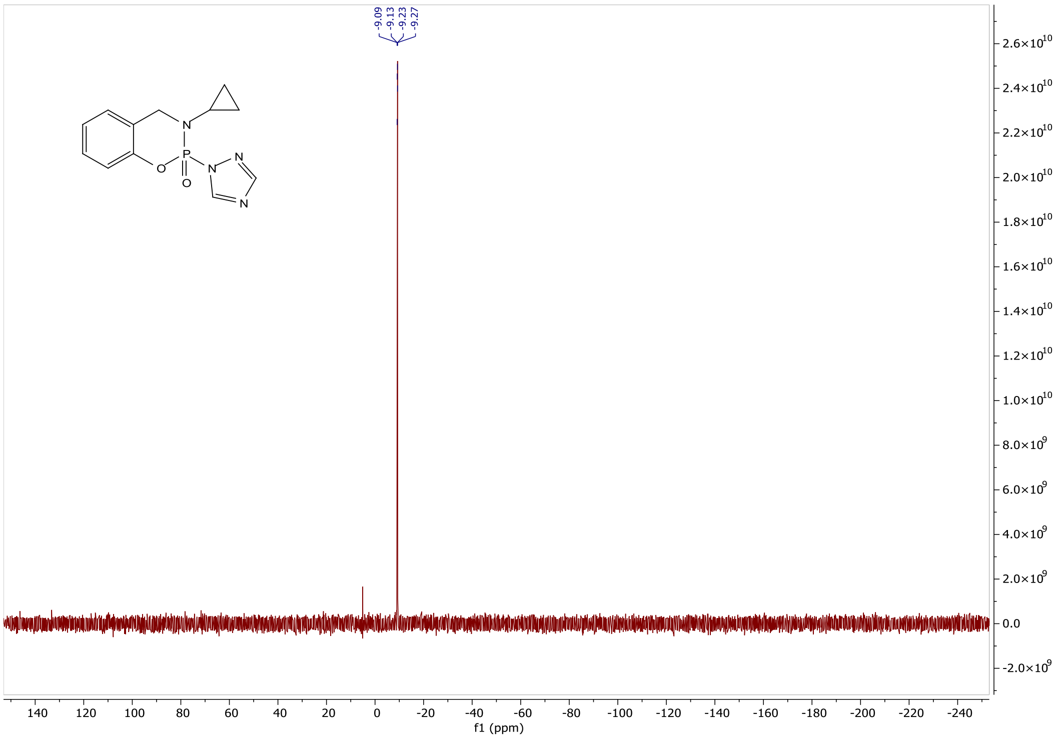

#### RJG-3130-1 $^{13}$C NMR (201 MHz, cdcl$_3$)

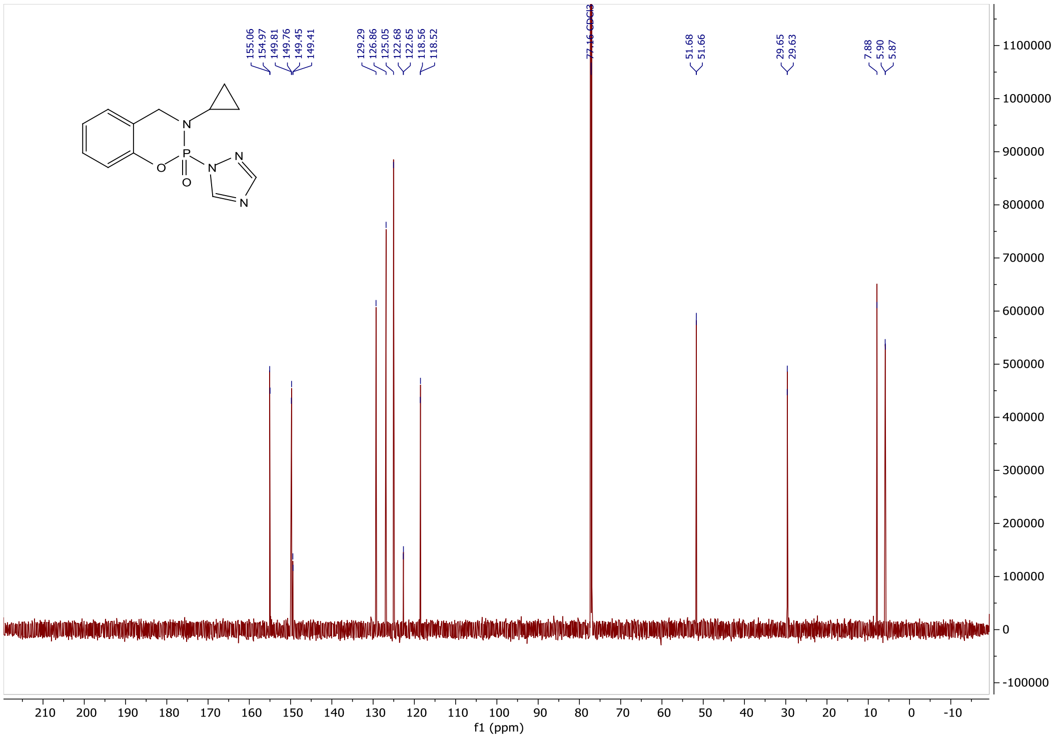


#### RJG-3045 $^{1}$H NMR (400 MHz, cdcl$_3$)

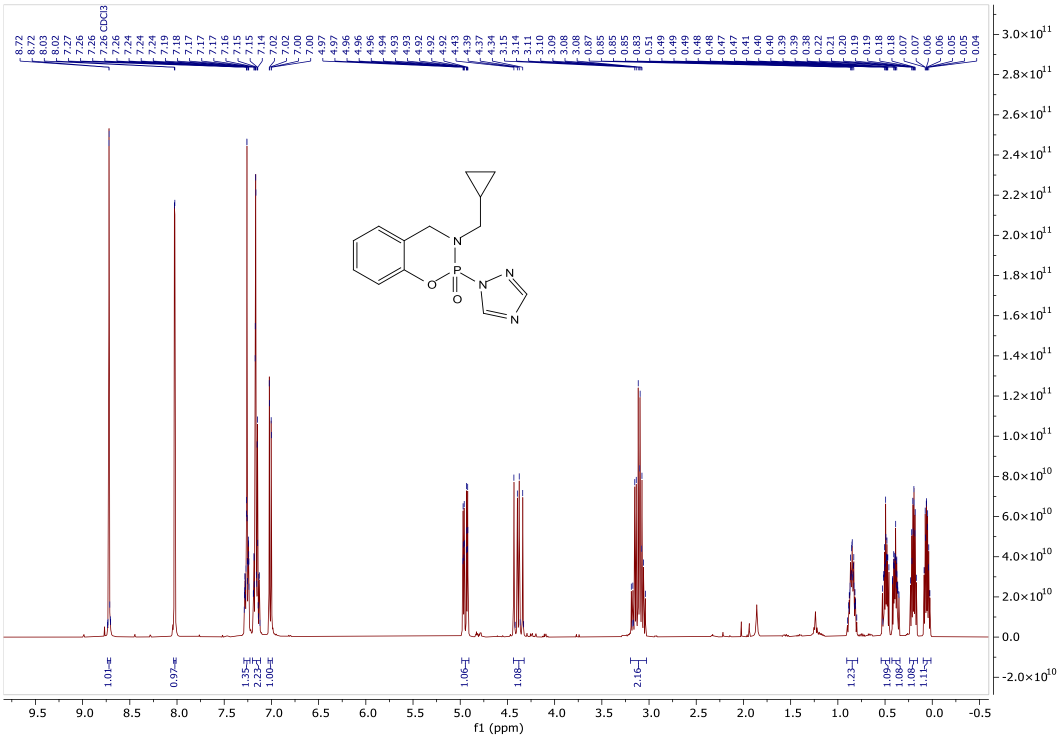

#### RJG-3045 $^{31}$P NMR (162 MHz, cdcl$_3$)

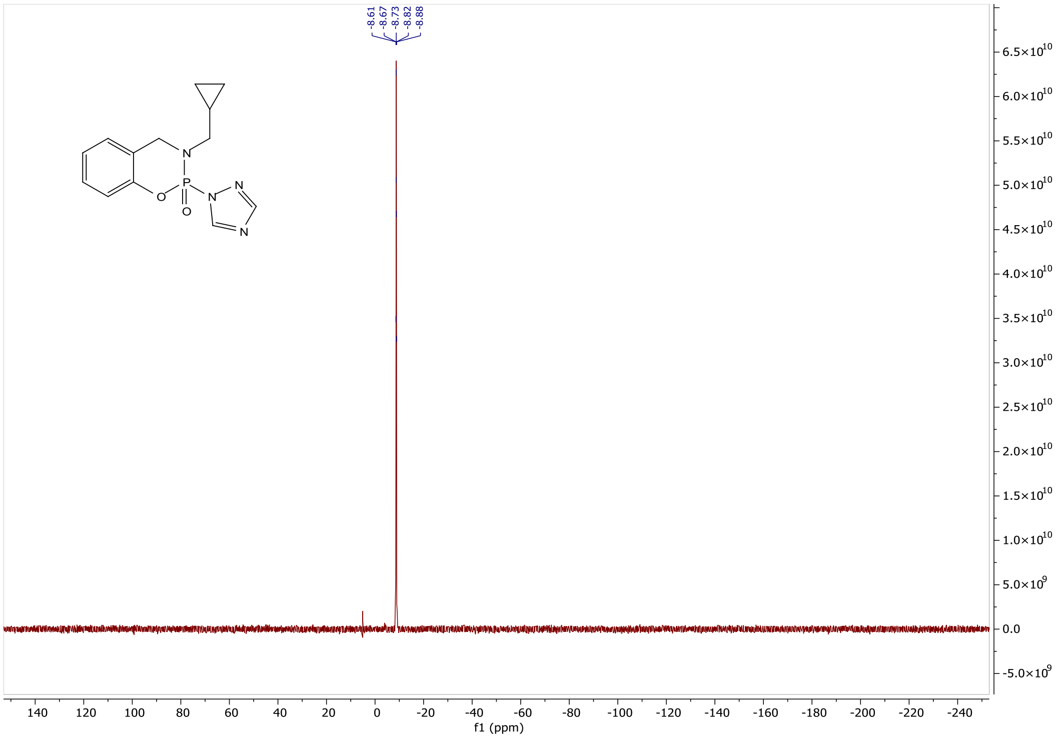


#### RJG-3045 $^{13}$C NMR (201 MHz, cdcl$_3$)

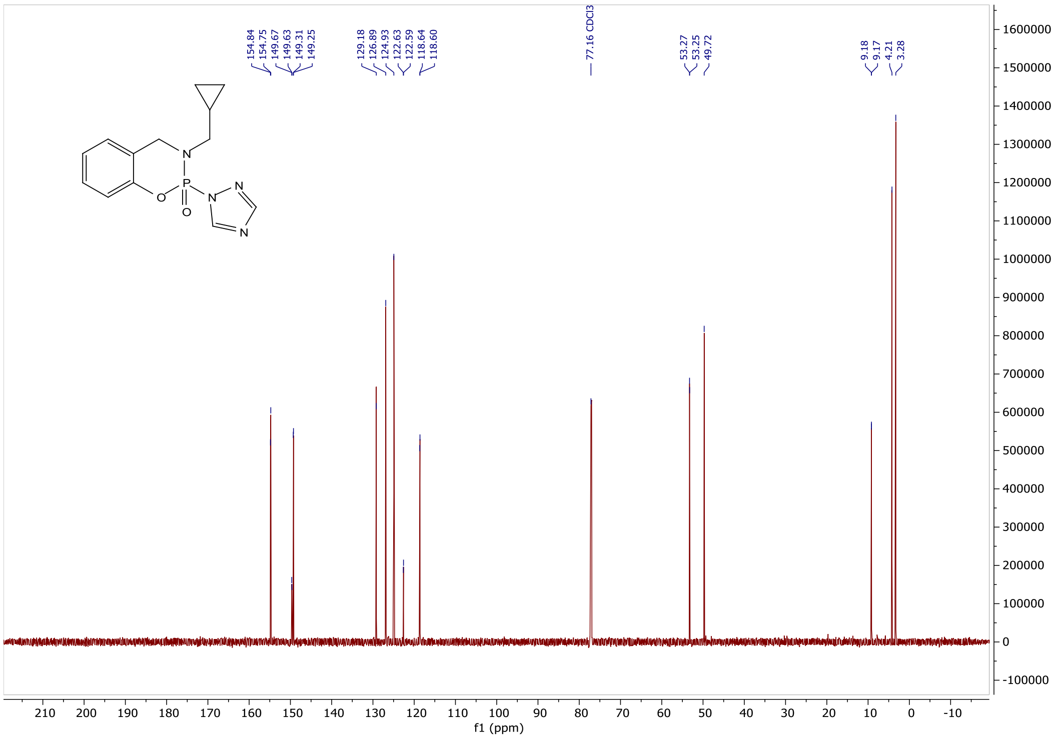

#### RJG-3130-3 $^{1}$H NMR (500 MHz, cdcl$_3$)

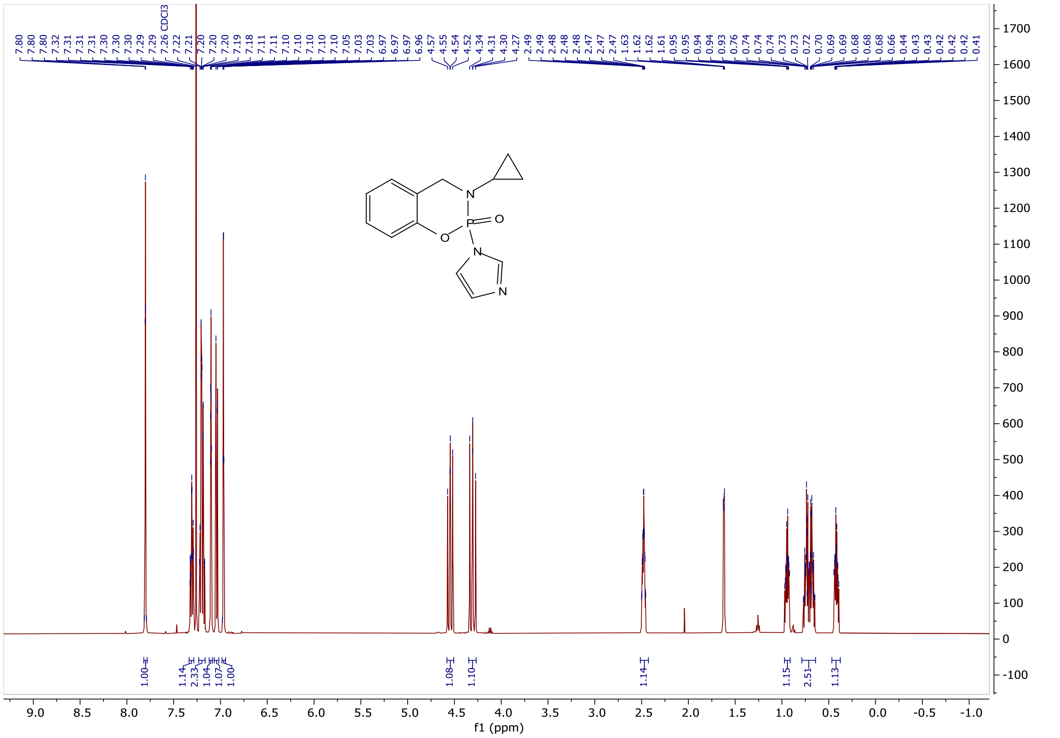

#### RJG-3130-3 $^{31}$P NMR (162 MHz, cdcl$_3$)

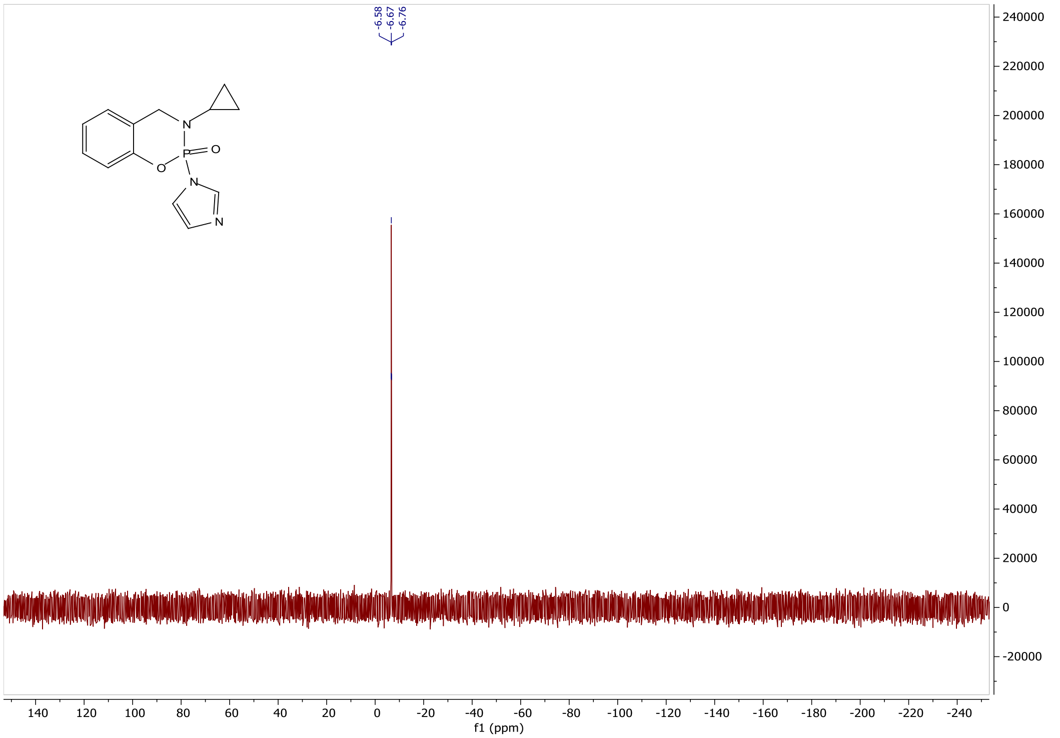

#### RJG-3130-3 $^{13}C$ NMR (126 MHz, cdcl$_3$)

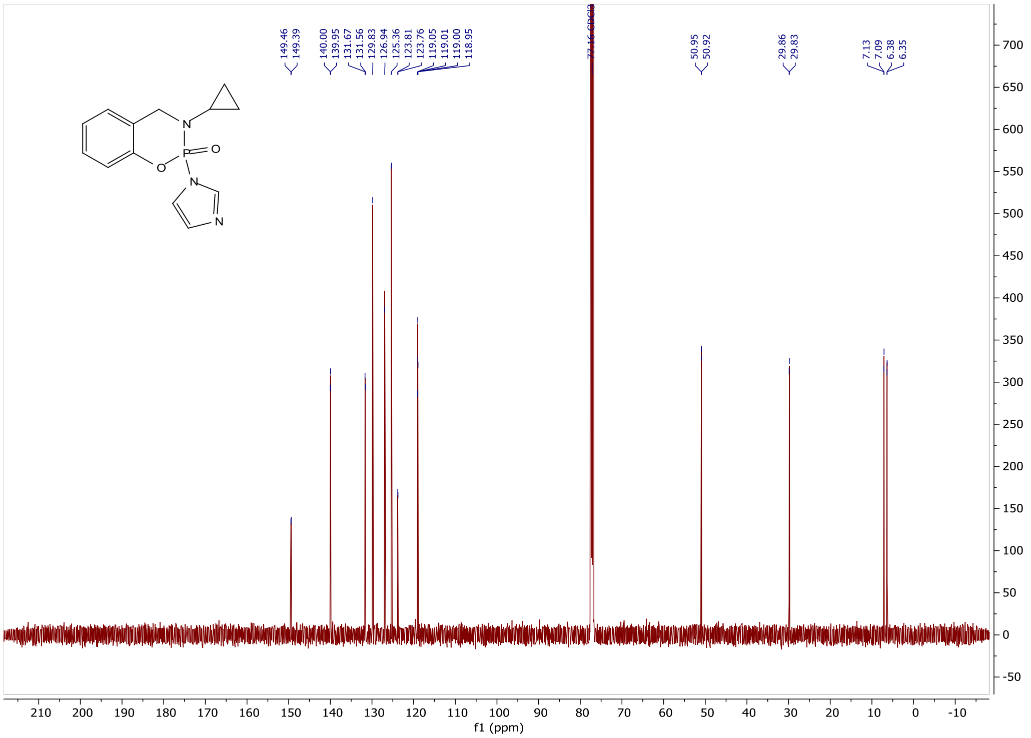

#### RJG-3130-4 $^{1}H$ NMR (500 MHz, cdcl$_3$)

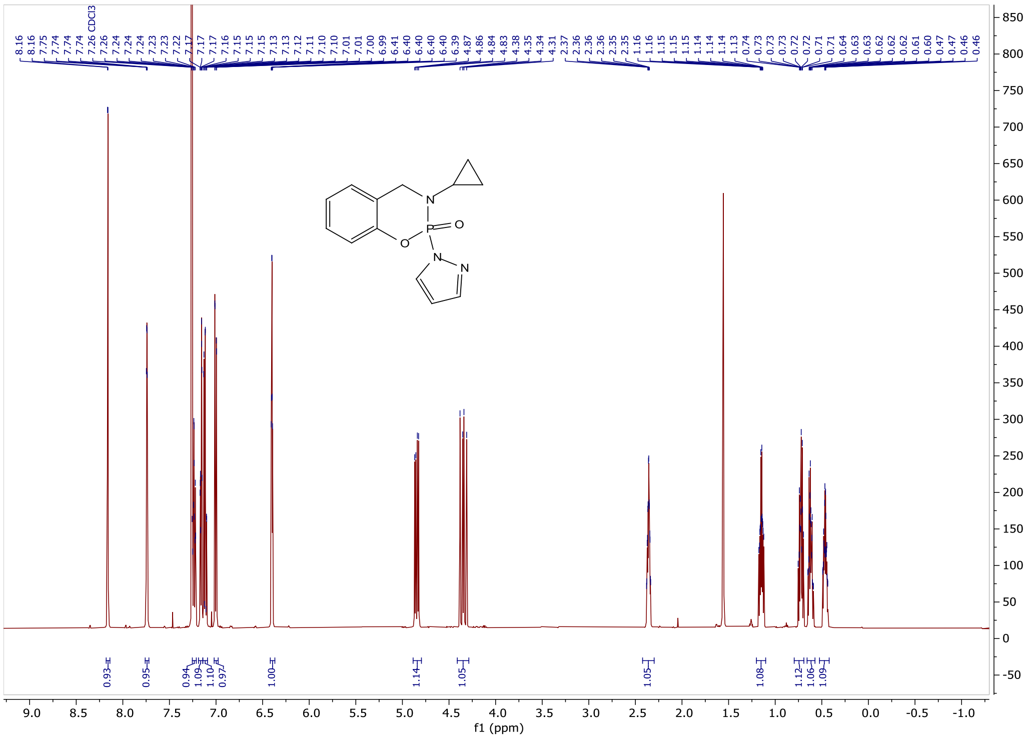

#### RJG-3130-4 $^{31}P$ NMR (162 MHz, cdcl$_3$)

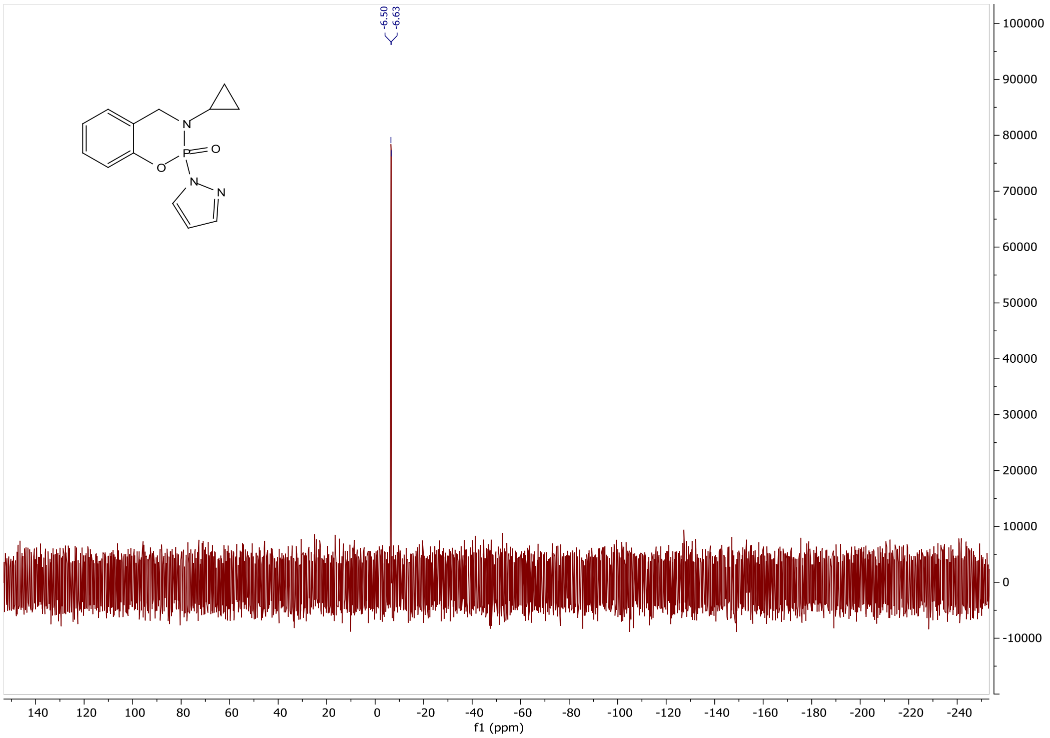


#### RJG-3130-4 $^{13}C$ NMR (126 MHz, cdcl$_3$)

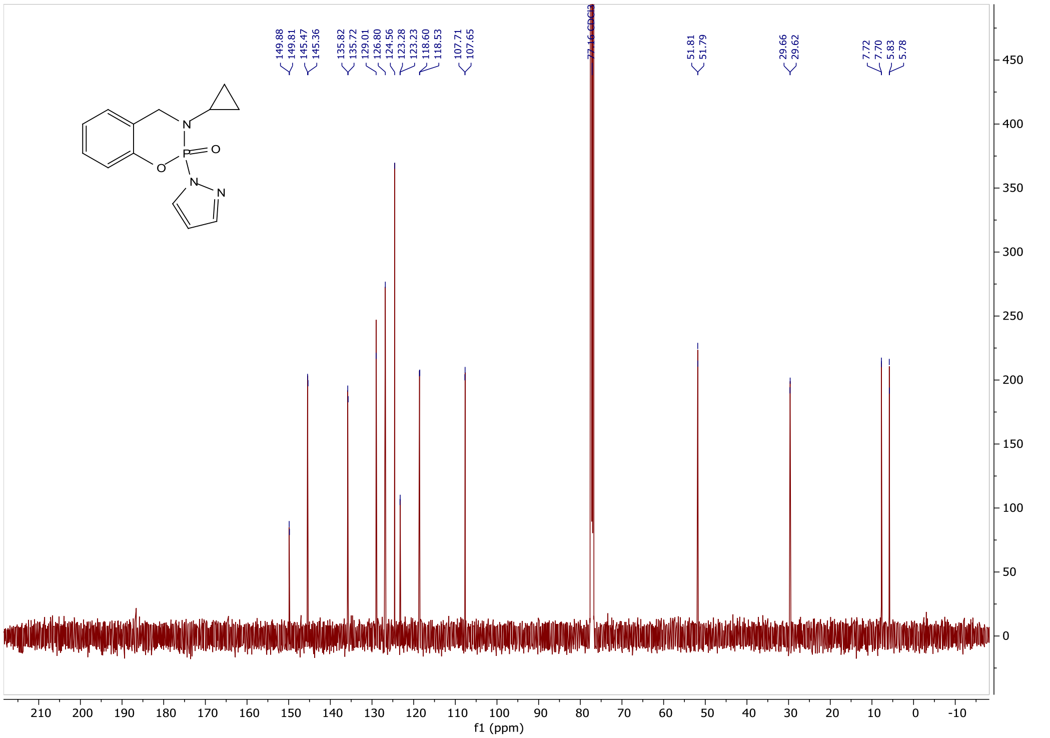

#### RJG-3130-7 $^{1}$H NMR (400 MHz, cdcl$_3$)

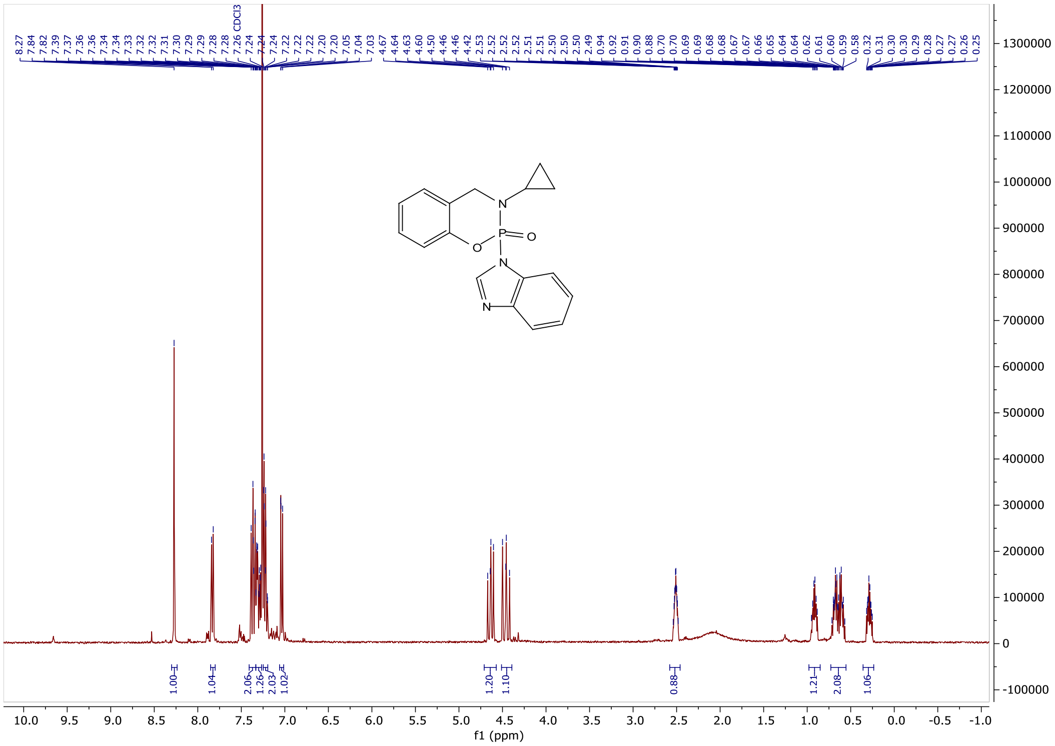

#### RJG-3130-7 $^{31}$P NMR (162 MHz, cdcl$_3$)

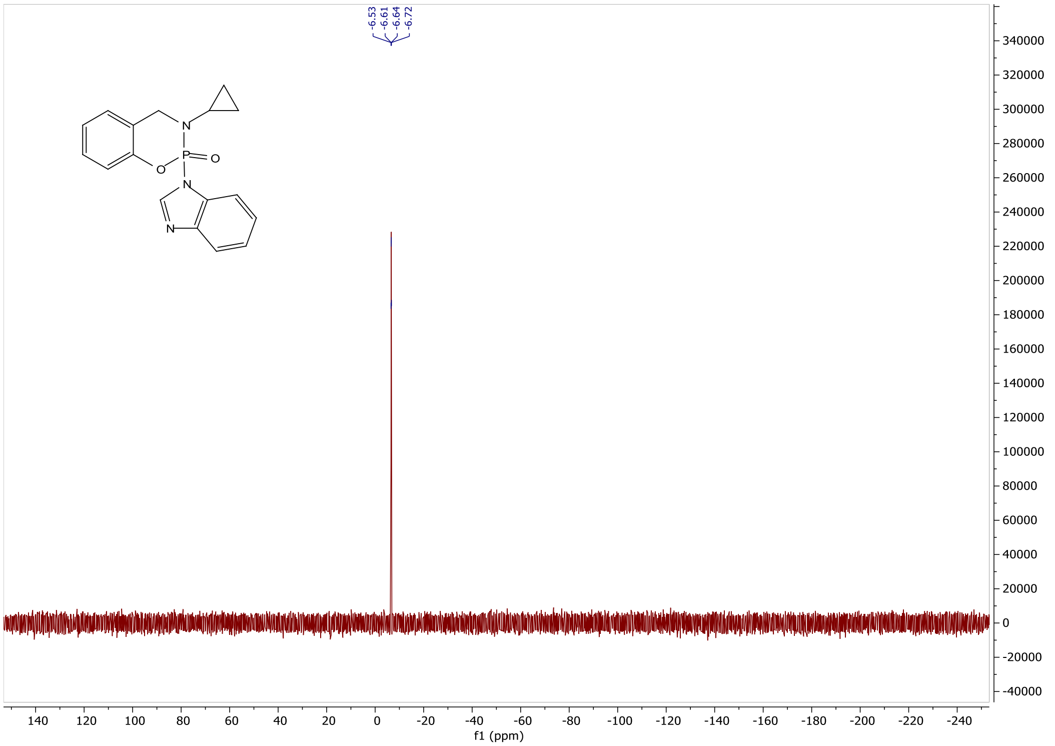

#### RJG-3130-7 $^{13}C$ NMR (126 MHz, cdcl$_3$)

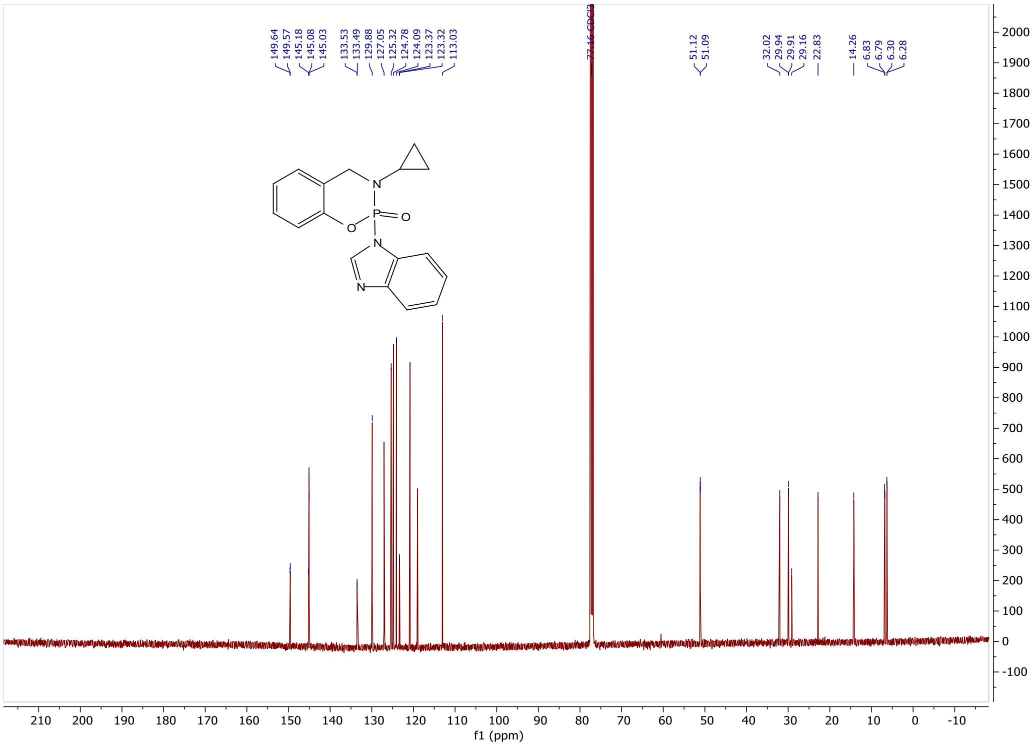

#### RJG-3130-8 $^{1}H$ NMR (500 MHz, cdcl$_3$)

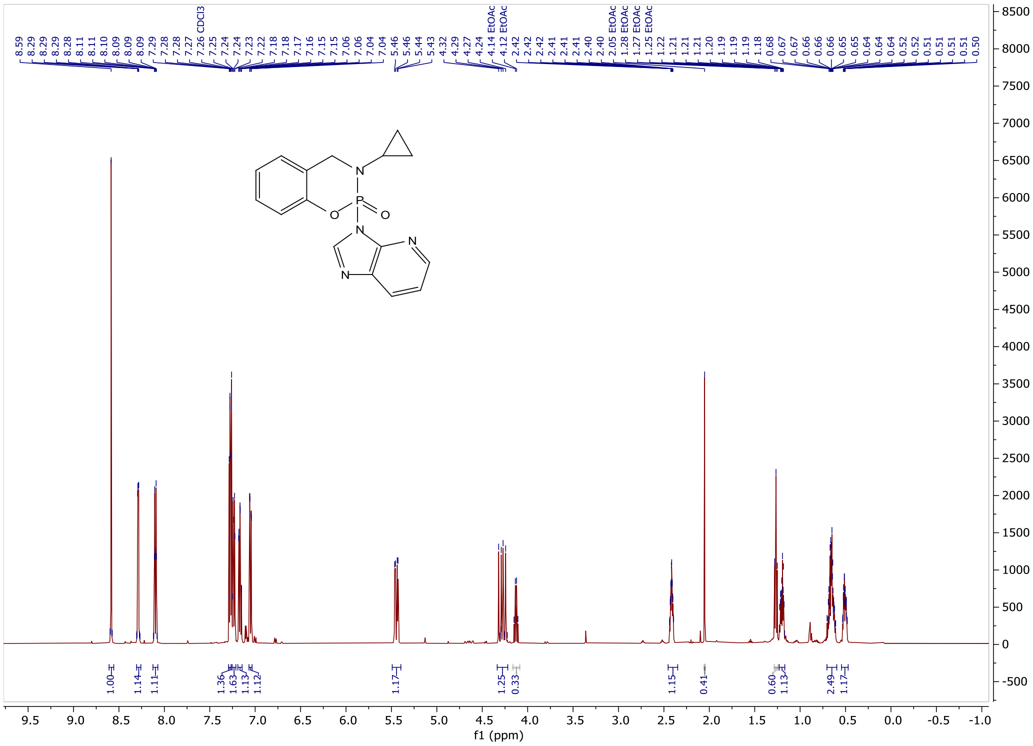

#### RJG-3130-8 $^{31}P$ NMR (162 MHz, $cdcl_3$)

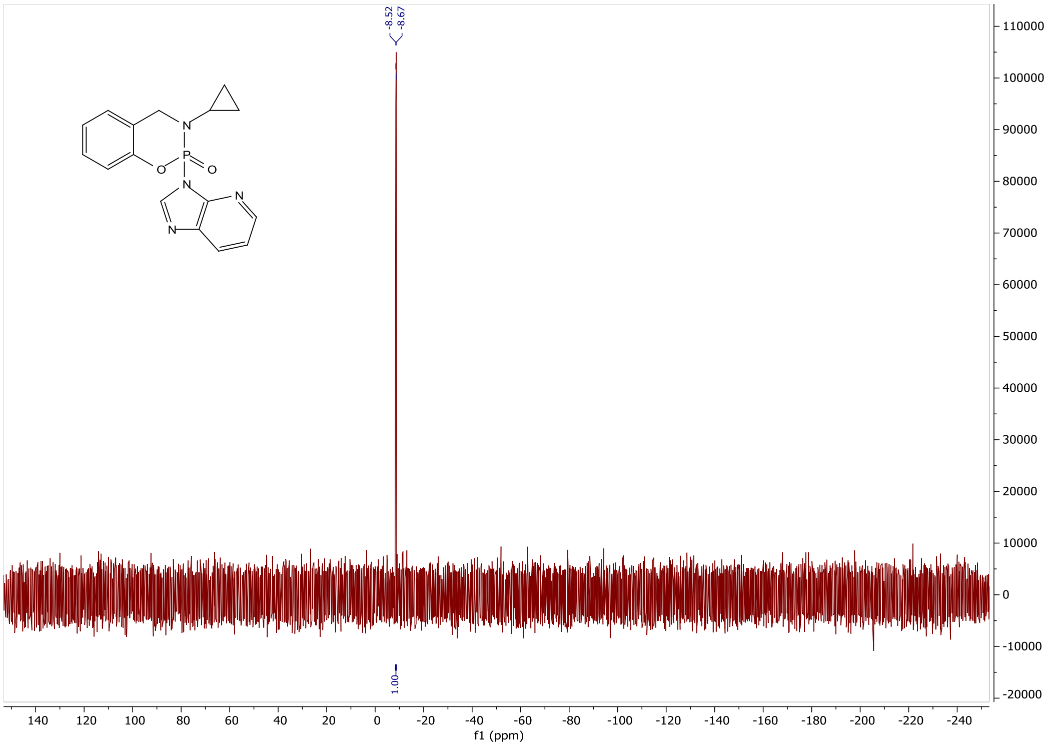

#### RJG-3130-8 $^{13}C$ NMR (126 MHz, $cdcl_3$)

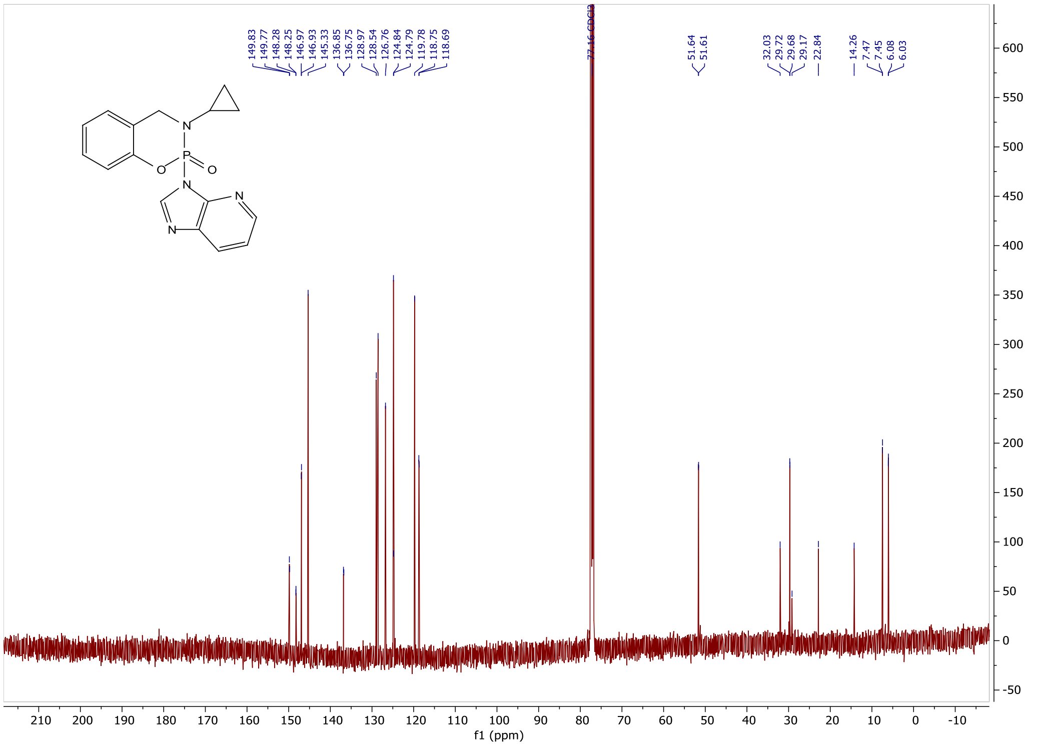

### RJG-3130-9 $^{1}$H NMR (400 MHz, cdcl$_3$)

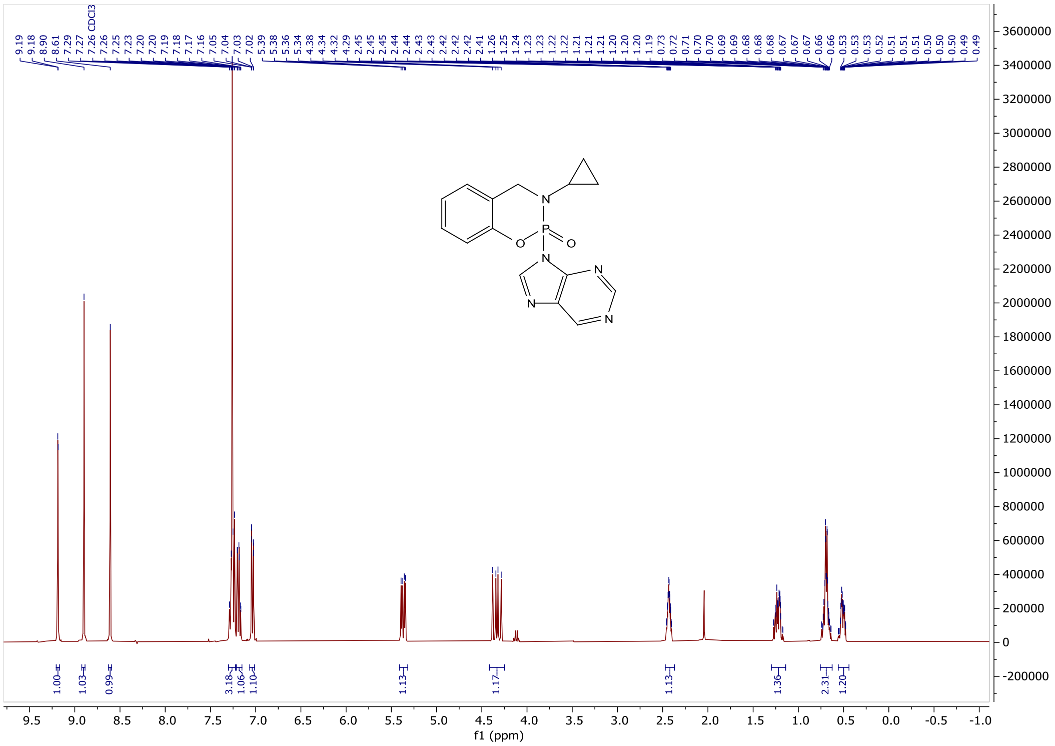

### RJG-3130-9 $^{31}$P NMR (162 MHz, cdcl$_3$)

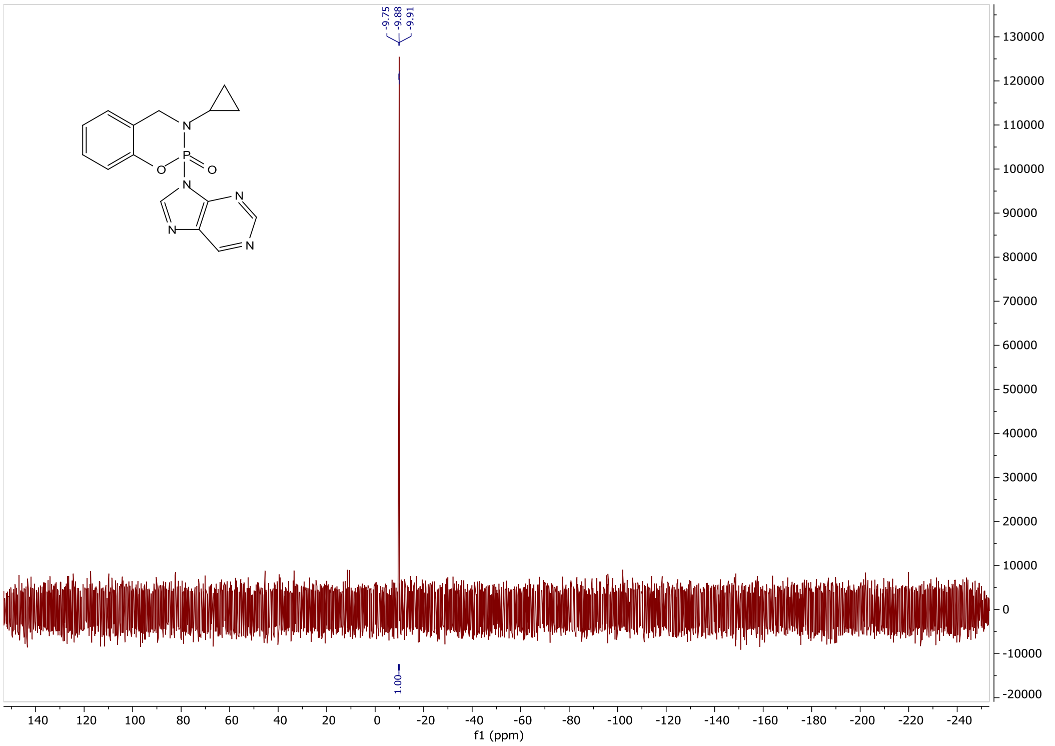

RJG-3130-9 $^{13}C$ NMR (126 MHz, cdcl$_3$)

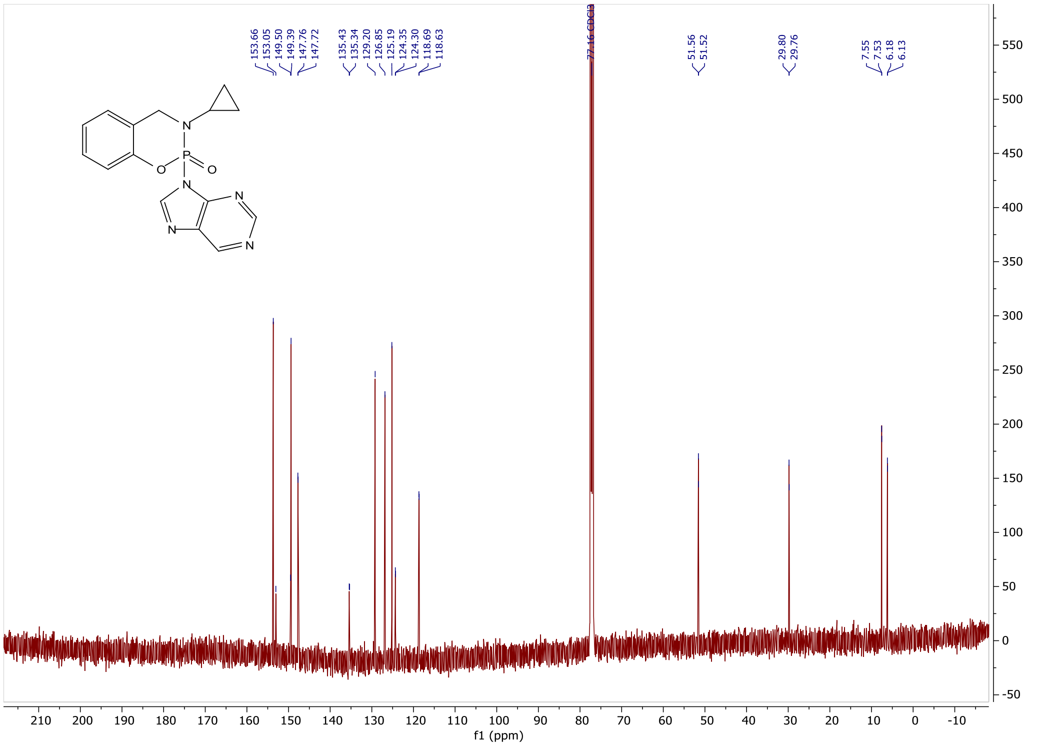

**X-Ray Crystal Structure Experimental and Data Tables**

**X-ray Experimental general**

Data were collected using an XtaLAB Synergy, Single source at home/near, HyPix diffractometer and on a SCXMini Single source diffractometer operating at $T$ = 100 K.
The diffraction pattern was indexed and the total number of runs and images was based on the strategy calculation from the program CrysAlisPro (Rigaku, V1.171.44.114a, 2025) The maximum resolution that was achieved was $\Theta = 76.484^{\circ}$ (0.79 Å) for (S)-(+)RJG-2273 and $\Theta$ = 64.04 (0.67 Å) for (S)-(+)RJG-2259A.
The diffraction pattern was indexed and the total number of runs and images was based on the strategy calculation from the program CrysAlisPro (Rigaku, V1.171.44.114a, 2025) and the unit cell was refined using CrysAlisPro (Rigaku, V1.171.44.114a, 2025).
Data reduction, scaling and absorption corrections were performed using CrysAlisPro (Rigaku, V1.171.44.114a, 2025). The final completeness is 100.00 % out to $76.484^{\circ}$ in $\Theta$. A multi-scan absorption correction was performed using CrysAlisPro 1.171.44.114a (Rigaku Oxford Diffraction, 2025) using spherical harmonics, implemented in SCALE3 ABSPACK scaling algorithm.

**(*S*)-(+)RJG-2273**

Single colourless plate-shaped crystals of rjg_so2_006 were obtained by recrystallisation from chloroform. A suitable crystal 0.32×0.12×0.05 $mm^3$ was selected and mounted on a suitable support on an XtaLAB Synergy. The structure was solved with the ShelXT[10,11] structure solution program using the Intrinsic Phasing solution method and by using Olex2[9] as the graphical interface. The model was refined with version 2019/3 of ShelXL 2019/3[10,11] using Least Squares minimisation.

| | | |
|---|---|---|
| Diffractometer | XtLAB Mini (JP) | |
| Identification code | rjg_so2_006 | |
| Empirical formula | $C_{10}H_9FNO_2P$ | |
| Formula weight | 225.15 | |
| Temperature/K | 100 | |
| Crystal system | monoclinic | |
| Space group | $P2_1$ | |
| Volume | a/Å=6.96520(10) | α/°=90 |
| | b/Å=5.22670(10) | β/°=96.391(2) |
| | c/Å=13.9445(3) | γ/°=90 |
| Volume/Å$^3$ | 504.495(16) | |
| Z | 2 | |
| ρcalcg/cm$^3$ | 1.482 | |
| μ/mm$^{-1}$ | 2.404 | |
| F(000) | 232.0 | |
| Crystal size/mm$^3$ | 0.32 × 0.12 × 0.05 | |
| Radiation | Cu Kα (λ = 1.54184) | |

| | |
|---|---|
| 2Θ range for data collection/° | 6.378 to 152.968 |
| Index ranges | $-7 \leq h \leq 8$, $-6 \leq k \leq 6$, $-17 \leq l \leq 17$ |
| Reflections collected | 9309 |
| Independent reflections | 2025 [Rint = 0.0567, Rsigma = 0.0318] |
| Data/restraints/parameters | 2025/1/136 |
| Goodness-of-fit on $F^2$ | 1.061 |
| Final R indexes [I>=2σ (I)] | $R_1$ = 0.0402, $wR_2$ = 0.1091 |
| Final R indexes [all data] | $R_1$ = 0.0405, $wR_2$ = 0.1095 |
| Largest diff. peak/hole / e $Å^{-3}$ | 0.47/-0.40 |
| Flack parameter | -0.01(2) |

**Table S1**: Cystal data and structure refinement for **(*S*)-(+)RJG-2273**

**(*S*)-(+)RJG-2259A**

Single crystals of RGJ_SO2_008 were obtained from crystallization from chloroform. A suitable crystal 0.11×0.24×0.35 $mm^3$ was selected and mounted on a SCXMini diffractometer equipped with Mo sealed tube source Mo Kα (λ = 0.71073) The crystal was kept at 100 K during data collection. Using Olex2[9], the structure was solved with the SHELXT[10] structure solution program using Intrinsic Phasing and refined with the XL[11] refinement package using Least Squares minimisation.

| | | |
|---|---|---|
| Diffractometer | XtLAB Mini (JP) | |
| Identification code | RJG-S02-008 | |
| Empirical formula | $C_{12}H_{11}N_4O_2P$ | |
| Formula weight | 274.22 | |
| Temperature/K | 100 | |
| Crystal system | Monoclinic | |
| Space group | $P2_1/c$ | |
| a/Å | 13.0826(10) | α/°=90 |
| b/Å | 6.8534(4) | β/°=114.370(9) |
| c/Å | 14.9481(12) | γ/°=90 |
| Volume/$Å^3$ | 1220.83(17) | |
| Z | 4 | |
| ρcalcg/$cm^3$ | 1.492 | |
| μ/$mm^{-1}$ | 0.23 | |
| F(000) | 568 | |
| Crystal size/$mm^3$ | 0.35 x 0.24 x 0.11 | |
| Radiation | Mo Kα (λ = 0.71073) | |
| 2Θ range for data collection/° | 3.418 to 64.04 | |
| Index ranges | $-19 \leq h \leq 19$, $-9 \leq k \leq 9$, $-19 \leq l \leq 20$ | |
| Reflections collected | 17671 | |
| Independent reflections | 4077 [$R_{int}$ = 0.0372, $R_{sigma}$ = 0.0336] | |
| Data/restraints/parameters | 4077/0/172 | |
| Goodness-of-fit on $F^2$ | 1.058 | |
| Final R indexes [I>=2σ (I)] | $R_1$ = 0.0423, $wR_2$ = 0.1084 | |

| | |
|---|---|
| Final R indexes [all data] | $R_1$ = 0.0553, $wR_2$ = 0.1185 |
| Largest diff. peak/hole / e Å$^{-3}$ | 0.86/-0.52 |

**Table S2**: Cystal data and structure refinement for **(*S*)-(+)RJG-2259A**

**(*S*)-(+)RJG-3130-9**

A small amount of the PH-UTXA-MCFTE-2024-001-S02-028-P2-0-1 sample was then spread over a holey carbon EM grid (Cu300 mesh *R*1.2/1.3, QUANTIFOIL) for MicroED test. Then the grid was placed on a single-tilt holder (ThermoFisher Scientific), which was further inserted into ThermoFisher Scientific Talos F200C electron microscope (operating voltage of 200 kV, wavelength of 0.0251 Å). Crystals were illuminated with a parallel electron beam in NanoProbe mode. After diffraction resolution screening, 25 crystals with high diffraction resolution (**Figure 2**) were selected for data collection. Sequential diffraction frames per crystal were collected by continuously rotating the sample stage at a goniometer rotation speed of 2° $s^{-1}$ using EPU-D software (ThermoFisher Scientific), and the diffraction patterns were recorded using a 4k × 4k Ceta-D CMOS camera (ThermoFisher Scientific). The total rotation angle of micro-crystal is 80-110° with an exposure time of 0.5 second for single diffraction pattern. High-quality MicroED data from 6 crystals were indexed and integrated in XDS, merged using XSCALE, and converted to SHELX format using XDSCONV. There are 12025 observed reflections and 2488 independent reflections in the merged data. Atomic scattering factor of electron was used for the calculation of theoretical structure factor during the structure determination. An initial structure model was obtained using intrinsic phasing algorithm in SHELXT and refined with SHELXL program. The atom type for all non-hydrogen atoms was determined by considering the electrostatic potential at each atom position, while taking into account the structural geometry, including bond length and assumed 2D structure. The identification of bond types in the structure was facilitated through an analysis of structural geometry, incorporating factors such as bond length and hydrogen atom positions. All atoms were refined anisotropically. All hydrogen atoms were placed using the riding model. After conducting structure solution using the aforementioned kinematical approximation, two possible absolute structures were identified, which have the centrosymmetric relationship with each other. The correct absolute structure was then determined through dynamical refinement, allowing for the absolute configuration determination of the small molecule. Five crystals with high-quality diffraction patterns were selected, each providing one independent MicroED dataset. PETS2 software was employed for data processing, and Jana2020 software was employed for subsequent dynamical refinement. The five datasets were individually utilized for refinement against the two potential absolute structures, resulting in five sets of independently refined outcomes.

| | | |
|---|---|---|
| **MicroED Instrument Details** | | |
| Polarized-light microscope | Axio Scope 5 POL (Ziess) | |
| CryoEM | Talos F200C (200 kV, ThermoFisher Scientific) | |
| Detector | Ceta-D (CMOS, ThermoFisher Scientific) | |
| Data collection software | EPU-D (Version 1.18.0.6930, ThermoFisher Scientific) | |
| CryoEM grid | Holey Carbon EM grid (Cu300 mesh *R*1.2/1.3, QUANTIFOIL) | |
| Cryogenic sample holder | Single-tilt holder (ThermoFisher Scientific) | |
| **Crystal Data and structure refinement** | | |
| Identification code | PH-UTXA-MCFTE-2024-001-S02-028-P2-0-1 | |
| Empirical formula | $C_{15}H_{14}N_5O_2P$ | |
| Formula weight | 327.28 | |
| Temperature/K | 293(2) | |
| Crystal system | monoclinic | |
| Space group | $P2_1$ (no.4) | |
| Volume | a/Å=5.74(1) | α/°=90 |
| | b/Å=14.50(4) | β/°=105.0(2) |
| | c/Å=9.59(5) | γ/°=90 |
| Volume/Å$^3$ | 771(5) | |
| Z | 2 | |
| ρcalcg/cm$^3$ | 1.410 | |
| μ/mm$^{-1}$ | 0.000 | |
| F(000) | 131.0 | |
| Crystal size/mm$^3$ | N.A. | |
| Radiation | Electron | |
| Wavelength/ Å | 0.0251 | |
| 2Θ range for data collection/° | 0.184 to 1.692 | |
| Resolution/ Å | 0.85 | |
| Reflections collected | 12025 | |
| Independent reflections | 2488 | |
| Completeness/% | 94.4 | |
| $R_{int}$ | 0.2218 | |
| $R_{sigma}$ | 0.1764 | |
| $I/\sigma_I$ | 5.7 | |
| Data/restraints/parameters | 2488/451/209 | |
| Goodness-of-fit on $F^2$ | 1.0754 | |
| Final R indexes [I>=2σ (I)] | $R_1$ = 0.1157, $wR_2$ = 0.2963 | |

| | |
|---|---|
| Final R indexes [all data] | $R_1$ = 0.1688, $wR_2$ = 0.3316 |

**Table S3**: Cystal data and structure refinement for **(*S*)-(+)RJG-3130-9**

#### References

(1) Franks, C. E.; Campbell, S. T.; Purow, B. W.; et al. The Ligand Binding Landscape of Diacylglycerol Kinases. *Cell Chem. Biol.* **2017**, *24* (7), 870-880.e875.
(2) Hahm, H. S.; Toroitich, E. K.; Borne, A. L.; et al. Global targeting of functional tyrosines using sulfur-triazole exchange chemistry. *Nat. Chem. Biol.* **2020**, *16* (2), 150-159.
(3) Hammond, P. S.; Forster, J. S. A microassay-based procedure for measuring low levels of toxic organophosphorus compounds through acetylcholinesterase inhibition. *Anal. Biochem.* **1989**, *180* (2), 380-383.
(4) Ellman, G. L.; Courtney, K. D.; Andres, V.; Featherstone, R. M. A new and rapid colorimetric determination of acetylcholinesterase activity. *Biochem. Pharmacol.* **1961**, *7* (2), 88-95.
(5) Adibekian, A.; Martin, B. R.; Wang, C.; et al. Click-generated triazole ureas as ultrapotent in vivo–active serine hydrolase inhibitors. *Nat. Chem. Biol.* **2011**, *7* (7), 469-478.
(6) Kim, Y. B.; Kopcho, L. M.; Kirby, M. S.; et al. Mechanism of Gly-Pro-pNA cleavage catalyzed by dipeptidyl peptidase-IV and its inhibition by saxagliptin (BMS-477118). *Arch. Biochem. Biophys.* **2006**, *445* (1), 9-18.
(7) Sun, S.; Smedley, C. J.; Homer, J. A.; et al. Phosphorus(V) Fluoride Exchange (PFEx): Multidimensional Click Chemistry from Phosphorus(V) Connective Hubs. *ChemRxiv* **2022**.
(8) Liu, Y.; Patricelli, M. P.; Cravatt, B. F. Activity-based protein profiling: The serine hydrolases. *Proc. Natl. Acad. Sci.* **1999**, *96* (26), 14694-14699.
(9) Dolomanov, O.V., Bourhis, L.J., Gildea, R.J, Howard, J.A.K. & Puschmann, H. (2009), J. Appl. Cryst. 42, 339-341.
(10) Sheldrick, G.M. (2015). Acta Cryst. A71, 3-8.
(11) Sheldrick, G.M. (2015). Acta Cryst. C71, 3-8.
(12) Agilent (2014). *CrysAlis PRO*. Agilent Technologies Ltd, Yarnton, Oxfordshire, England.
